# Linkage Disequilibrium and Heterozygosity Modulate the Genetic Architecture of Human Complex Phenotypes

**DOI:** 10.1101/705285

**Authors:** Dominic Holland, Oleksandr Frei, Rahul Desikan, Chun-Chieh Fan, Alexey A. Shadrin, Olav B. Smeland, Ole A. Andreassen, Anders M. Dale

**Affiliations:** Center for Multimodal Imaging and Genetics, University of California at San Diego, La Jolla, CA 92037, USA; Department of Neurosciences, University of California, San Diego, La Jolla, CA 92093, USA; Department of Radiology, University of California, San Francisco, San Francisco, CA 94158, USA; NORMENT, KG Jebsen Centre for Psychosis Research, Institute of Clinical Medicine, University of Oslo 0424 Oslo, Norway; Department of Cognitive Sciences, University of California at San Diego, La Jolla, CA 92093, USA; Department of Radiology, University of California, San Diego, La Jolla, CA 92093, USA; Division of Mental Health and Addiction, Oslo University Hospital, 0407 Oslo, Norway; Department of Psychiatry, University of California, San Diego, La Jolla, CA 92093, USA

**Keywords:** Heritability, Polygenicity, Effect size, Linkage disequilibrium, Minor allele frequency, Natural selection

## Abstract

We propose an extended Gaussian mixture model for the distribution of causal effects of common single nucleotide polymorphisms (SNPs) for human complex phenotypes that depends on linkage disequilibrium (LD) and heterozygosity (H), while also allowing for independent components for small and large effects. Using a precise methodology showing how genome-wide association studies (GWAS) summary statistics (z-scores) arise through LD with underlying causal SNPs, we applied the model to GWAS of multiple human phenotypes. Our findings indicated that causal effects are distributed with dependence on total LD and H, whereby SNPs with lower total LD and H are more likely to be causal with larger effects; this dependence is consistent with models of the influence of negative pressure from natural selection. Compared with the basic Gaussian mixture model it is built on, the extended model – primarily through quantification of selection pressure – reproduces with greater accuracy the empirical distributions of z-scores, thus providing better estimates of genetic quantities, such as polygenicity and heritability, that arise from the distribution of causal effects.

## INTRODUCTION

There is currently great interest in the distribution of causal effects among trait-associated single nucleotide polymorphisms (SNPs), and recent analyses of genome-wide association studies (GWAS) have begun uncovering deeper layers of complexity in the genetic architecture of complex human traits and disorders (Gazal et al., 2017; Zeng et al., 2018; Zhang et al., 2018; Frei et al., 2019; Holland et al., 2020). This research is facilitated by using new analytic approaches to interrogate structural features in the genome and their relationship to phenotypic expression. Some of these analyses take into account the fact that different classes of SNPs have different characteristics and play a multitude of roles (Schork et al., 2013; Finucane et al., 2015; Shadrin et al., 2019). Along with different causal roles for SNPs, which in itself would suggest differences in distributions of effect-sizes for different categories of causal SNPs, the effects of minor allele frequency (MAF) of the causal SNPs and their total correlation with neighboring SNPs are providing new insights into the action of selection on the genetic architecture of complex traits (Gazal et al., 2017; Wray et al., 2018; Zhang et al., 2018).

Any given mutation is likely to be neutral or deleterious to fitness (Fay et al., 2001). Natural selection partly determines how the prevalence of a variant develops over time in a population, and evidence for its action can be found in the relationship between effect size and MAF in complex traits and common diseases (Zeng et al., 2018). Negative selection acts predominantly to keep variants deleterious to fitness at low frequency, or ultimately remove them. The larger the effect of a deleterious variant the more efficient negative selection will be, suggesting that the lower the MAF the larger the effect size – and this is expected under evolutionary models (Pritchard and Cox, 2002; Eyre-Walker, 2010), and consistent with empirical findings (Park et al., 2011) and recent analyses based on genome-wide associations (Zeng et al., 2018; Schoech et al., 2019; O’Connor et al., 2019).

The effect of linkage disequilibrium (LD) has also been studied, suggesting that SNPs with low “levels of LD” (LLD) explain more heritability, which is again consistent with the action of negative selection (Gazal et al., 2017). One unexplored issue is how the prior probability of a SNP being causal depends on its LD score (or related measures). Due to the complexity of genetic forces acting on alleles, it is not clear what form any such dependency might take. However, explicitly modeling any such role promises to yield a closer match between empirical distributions of GWAS summary statistics and model predictions, and thereby can provide more accurate estimates of quantities of interest like polygenicity and heritability.

In previous work (Holland et al., 2020), building on earlier reports of others (e.g., George and McCulloch (1993); Erbe et al. (2012); Zhou et al. (2013)), we presented a basic Gaussian Gaussian mixture model to describe the distribution of underlying causal SNP effects (the per unit allelic effect, *β*, estimated from simple linear regression). Due to extensive and complex patterns of LD among SNPs, many non-causal SNPs will exhibit strong association with phenotypes, resulting in a far more complicated distribution for the summary z-scores. The basic model for the distribution of the causal *β*’s is a mixture of non-null and null normal distributions, the latter (denoted 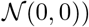 being just a delta function (or point mass at zero):

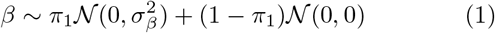

where *π*_1_ is the polygenicity, i.e., the proportion of SNPs that are causal (equivalently, the prior probability that any particular SNP is causal), and 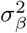 is the discoverability (the expectation of the square of the effect size), which was taken to be a constant across all causal SNPs. The distribution of z-scores arising from this shows strong heterozygosity (H=MAF×(1-MAF)) and total LD (TLD) dependence; TLD, defined in the subsection 1.3, is similar to LD score (Bulik-Sullivan et al., 2015) but takes into account more neighboring SNPs.

The recent work by others focusing on selection pressure using a single causal Gaussian (Zeng et al., 2018; Schoech et al., 2019) indicated that it is important to take SNP heterozygosity into account as a component in discoverability (initially explored by examining four fixed relationships in (Speed et al., 2012); see also (Lee et al., 2013)). It was also recently shown that an additional Gaussian distribution for the *β*’s might be appropriate if large and small effects are distributed differently (Zhang et al., 2018). These approaches, however, were not combined – a single causal Gaussian incorporating heterozygosity, versus two causal Gaussians with no heterozygosity dependence. An extra complexity is that measures related to total LD, such as LLD (Gazal et al., 2017), can be expected to play an important role in the distribution of causal effects. It is unclear how all these factors impact each other. A final and very important matter is that many analyses in the literature by other groups were conducted in the “infinitesimal” model framework (Finucane et al., 2015; Bulik-Sullivan et al., 2015; Gazal et al., 2017; Schoech et al., 2019; Speed and Balding, 2019), where all SNPs are causal, though there is compelling evidence that only a very small fraction of SNPs are in fact causal for any given phenotype (Zhu and Stephens, 2017; Zeng et al., 2018; Zhang et al., 2018; Frei et al., 2019; Shadrin et al., 2019; Holland et al., 2020).

In the current work, we sought to extend our earlier work to incorporate multiple Gaussians, while taking into account TLD as a factor in polygenicity and selection effects reflected in heterozygosity, in modeling the distribution of causal *β*’s. With a wide range of model parameter values across real phenotypes, the specificity of the individual parameters for a given phenotype make them more narrowly defining of the distribution of summary statistics for that phenotype.

## 1. Materials and Methods

### 1.1. The model: an extension of prior work

The methodology calls for using an extensive reference panel that likely involves all possible causal SNPs with MAF greater than some threshold (e.g., 1%), and regard the z-scores for typed or imputed SNPs – a subset of the reference SNPs – as arising, directly or through LD, from the underlying causal SNPs in the reference panel.

Since the single causal Gaussian, Eq. 1, has provided an appropriate starting point for many phenotypes, it is reasonable to build from it. With additional terms included, if it turns out that this original term is not needed, the fitting procedure, if implemented correctly, should eliminate it. Also, anticipating extra terms in the distribution of causal *β*’s, we introduce a slight change in labeling the Gaussian variance 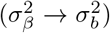, and write the distributions for the causal component only – it being understood that the full distribution will include the last term on the right side of Eq. 1 for the prior probability of being null.

Given that for some phenotypes there is strong evidence that rarer SNPs have larger effects, we next include a term that reflects this: a Gaussian whose variance is proportional to *H^S^*, where *H* is the SNP’s heterozygosity and *S* is a new parameter which, if negative, will reflect the noted behavior (Zeng et al., 2018). With the addition of the new term, the total prior probability for the SNP to be causal is still given by *π*_1_. Thus, extending Eq. 1, we get:

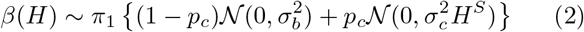

where *p*_*c*_ (0 ⩽ *p*_*c*_ ⩽ 1) is the prior probability that the SNP’s causal component comes from the “c” Gaussian (with variance 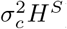), and *p*_*b*_ ≡ 1−*p*_*c*_ is the prior probability that the SNP’s causal component comes from the “b” Gaussian (with variance 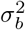). This extension introduces an extra three parameters *p*_*c*_, *σ_c_*, and *S*, assumed for the moment to be the same for all SNPs. Ignoring inflation and implementation details like choice of reference panel and parameter estimation scheme, setting *p*_*c*_ ≡ 1 recovers the model distribution assumed in (Zeng et al., 2018); additionally setting *π*_1_ ≡ 1 recovers the primary model distribution assumed in (Schoech et al., 2019); and additionally setting *S* ≡ −1 recovers the model assumed in (Bulik-Sullivan et al., 2015), while instead setting *S* ≡ −0.25 partially recovers the “recommended” LD-adjusted kinships (LDAK) model distribution in (Speed et al., 2017; Speed and Balding, 2019). For a further discussion of LDAK and variants of the LD Score regression model, including “stratified LD fourth moments regression” (S-LD4M) (O’Connor et al., 2019) which introduces an effective number of causal SNPs (*M_e_*), see 3.

Within the infinitesimal models (Speed et al., 2012; Finucane et al., 2015; Speed et al., 2017; Gazal et al., 2017; Schoech et al., 2019; Speed and Balding, 2019), it is not clear the degree to which explicit LD-dependence in the variance of the causal effect size merely takes into account the effect on z-scores due to LD with causal SNPs, and how much it models any true effect of LD on underlying causal effect size. Also, such models preclude examining if the TLD of a variant has any bearing on whether the variant is causal. In contrast is the “BayesS” model (Zeng et al., 2018), a causal mixture model (i.e., not infinitesimal), using individual genotype data and a reference panel of ∼484k non-imputed SNPs, that examines the effects of heterozygosity on effect size. The model we present here is in some respects an extension of that, but based on summary statistics, adding TLD dependence and an additional Gaussian, and we fit the model from a reference panel of 11 million SNPs using the exact procedure – convolution – to relate posited underlying distributions of causal effects to empirical distributions of z-scores.

Along with heterozygosity, SNP effect size might independently depend on TLD (for which we use the variable *L* in equations below), and in principle this could be explored in a manner similar to how heterozygosity is incorporated in the “c” causal effect Gaussian variance (e.g., with an extra factor *L^T^*, say, scaling 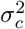, where *T* would be a new parameter). However, TLD and heterozygosity are often related (given TLD, the expected heterozygosity shows a distinct well-defined pattern for SNPs with TLD<200, i.e., about 80% of SNPs – see Supplemental Material, Figure S6), and independent contributions might be difficult to disentangle. Instead, here we explore a separate mathematical role for TLD.

There is no obvious *a priori* reason why the probability (in a Bayesian sense) of a SNP’s being causally associated with any particular trait should be independent of the SNP’s TLD. Indeed, with the complex interaction of multiple genetic forces such as mutation, genetic drift, and selection, the net relationship between TLD – through mechanisms like background selection – and causal association with a particular phenotype is not clear. The results of Gazal et al. (2017), however, indicate that SNPs with low LLD have significantly larger per-SNP heritability. Note that in Eq. 2, for 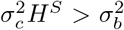 the “c” Gaussian will describe larger effect. In this case, the LLD dependence suggests modulating the “c” Gaussian such that *p*_*c*_ is larger for smaller TLD. Whatever the relationship, however, the more accurately it is incorporated in a model for the distribution of effect sizes should lead to more accurate reproduction of the distribution of empirical summary statistics and estimation of quantities of interest like polygenicity, heritability, and selection effects.

As heterozygosity decreases, SNPs will continue to have a range of total LD (see Supplemental Material, Figure S7 for the number density of SNPs with respect to heterozygosity and TLD). We explore here the possibility that the prior probability of being causal with large effect decreases with TLD. If the “c” Gaussian is capturing larger effects from rarer SNPs, reflecting selection pressure, we inquire if the prior probability for a causal SNP’s contribution from the “c” Gaussian is TLD-mediated. Specifically, instead of treating *p*_*c*_ as a constant, we explore the possibility that it is larger for SNPs with lower TLD; in the event that this probability is in fact constant, or increasing, with respect to TLD, we would at least not expect to find it decreasing. This can be accomplished by means of a generalized sigmoidal function that will have a maximum at very low TLD, might maintain that maximum for all SNPs (equivalently, *p*_*c*_ is a constant), or decrease in amplitude slowly or rapidly, possibly to 0, for SNPs with higher TLD. With the variable *L* denoting the TLD of a SNP, such a function of TLD can be characterized by three parameters: its amplitude (at *L* = 1), the TLD at the midpoint of the sigmoidal transition, and the width of the sigmoidal transition (over a wide or narrow range of TLD). We use the following general form with three parameters, *y*_*max*_, *x*_*mid*_, and *x*_*width*_:

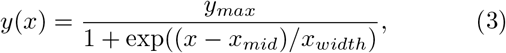

defined in the range −∞ < *x* < ∞, for which *y*(*x*) is monotonically decreasing and bounded 0 < *y*(*x*) < *y_max_*, with 0 ⩽ *y*_*max*_ ⩽ 1; −∞ < *x_mid_* < ∞ locates the mid point of the overall sigmoidal transition (*y*(*x*_*mid*_) = *y_max_/*2), and 0 < *x_width_* < ∞ controls its width (*y*(*x*) smoothly changing from a Heaviside step function at *x*_*mid*_ as *x_width_* → 0 to a constant function as *x_width_* → ∞). Examples are shown in Figure 1 (scaled by *π*_1_, giving the physically interesting total prior probability of a SNP being causal with respect to the selection “c” Gaussian, as a function of the SNP’s TLD); parameter values are in 3 Table G1. Mathematically, the curve can continue into the “negative TLD” range, revealing a familiar full sigmoidal shape; since we are interested in the range 1 ⩽ *x* ⩽ max(TLD), below we report the actual mid point (denoted *m_c_*) and width (*w_c_*, defined below) of the transition that occurs in this range.

Then

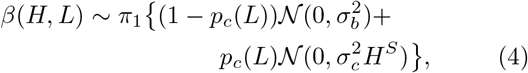

where *p*_*c*_(*L*) is the sigmoidal function (0 ⩽ *p*_*c*_(*L*) ⩽ 1 for all *L*) given by *y*(*x*) in Eq. 3 for *L* = *x* ⩾ 1, which numerically can be found by fitting for its three characteristic parameters.

**Figure 1:**
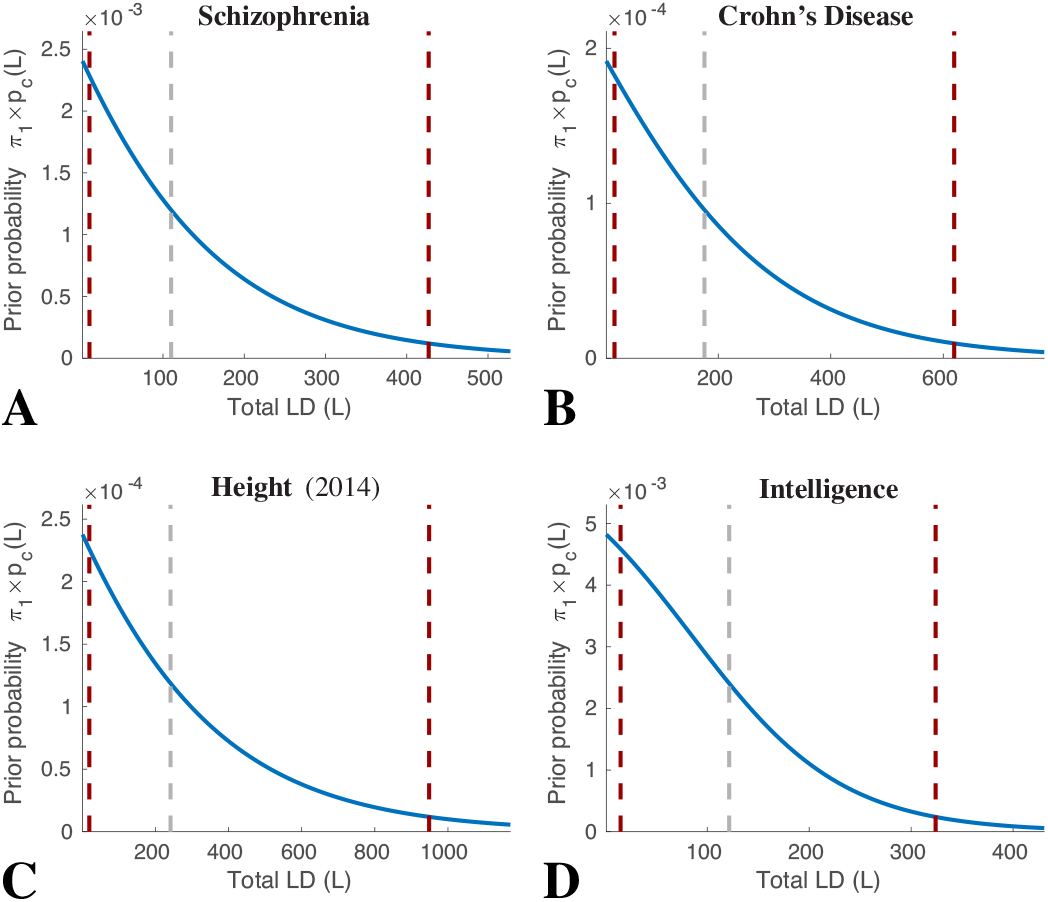
Examples of *π*_1_× prior probability functions *p*_*c*_(*L*) used in Eqs. 4 and 5, where *L* is reference SNP total LD (see Eq. 3 for the general expression, and 3 Table G1 for parameter values). These functions can be summarized by three quantities: the maximum value, *p*_*c*1_, which occurs at *L* = 1; the total LD value, *L* = *m_c_*, where *p*_*c*_(*m_c_*) = *p*_*c*1_/2, give by the gray dashed lines in the figure; and the total LD width of the transition region, *wc*, defined as the distance between where *p*_*c*_(*L*) falls to 95% and 5% of *p*_*c*1_ given by the flanking red dashed lines in the figure. Numerical values of *p*_*c*1_, *m_c_*, and *w_c_* are given in Table 1 and Figures 2 and 3. *p_d_*(*L*) is similar. Plots of *p*_*c*_(*L*) and *p_d_*(*L*), where relevant, for all phenotypes are shown in Supplemental Material, Figures S3-S5.

**Table 1:** Model parameters for phenotypes, case-control (upper section) and quantitative (lower section). *π*_1_ is the overall proportion of the 11 million SNPs from the reference panel that are estimated to be causal. *p*_*c*_(*L* ⩾ 1) is the prior probability multiplying the “c” Gaussian, which has variance 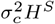, where *H* is the reference SNP heterozygosity. Note that *p*_*c*_(*L*) is just a sigmoidal curve, and can be characterized quite generally by three parameters: the value *p*_*c*1_ ≡ *p*_*c*_(1) at *L* = 1; the total LD value *L* = *m_c_* at the mid point of the transition, i.e., *p*_*c*_(*m_c_*) = *p*_*c*1_/2 (see the middle gray dashed lines in Figure 1, which shows examples of the function *p*_*c*_(*L*)); and the width *w_c_* of the transition, defined as the distance (in *L*) between where the curve falls to 95% and 5% of *p*_*c*1_ (distance between the flanking red dashed lines in Figure 1). Note that for AD Chr19, AD NoC19, and ALS Chr9, *π*_1_ is the fraction of reference SNPs on chromosome 19, on the autosome excluding chromosome 19, and on chromosome 9, respectively. Examples of *H^S^* multiplying 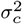 are shown in Supplemental Material, Figure S9. Model selection was performed using Bayesian information criterion (BIC). Except for Height 2018 which is shown here for model C for direct comparison with the 2010 and 2014 GWASs, and whose model D fit is shown in Supplemental Material, Figure S35, missing model D parameters indicate that those parameters could not reliably be estimated. For Crohn’s disease, ulcerative colitis, and total cholesterol *p_d_*(*L*) = *p*_*d*1_ for all L. Estimated BIC values for three models (B, C, and D) are shown in 3 Table D1: the 3-parameter model B with only the “b” Gaussian (*π*_1_, *σ_b_, σ*_0_); the 8-parameter model C with both the “b” with “c” Gaussians (Eq. 4); and the 12-parameter model D with “b”, “c” and “d” Gaussians (Eq. 5). 95% confidence intervals are in 3 Tables E1–E3.

| Phenotype | $\pi_1$ | $\sigma_b^2$ | $\sigma_c^2$ | $S$ | $p_{c1}$ | $m_c$ | $w_c$ | $\sigma_d^2$ | $p_{d1}$ | $m_d$ | $w_d$ | $\sigma_0^2$ |
| --- | --- | --- | --- | --- | --- | --- | --- | --- | --- | --- | --- | --- |
| SCZ 2014 | 5.28e-2 | 1.4e-6 | 3.6e-5 | -0.52 | 0.07 | 87 | 352 | 1.4e-4 | 5.2e-3 | 519 | 7 | 1.07 |
| BIP | 5.91e-2 | 1.2e-6 | 4.5e-5 | -0.40 | 6.7e-2 | 102 | 414 | — | — | — | — | 1.01 |
| CD | 9.55e-4 | 5.0e-5 | 5.5e-4 | -0.64 | 0.20 | 176 | 604 | 7.6e-2 | 1.0e-4 | — | — | 1.14 |
| UC | 1.16e-3 | 3.6e-5 | 4.0e-4 | -0.67 | 0.16 | 173 | 627 | 8.0e-2 | 1.0e-4 | — | — | 1.12 |
| CAD | 1.88e-3 | 1.1e-5 | 9.2e-5 | -0.51 | 6.9e-2 | 171 | 683 | 5.3e-3 | 3.6e-4 | 102 | 7 | 0.92 |
| AD Chr19 | 4.34e-4 | 1.0e-4 | 6.1e-3 | -0.57 | 0.73 | 35 | 89 | — | — | — | — | 1.09 |
| AD NoC19 | 1.05e-3 | 1.8e-5 | 2.6e-4 | -0.52 | 2.9e-2 | 264 | 6 | — | — | — | — | 1.04 |
| ALS Chr9 | 1.12e-2 | 7.3e-6 | 3.9e-3 | -0.01 | 1.8e-3 | 106 | 6 | — | — | — | — | 0.99 |
| Edu | 1.43e-2 | 1.7e-6 | 7.8e-6 | -0.44 | 0.45 | 111 | 339 | 8.5e-5 | 6.3e-3 | 441 | 7 | 0.94 |
| IQ 2018 | 1.27e-2 | 7.5e-7 | 6.2e-6 | -0.51 | 0.38 | 122 | 309 | 3.6e-5 | 7.8e-2 | 561 | 6 | 1.17 |
| Height 2010 | 1.02e-3 | 4.1e-5 | 2.0e-4 | -0.44 | 0.20 | 322 | 1243 | — | — | — | — | 0.90 |
| Height 2014 | 1.15e-3 | 3.7e-5 | 1.6e-4 | -0.46 | 0.21 | 242 | 929 | — | — | — | — | 1.57 |
| Height 2018 | 2.50e-3 | 8.7e-6 | 8.9e-5 | -0.43 | 0.37 | 210 | 739 | — | — | — | — | 2.12 |
| HDL | 2.54e-3 | 1.1e-5 | 4.5e-4 | -0.79 | 1.4e-2 | 143 | 599 | 2.2e-2 | 1.1e-3 | 66 | 7 | 0.91 |
| LDL | 5.84e-3 | 3.3e-6 | 2.4e-4 | -0.52 | 8.8e-3 | 336 | 1417 | 7.3e-3 | 2.2e-4 | 346 | 6 | 0.92 |
| BMI GIANT 2015 | 1.54e-3 | 2.2e-5 | 4.5e-4 | 0.00 | 4.4e-3 | 288 | 12 | — | — | — | — | 0.85 |
| BMI 2018 | 2.34e-3 | 1.6e-5 | 3.0e-4 | 0.11 | 3.8e-3 | 268 | 8 | — | — | — | — | 1.72 |
| TC | 1.15e-3 | 1.7e-5 | 6.2e-4 | -0.97 | 2.1e-2 | 140 | 583 | 2.9e-4 | 3.4e-2 | — | — | 0.92 |

As a final possible extension, we add an extra term – a “d” Gaussian – to describe larger effects not well captured by the “b” and “c” Gaussians. This gives finally:

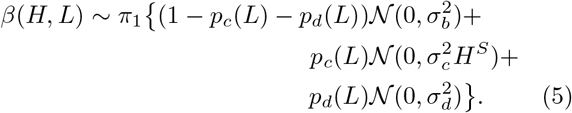

where 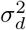 is a new parameter, *p_d_*(*L*) is another general sigmoid function (0 ⩽ *p_d_*(*L*) ⩽ 1 for all *L*) where now there is the added constraint 0 ⩽ *p*_*c*_(*L*) + *p_d_*(*L*) ⩽ 1, and the prior probability for the “b” Gaussian becomes *p*_*b*_(*L*) ≡1 − *p*_*c*_(*L*) − *p_d_*(*L*).

Depending on the phenotype and the GWAS sample size, it might not be feasible, or meaningful, to implement the full model. In particular, for low sample size and/or low discoverability, the “b” Gaussian is all that can be estimated, but in most cases both the “b”and “c” Gaussians can be estimated, and *β* will be well characterized by Eq. 4. We refer to the model given by Eq. 1 as model B; models C and D are given by Eqs. 4 and 5, respectively.

As we described in our previous work (Holland et al., 2020), a z-score is given by a sum of random variables, so the *a posteriori* pdf (given the SNP’s heterozygosity and LD structure, and the phenotype’s model parameters) for such a composite random variable is given by the convolution of the pdfs for the component random variables. This facilitates an essentially exact calculation of the z-score’s *a posteriori* pdf arising from the underlying model of causal effects as specified by Eq. 1, 4, or 5.

For our reference panel we used the 1000 Genomes phase 3 data set for 503 subjects/samples of European ancestry (Consortium et al., 2015, 2012; Sveinbjornsson et al., 2016). In ref. (Holland et al., 2020) we describe how we set up the general framework for analysis, including the use of the reference panel and dividing the reference SNPs into a sufficiently fine 10×10 heterozygosity×TLD grid to facilitate computations.

In our earlier work on the “b” model we gave an expression, denoted *G*(*k*), for the Fourier transform of the genetic contribution to a z-score, where *k* is the running Fourier parameter. The extra complexity in the “c” and “d” models here requires a modification only in this term, which we describe below in the 1.4 subsection. In addition to the parameters presented above, we also include an inflation parameter *σ*_0_: if *z_u_* denotes uninflated GWAS z-scores and *z* denotes actual GWAS z-scores, then *σ*_0_ is defined by *z* = *σ*_0_*z_u_* (Devlin and Roeder, 1999). The optimal model parameters for a particular phenotype are estimated by minimizing the negative of the log likelihood of the data (z-scores) as a function of the parameters. This is done with the “b” model as before, and then proceeding iteratively with the more complex models, continually re-estimating the new values of the earlier parameters that maximize the likelihood when a new parameter is introduced, with extensive single and multiple parameter searches (to avoid being trapped in local minima) until all parameters *simultaneously* are at a convex minimum. This involved many explicit and repeated coarse- and fine-grained linear (for each individual parameter) and grid (for multiple parameters simultaneously) searches as well as many Nelder-Mead multidimensional unconstrained non-linear minimizations. There is no artificial constraining on the parameters; for example, no prior assumption is made about the relative sizes of the causal *σ*’s, or the parameter values of the TLD-dependent prior probabilities (the full range of amplitudes, transition widths, and location of the mid-point of the transitions are searched).

The total SNP heritability is given by the sum of heritability contributions of each SNP in the reference panel from each of the relevant Gaussians. In the Bayesian approach, we do not know the value of the causal effect of any particular SNP, but we assume it comes from a distribution, *β*(*H, L*), which characterize our ignorance of it. For a specific Gaussian (“b”, “c”, or “d”) in our model, the contribution of the SNP to heritability is given by the prior probability that the SNP is causal with respect to that Gaussian, times the expected value of the square of the effect size, E(*β*^2^), times *H*. For the “c” Gaussian, for example, the prior probability that the SNP is causal is *π*_1_*p*_*c*_(*L*), and E(*β*^2^) is just the variance, 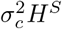. Thus, the contribution of this SNP to the overall heritability associated with the “c” Gaussian, 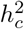, is 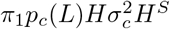. Below we report the sums over all such contributions. The number of causal SNPs associated with the “c” Gaussian, *n*_*c*_, is given by summing *π*_1_*p*_*c*_(*L*) for each reference panel SNP, and similarly for the other Gaussians. All heritabilities and discoverabilities are, as before, corrected with respect to the inflation parameter, i.e., divided by 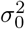.

All code used in the analyses, including simulations, is publicly available on GitHub (Holland, 2019b,a).

### 1.2. Data Preparation

We analyzed summary statistics for fourteen phenotypes-genotypes (in what follows, where sample sizes varied by SNP, we quote the median value): (1) bipolar disorder (*N_cases_* = 20,352, *N_controls_* = 31,358) (Stahl et al., 2019); (2) schizophrenia (*N_cases_* = 35,476, *N_controls_* = 46,839) (Schizophrenia Working Group of the PGC, 2014); (3) coronary artery disease (*N_cases_* = 60,801, *N_controls_* = 123,504) (Nikpay et al., 2015); (4) ulcerative colitis (*N_cases_* = 12,366, *N_controls_* = 34,915) and (5) Crohn’s disease (*N_cases_* = 12,194, *N_controls_* = 34,915) (de Lange et al., 2017); (6) late onset Alzheimer’s disease (LOAD; *N_cases_* = 17,008, *N_controls_* = 37,154) (Lambert et al., 2013) (in the Supplemental Material we present results for a more recent GWAS with *N_cases_* = 71,880 and *N_controls_* = 383,378 (Jansen et al., 2018)); (7) amyotrophic lateral sclerosis (ALS) (*N*_*cases*_ = 12,577, *N_controls_* = 23,475) (Van Rheenen et al., 2016); (8) number of years of formal education (*N* = 293,723) (Okbay et al., 2016); (9) intelligence (*N* = 262,529) (Sniekers et al., 2017; Savage et al., 2018); (10) body mass index (UKB-GIANT 2018) (*N* = 690,519) (Yengo et al., 2018); (11) height (2010) (*N* = 133,735) (Yang et al., 2010); and height (2014) (*N* = 251,747) (Wood et al., 2014); (12) low- (*N* = 89,873) and (13) high-density lipoprotein (*N* = 94,295) (Willer et al., 2013); and (14) total cholesterol (*N* = 94,579) (Willer et al., 2013). Most participants were of European ancestry. (A spreadsheet giving data sources is provided in the Supplemental Material.) In the tables, we also report results for body mass index (GIANT 2015) (*N* = 233,554) (Locke et al., 2015), and height (UKB-GIANT 2018) (Yengo et al., 2018).

In Figure 2 and Supplemental Material, Figure S1 we report effective sample sizes, *N_eff_*, for the case-control GWASs. This is defined as *N_eff_* = 4/(1*/N_cases_*+1*/N_controls_*), so that when *N_cases_* = *N_controls_*, *N_eff_* = *N_cases_* +*N_controls_*= *N*, the total sample size, allowing for a straightforward comparison with quantitative traits.

**Figure 2:**
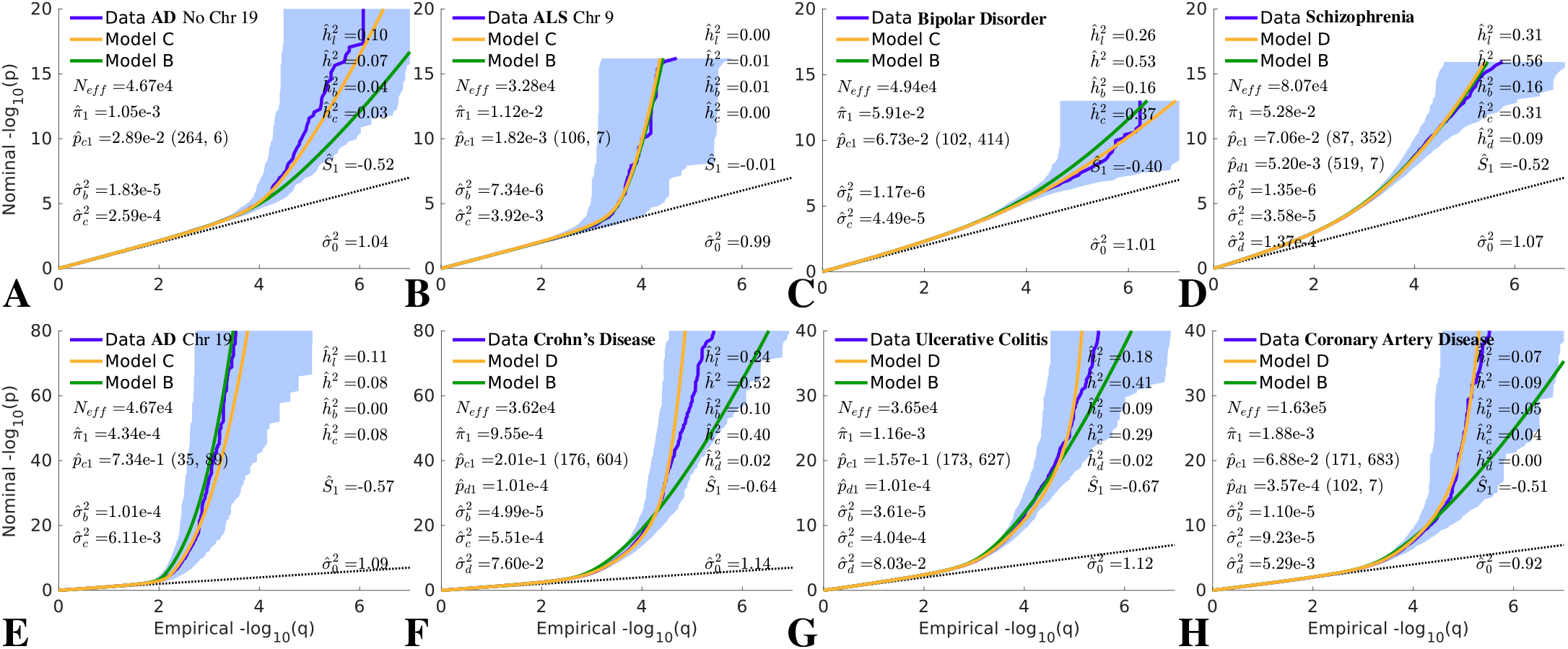
QQ plots of (pruned) z-scores for qualitative phenotypes (dark blue, 95% confidence interval in light blue) with model prediction (yellow). See Supplemental Material, Figures S15 to S22. The value given for *p*_*c*1_ is the amplitude of the full *p*_*c*_(*L*) function, which occurs at *L* = 1; the values (*m*_*c*_, *w*_*c*_) in parentheses following it are the total LD (*m*_*c*_) where the function falls to half its amplitude (the middle gray dashed lines in Figure 1 are examples), and the total LD width (*w*_*c*_) of the transition region (distance between flanking red dashed lines in Figure 1). Similarly for *p_d_*_1_ (*m*_*d*_, *w*_*d*_), where given. 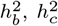, and 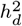 are the heritabilities associated with the “b”, “c”, and “d” Gaussians, respectively. *h*^2^ is the total SNP heritability, reexpressed as 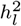 on the liability scale for binary phenotypes. Parameter values are also given in Table 1 and heritabilities are also in Table 3; numbers of causal SNPs are in Table 2. Reading the plots: on the vertical axis, choose a p-value threshold for typed or imputed SNPs (SNPs with z-scores; more extreme values are further from the origin), then the horizontal axis gives the proportion, q, of typed SNPs exceeding that threshold (higher proportions are closer to the origin). See also Supplemental Material, Figure S1, where the y-axis is restricted to 0 ⩽ −log_10_(*p*) ⩽ 10. The 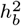 values reported here are from one component in the extended model; values for the exclusive basic model are reported as 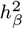 in (Holland et al., 2020).

**Table 2:** Numbers of causal SNPs. *n_causal_* is the total number of causal SNPs (from the 11 million in the reference panel); *n_b_*, *n*_*c*_, and *n_d_* are the numbers associated with the “b”, “c”, and “d” Gaussians, respectively. 95% confidence intervals are in 3, Table F1.

| Phenotype | $n_b$ | $n_c$ | $n_d$ | $n_{causal}$ |
| --- | --- | --- | --- | --- |
| SCZ 2014 | 5.6e5 | 2.3e4 | 3.0e3 | 5.82e5 |
| BIP | 6.2e5 | 2.6e4 | — | 6.51e5 |
| CD | 9.0e3 | 1.5e3 | 1 | 1.05e4 |
| UC | 1.1e4 | 1.4e3 | 2 | 1.27e4 |
| CAD | 2.0e4 | 1.0e3 | 5 | 2.07e4 |
| AD Chr19 | 80 | 33 | — | 113 |
| AD NoC19 | 1.1e4 | 294 | — | 1.12e4 |
| ALS Chr9 | 5.3e3 | 7 | — | 5.29e3 |
| Edu | 1.1e5 | 4.4e4 | 956 | 1.58e5 |
| IQ 2018 | 9.6e4 | 3.4e4 | 1.1e4 | 1.40e5 |
| Height 2010 | 9.4e3 | 1.9e3 | — | 1.13e4 |
| Height 2014 | 1.1e4 | 2.0e3 | — | 1.26e4 |
| Height 2018 | 2.0e4 | 7.6e3 | — | 2.75e4 |
| HDL | 2.7e4 | 264 | 15 | 2.79e4 |
| LDL | 6.4e4 | 463 | 14 | 6.43e4 |
| BMI 2015 | 1.7e4 | 68 | — | 1.69e4 |
| BMI 2018 | 2.6e4 | 88 | — | 2.57e4 |
| TC | 1.2e4 | 180 | 434 | 1.27e4 |

**Table 3:**
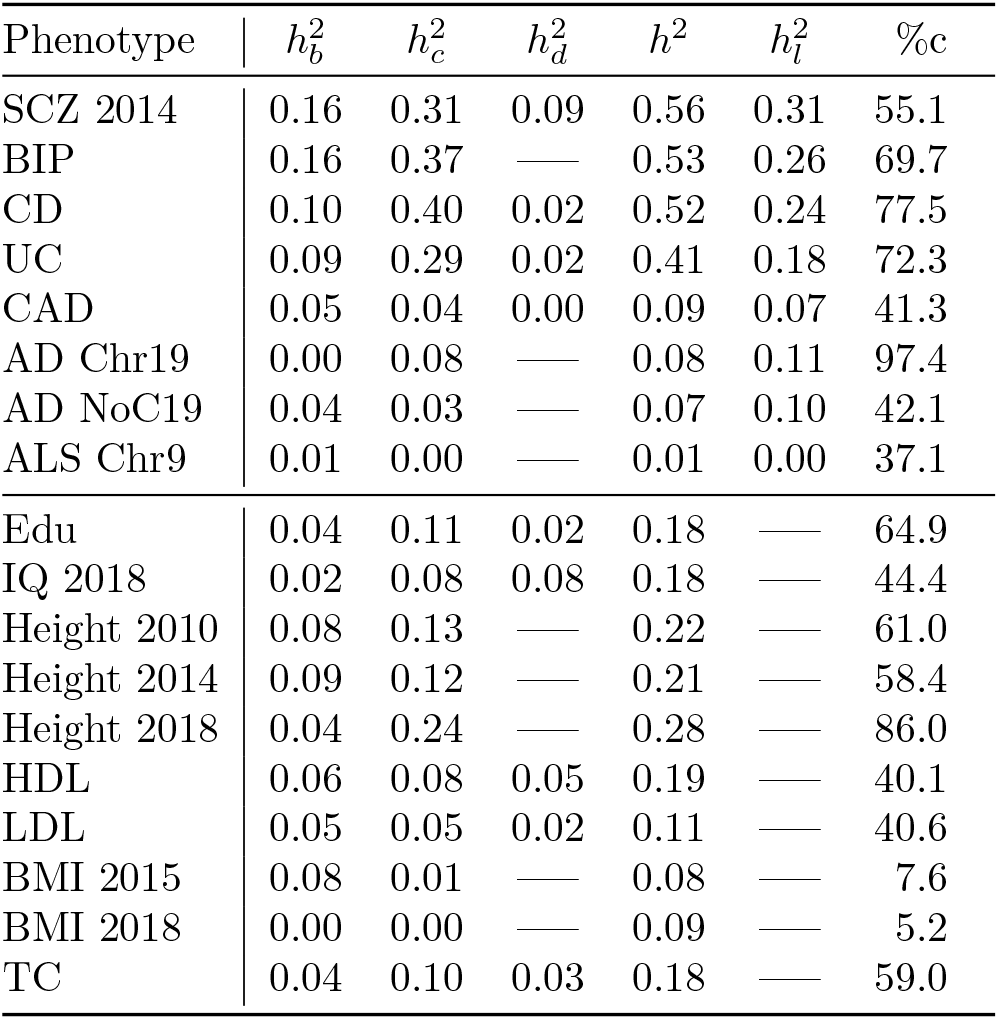
Heritabilities: *h*^2^ is the total additive SNP heritability, reexpressed on the liability scale as 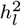 for the qualitative traits (upper section). 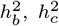, and 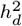 are the heritabilities associated with the “b”, “c”, and “d” Gaussians, respectively. The last column, labeled %c, gives the percentage of SNP heritability that comes from the “c” Gaussian. 95% confidence intervals are in 3, Table F2. A comparison of the total heritabilities with estimates from our basic model alone (Holland et al., 2020), and with estimates from other models (Zhang et al., 2018; Zeng et al., 2018; Schoech et al., 2019) are given in 3, Table B1.

| Phenotype | $h_b^2$ | $h_c^2$ | $h_d^2$ | $h^2$ | $h_l^2$ | %c |
| --- | --- | --- | --- | --- | --- | --- |
| SCZ 2014 | 0.16 | 0.31 | 0.09 | 0.56 | 0.31 | 55.1 |
| BIP | 0.16 | 0.37 | — | 0.53 | 0.26 | 69.7 |
| CD | 0.10 | 0.40 | 0.02 | 0.52 | 0.24 | 77.5 |
| UC | 0.09 | 0.29 | 0.02 | 0.41 | 0.18 | 72.3 |
| CAD | 0.05 | 0.04 | 0.00 | 0.09 | 0.07 | 41.3 |
| AD Chr19 | 0.00 | 0.08 | — | 0.08 | 0.11 | 97.4 |
| AD NoC19 | 0.04 | 0.03 | — | 0.07 | 0.10 | 42.1 |
| ALS Chr9 | 0.01 | 0.00 | — | 0.01 | 0.00 | 37.1 |
| Edu | 0.04 | 0.11 | 0.02 | 0.18 | — | 64.9 |
| IQ 2018 | 0.02 | 0.08 | 0.08 | 0.18 | — | 44.4 |
| Height 2010 | 0.08 | 0.13 | — | 0.22 | — | 61.0 |
| Height 2014 | 0.09 | 0.12 | — | 0.21 | — | 58.4 |
| Height 2018 | 0.04 | 0.24 | — | 0.28 | — | 86.0 |
| HDL | 0.06 | 0.08 | 0.05 | 0.19 | — | 40.1 |
| LDL | 0.05 | 0.05 | 0.02 | 0.11 | — | 40.6 |
| BMI 2015 | 0.08 | 0.01 | — | 0.08 | — | 7.6 |
| BMI 2018 | 0.00 | 0.00 | — | 0.09 | — | 5.2 |
| TC | 0.04 | 0.10 | 0.03 | 0.18 | — | 59.0 |

In estimating heritabilities on the liability scale for the qualitative phenotypes (Holland et al., 2020), we assumed prevalences of: BIP 0.5% (Merikangas et al., 2011), SCZ 1% (Speed et al., 2017), CAD 3% (Sanchis-Gomar et al., 2016), UC 0.1% (Burisch et al., 2013), CD 0.1% (Burisch et al., 2013), AD 14% (for people aged 71 and older in the USA (Plassman et al., 2007; Alzheimer’s Association, 2018)), and ALS 5 × 10^−5^ (Mehta et al., 2018).

Confidence intervals for parameters were estimated using the inverse of the observed Fisher information matrix (FIM). The full FIM was estimated for up to eight parameters used in model C, and for the remaining parameters that extend the analysis to model D the confidence intervals were approximated ignoring off-diagonal elements.

Additionally, the *w_d_* parameter was treated as fixed quantity, the lowest value allowing for a smooth transition of the *p_d_*(*L*) function to 0 (see Supplemental Material, Figure S5; for CD, UC, and TC, however, the function *p_d_*(*L*) was a constant (=*p_d_*(1)). For the derived quantities *h*^2^ and *n_causal_*, which depend on multiple parameters, the covariances among the parameters, given by the off-diagonal elements of the inverse of the FIM, were incorporated. Numerical values are in 3 Tables E1–E3, and 3 Tables F1–F2.

In order to carry out realistic simulations (i.e., with realistic heterozygosity and LD structures for SNPs), we used HAPGEN2 (Li and Stephens, 2003; Spencer et al., 2009; Su et al., 2011) to generate genotypes for 10^5^ samples; we calculated SNP MAF and LD structure from 1000 simulated samples.

### 1.3. Total Linkage Disequilibrium

Sequentially moving through each chromosome in contiguous blocks of 5,000 SNPs in the reference panel, for each SNP in the block we calculated its Pearson *r*^2^ correlation coefficients (that arise from linkage disequilibrium, LD) with all SNPs in the central block itself and with all SNPs in the pair of flanking blocks of size up to 25,000 each. For each SNP we calculated its total linkage disequilibrium (TLD), given by the sum of LD *r*^2^’s thresholded such that if 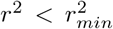 we set that to zero 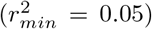.The fixed window size corresponds on average to a window of ±8 centimorgans. This is deliberately larger than the 1-centimorgan window used to define LD Score (Bulik-Sullivan et al., 2015), because the latter appears to exclude a noticeable part of the LD structure.

In applying the model to summary statistics, we calculated histograms of TLD (using 100 bins) and ignoring SNPs whose TLD was so large that their frequency was less than a hundredth of the respective histogram peak; typically this amounted to restricting to SNPs for which TLD ⩽ 600. We also ignored summary statistics of SNPs for which MAF ⩽ 0.01.

Since we are estimating a dozen or fewer parameters from millions of data points, it is reasonable to coarse-grain the data. Knowing the TLD and heterozygosity (H) of each SNP, we divided the full set of GWAS SNPs into a H×TLD coarse-grained grid; we found that 10×10 is more than sufficient for converged results.

### 1.4. Model PDF

When implementing the discrete Fourier transform (DFT) to calculate model *a posteriori* probabilities for z-score outcomes for a single SNP, we discretize the range of possible z-scores into the ordered set of *n* (equal to a power of values *z*_1_,…, *z_n_* with equal spacing between neighbors given by Δ*z* (*z_n_* = −*z*_1_ − Δ*z*, and *z_n/_*_2+1_ = 0). Taking *z*_1_ = −38 allows for the minimum p-values of 5.8 × 10^−316^ (near the numerical limit); with *n* = 2^10^, Δ*z* = 0.0742. Given Δ*z*, the Nyquist critical frequency is 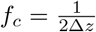, so we consider the Fourier transform function for the z-score pdf at *n* discrete values *k*_1_,…, *k_n_*, with equal spacing between neighbors given by Δ*k*, where *k*_1_ = −*f_c_* (*k_n_* = −*k*_1_ − Δ*k*, and *k_n/_*_2+1_ = 0; the DFT pair Δ*z* and Δ*k* are related by Δ*z*Δ*k* = 1*/n*).

In our earlier work on the “b” model (Holland et al., 2020) we gave an expression, denoted *G*(*k*_*j*_), for the Fourier transform of the genetic contribution to a z-score, where *k_j_* is the discrete Fourier variable described above. In constructing the *a posteriori* pdf for a z-score, the extra complexity in the “c” and “d” models presented in the current work requires a modification only in this term.

The set of typed SNPs are put in a relatively fine 10×10 grid, called the “H-L” grid, based on their heterozygosity and total LD (coarser grids are refined until converged results are achieved, and 10 × 10 is more than adequate). Given a typed SNP in LD with many tagged SNPs in the reference panel (some of which might be causal), divide up those tagged SNPs based on their LD with the typed SNP into *w_max_* LD-*r*^2^ windows (we find that *w_max_* = 20, dividing the range 0 ⩽ *r*^2^ ⩽ 1 into 20 bins, is more than adequate to obtain converged results); let 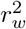 denote the LD of the *w*^th^ bin, *w* = 1*,…, w_max_*. Denote the number of tagged SNPs in window *w* as *n_w_* and their mean heterozygosity as *H_w_*. Then, given model “b” parameters *π*_1_, 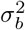, and 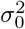, and sample size *N*, *G*(*k*_*j*_) is given by

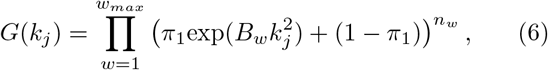

where

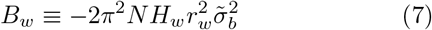

and 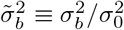 For model “c”, this becomes

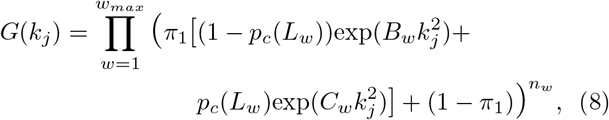

where

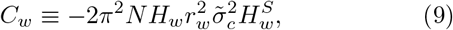

*S* is the selection parameter, 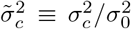 (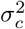 is defined in Eq. 2), *L_w_* is the mean total LD of reference SNPs in the *w* bin, and *p*_*c*_(*L_w_*) is the sigmoidal function (see Eq. 3) giving, when multiplied by *π*_1_, the prior probability of reference SNPs with this TLD being causal (with effect size drawn from the “c” Gaussian). For model “d”, *G*(*k_j_*)
becomes

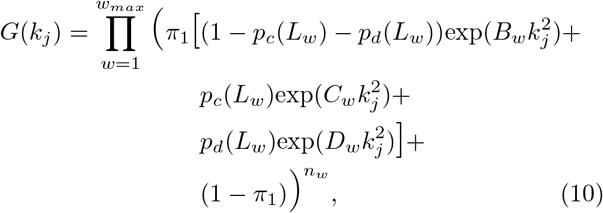

where

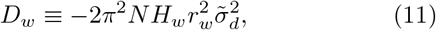

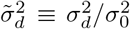 (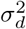is defined in Eq. 5), and *p_d_*(*L_w_*) is the sigmoidal function (again, see Eq. 3) giving, when multiplied by *π*_1_, the prior probability of reference SNPs with this TLD being causal (with effect size drawn from the “d” Gaussian).

Let 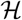 denote the LD and heterozygosity structure of a particular SNP, a shorthand for the set of values {*n_w_, H_w_, L_w_*: *w* = 1,…, *w_max_*} that characterize the SNP, and let 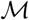 denote the set of model parameters for whichever model “b”, “c”, or “d” – is implemented. The Fourier transform of the environmental contribution, denoted *E_j_* ≡ *E*(*k_j_*), is

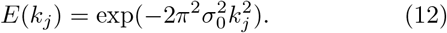

Let **F_z_** = (*G*_1_*E*_1_,…, *G_n_E_n_*), where *G_j_* ≡ *G*(*k_j_*), denote the vector of products of Fourier transform values, and let 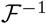 denote the inverse Fourier transform operator. Then for the SNP in question, the vector of pdf values, **pdf_z_**, for the uniformly discretized possible z-score outcomes *z*_1_*,…, z_n_* described above, i.e., **pdf_z_** = (*f*_1_,…, *f_n_*) where 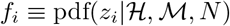, is

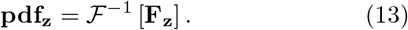

Thus, the i^th^ element **pdf_z*i*_** = *f_i_* is the *a posteriori* probability of obtaining a z-score value *z_i_* for the SNP, given the SNP’s LD and heterozygosity structure, the model parameters, and the sample size.

### 1.5. Data Availability

File S2 contains detailed descriptions of all GWAS data used, all of which is publicly available.

## 2. Results

### 2.1. Phenotypes

Summary QQ plots for pruned z-scores are shown in Figure 2 for seven binary phenotypes (for AD we separate out chromosome 19, which contains the APOE gene), and Figure 3 for seven quantitative phenotypes (including two separate GWAS for height), with model parameter values in Table 1. An example of the breakdowns of a summary plot with respect to a 4×4 grid of heterozygosity×TLD (each grid a subset of a 10×10 grid) is in Figure 4 for HDL; similar plots for all phenotypes are in Supplemental Material, Figures S15-S29. For each phenotype, model selection (B, C, or D) was performed by testing the Bayesian information criterion (BIC) – see 3 Table D1. For comparison, all QQ figures include the basic (B) model in green; the extended model C or D, consistently demonstrating improved fits, in yellow; and the data in blue.

**Figure 3:**
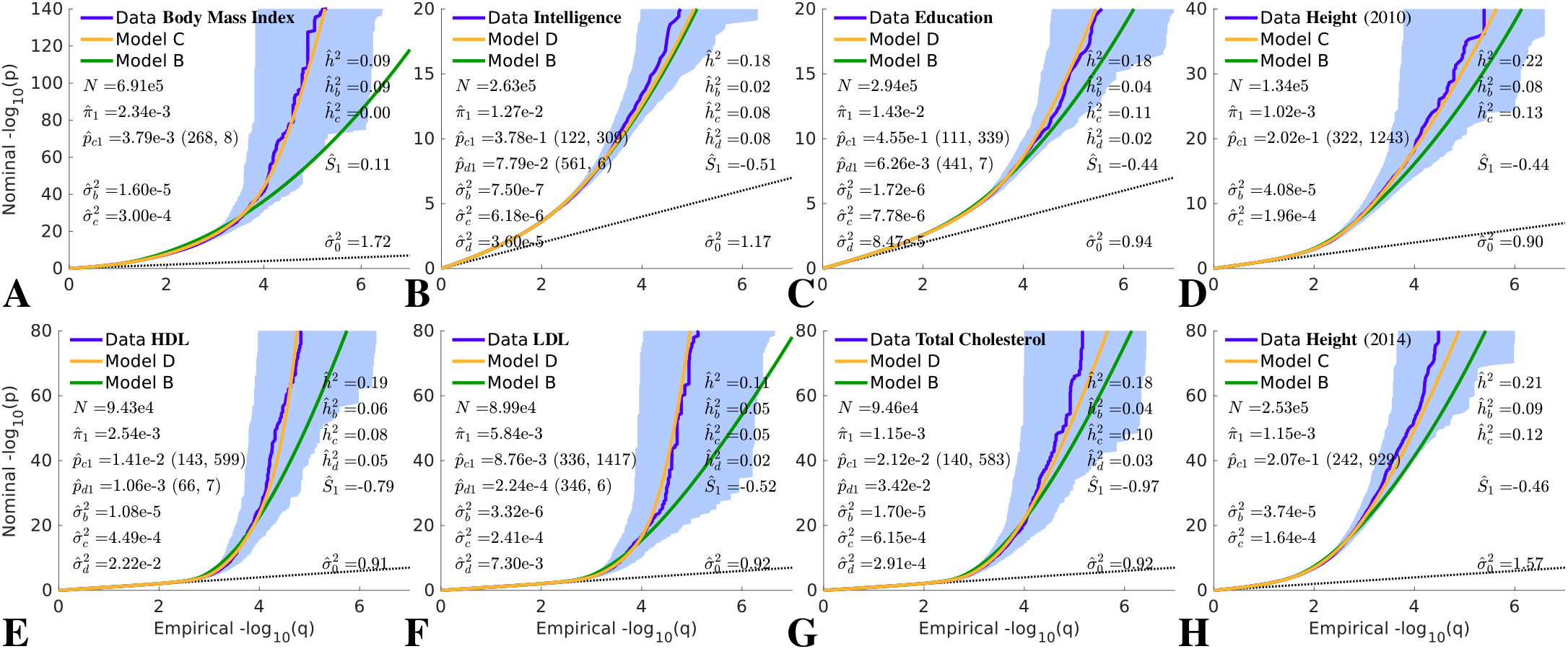
QQ plots of (pruned) z-scores for quantitative phenotypes (dark blue, 95% confidence interval in light blue) with model prediction (yellow). See Supplemental Material, Figures S23 to S29. For HDL, *p*_*c*_(*L*) = *p*_*c*1_ for all L; for bipolar disorder and LDL, *p_d_*(*L*) = *p_d_*_1_ for all L. See caption to Figure 2 for further description. See also Supplemental Material, Figure S2, where the y-axis is restricted to 0 ⩽ −log_10_(*p*) ⩽ 10. For (A) BMI, see also Supplemental Material, Figure S37.

**Figure 4:**
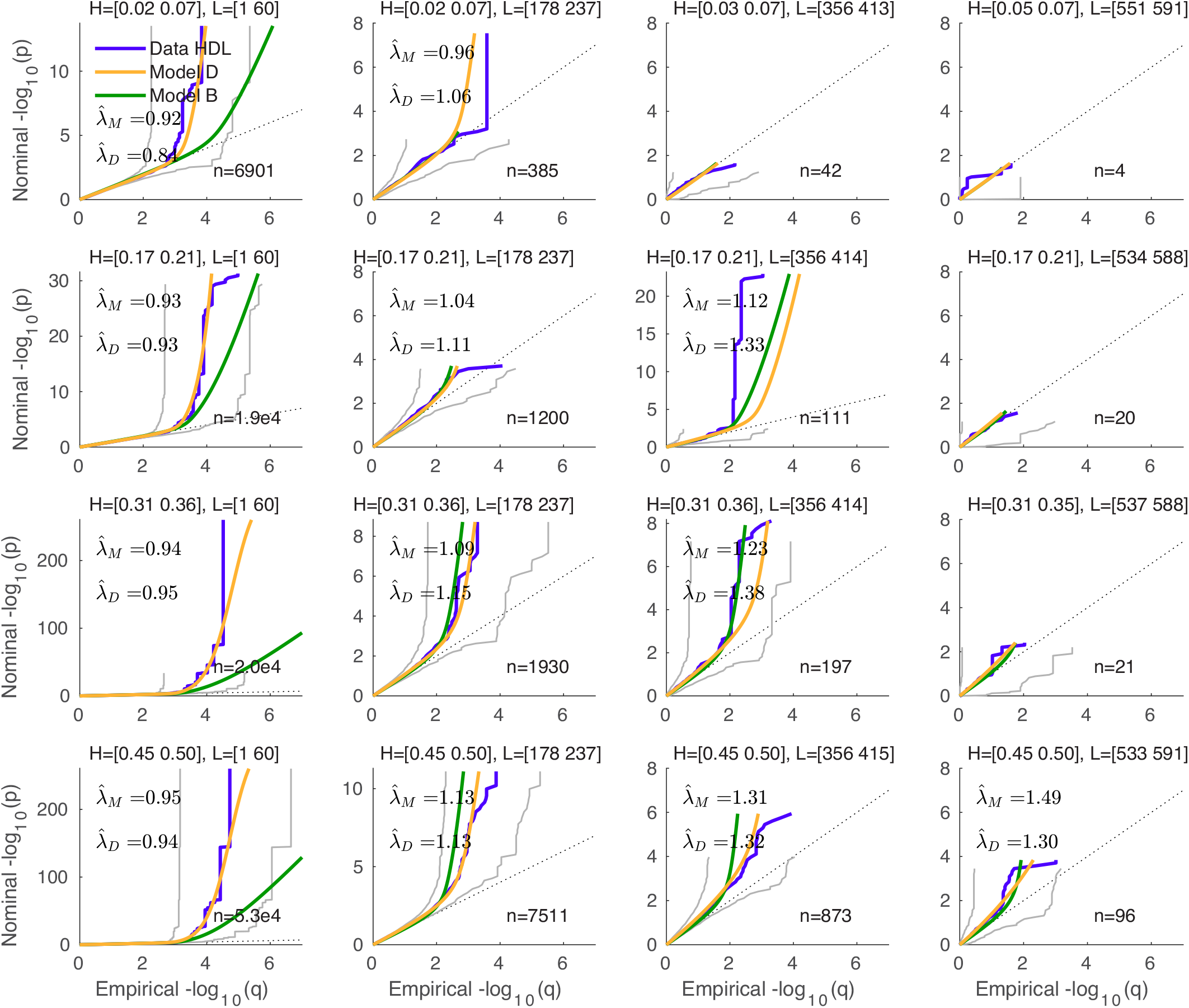
A 4×4 subset from a 10×10 heterozygosity×TLD grid of QQ plots for HDL; see Figure 3 (E) for the overall summary plot. Similar plots for all phenotypes are in Supplemental Material, Figures S15 to S29. The light gray curves are 95% confidence intervals for the data; 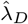 and 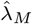 are the “genomic inflation factors” calculated from the QQ subplots, for the data and the model prediction, respectively; *n* is the number of SNPs; *H* is heterozygosity, *L* is total LD, and the square brackets give their ranges for GWAS SNPs in each grid element.

The distributions of z-scores for different phenotypes are quite varied. Nevertheless, for most phenotypes analyzed here, we find evidence for larger and smaller effects being distributed differently, with strong dependence on total LD, *L*, and heterozygosity, *H*.

For 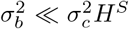, and so focusing on the “c” Gaussian (and “d” Gaussian if applicable for 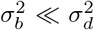), our model estimates an effective polygenicity as a one-dimensional function of *L*. We find that polygenicity is dominated by SNPs with low *L*. However, the degree of restriction varies widely across phenotypes, depending on the shapes and sizes of *p*_*c*_(*L*) and *p_d_*(*L*) in Eq. 5, the prior probabilities that a causal SNP belongs to the “c” and “d” Gaussians. These prior probabilities, multiplied by *π*_1_, are shown in Figure 1 and Supplemental Material, Figures S3-S5. Taking into account the underlying distribution of reference SNPs with respect to heterozygosity, these distributions lead to a varied pattern across phenotypes of the expected number of causal SNPs in equally-spaced elements in a two-dimensional H×TLD grid, as shown for height (2014) in Figure 5 C, and for all phenotypes in Supplemental Material, Figures S11-S14 (third columns). Further, for any given phenotype, the effect *sizes* of causal variants come from distributions whose variances can be widely different – by up to two orders of magnitude. Thus, given the prior probabilities (*p*_*b*_, *p*_*c*_, and *p_d_*) by which these distributions are modulated as a function of *L*, we are able to estimate the expected effect size per causal-SNP, *E*(*β*^2^), in each H×TLD grid element, as shown in Figure 5 D and Supplemental Material, Figures S11-S14 (fourth columns). In general, SNPs with lower *L* have larger *E*(*β*^2^). However, the selection parameter *S* in the “c” Gaussian has a large impact on *E*(*β*^2^) as a function of *H* (see Supplemental Material, Figure S9). As a result, for most phenotypes, we find that the effect sizes for low MAF causal SNPs (*H* < 0.05) are several times larger than for more common causal SNPs (*H* > 0.1). We find that heritability per causal-SNP is larger for lower *L*, a general pattern that follows from at least one of the prior probabilities, *p*_*c*_(*L*) or *p_d_*(*L*), being non-constant. However, because heritability per causal-SNP is proportional to *H*, we find that, even with negative selection parameter, *S* (and thus larger *E*(*β*^2^) for lower *H*), the heritability per causal-SNP is largest for the most common causal SNPs (*H* > 0.45).

**Figure 5:**
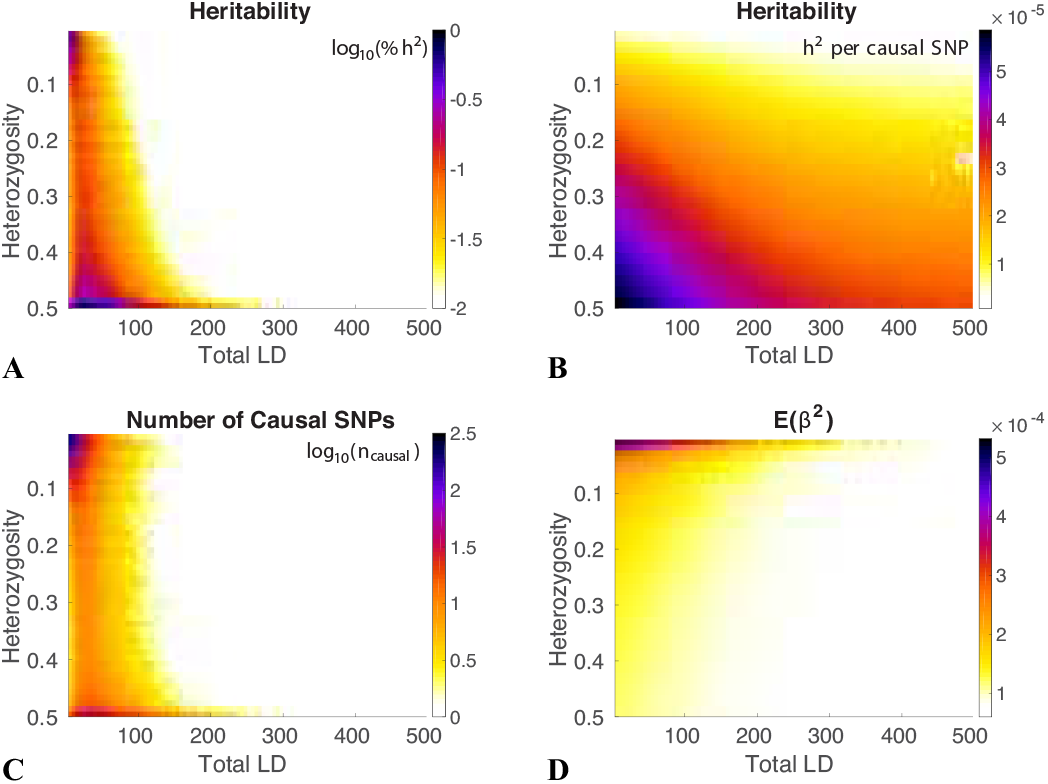
Model results for height (2014) using the BC model. The reference panel SNPs are binned with respect to both heterozygosity (*H*) and total LD (*L*) in a 50 × 50 grid for 0.02 ⩽ *H* ⩽ 0.5 and 1 ⩽ *L* ⩽ 500. Shown are model estimates of: (A) log_10_ of the percentage of heritability in each grid element; (B) for each element, the average heritability per causal-SNP in the element; (C) log_10_ of the number of causal SNPs in each element; and (D) the expected *β*^2^ for the element-wise causal SNPs. Note that *H* increases from top to bottom.

SNP heritability estimates from the extended model, as shown in Table 3, are uniformly larger than for the exclusive basic model. With the exception of BMI, generally a major portion of SNP heritability was found to be associated with the “selection” component, i.e., the “c” Gaussian, with a smaller contribution from the “d” Gaussian – see the right-most column in Table 3. In 3 Table B2 is a comparison of basic model estimates of the number of causal SNPs and heritability with the corresponding net contributions *n*_*c*_ + *n_d_* and 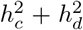 from the large effects components of the extended model. The basic model can be seen as a good approxiation to the large effects components of the extended model. Correspondingly, the “b” Gaussian as a components of the extended model now represents a relatively large number of much weaker effects.

### 2.2. Simulations

To test the specificity of the model for each real phenotype, we constructed simulations where, in each case, the true causal *β*’s (a single vector instantiation) for all reference panel SNPs were drawn from the overall distribution defined by the real phenotype’s parameters (thus being the “true” simulation parameters). We set up simulated phenotypes for 100,000 samples by adding noise to the genetic component of the simulated phenotype, and performed a GWAS to calculate z-scores. We then sought to determine whether the true parameters, and the component heritabilities, could reasonably be estimated by our model. In Figures 6 and 7 we show the results for the simulated case-control and quantitative phenotypes, respectively. Overall heritabilities were generally faithful to the true values (the values estimated for the real phenotypes) see 3, Table C2 – though for Crohn’s disease the simu lated value was overestimated due to the *h_d_* component. Note that for the case-control simulated phenotypes, the heritabilities on the observed scale, denoted 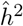 in Figure 6, should be compared with the corresponding values in Figure 2, not with 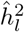, which denotes heritability on the liability scale, i.e., adjusted for population prevalence; note also that for case-control phenotypes, we implicitly assume the same proportion *P* of cases in the real and simulated GWAS (for the basic model, assuming the liability-scale model is correct, one can easily see from the definition of 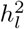 that discoverability 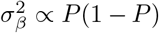; this carries over to *σ_b_*, *σ_c_*, and *σ_d_* used here). Polygenicities and discoverabilities were also generally faithfully reproduced. However, for ALS restricted to chromosome 9, and BMI, the selection parameter was incorrectly estimated, owing to the weak signal in these GWAS (e.g., for BMI, *π*_1_*p*_*c*_(1) ≃ 8 × 10^−6^, Fig. S4 (A), and only around 5% of SNP heritability was found to be associated with the “c” Gaussian, Table 3) and very low polygenicity (small number of causal SNPs) for the “c” Gaussian. Given the wide variety and even extreme ranges in parameters and heritability components across diverse simulated phenotypes, the multiple simulated examples provide checks with respect to each-other for correctly discriminating phenotypes by means of their model parameter estimates: the results for individual cases are remarkably faithful to the respective true values, demonstrating the utility of the model in distinguishing different phenotypes.

**Figure 6:**
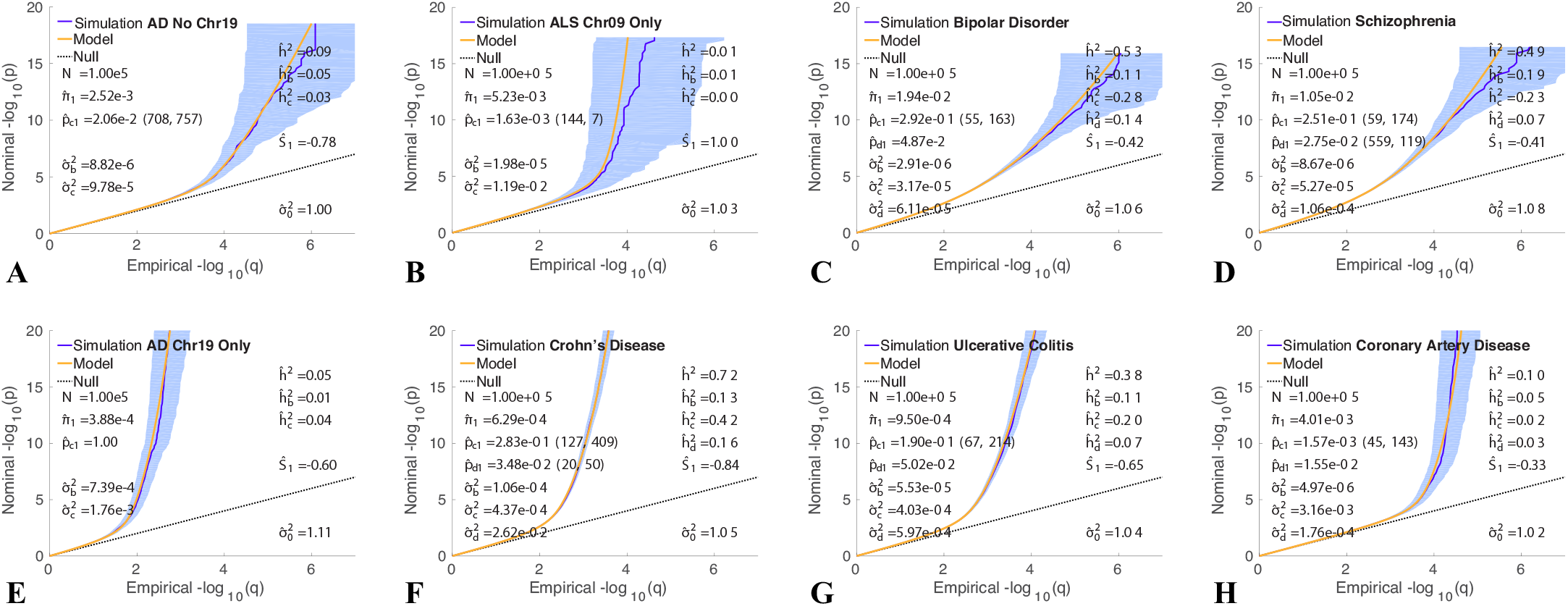
QQ plots of (pruned) z-scores for simulated qualitative phenotypes (dark blue, 95% confidence interval in light blue) with model prediction (yellow). See Figure 2. The value given for *p*_*c*1_ is the amplitude of the full *p*_*c*_(*L*) function, which occurs at *L* = 1; the values (*m*_*c*_, *w*_*c*_) in parentheses following it are the total LD (*m*_*c*_) where the function falls to half its amplitude (the middle gray dashed lines in Figure 1 are examples), and the total LD width (*w*_*c*_) of the transition region (distance between flanking red dashed lines in Figure 1). Similarly for *p_d_*_1_ (*m*_*d*_, *w*_*d*_), where given. 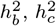 and 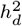 are the heritabilities associated with the “b”, “c”, and “d” Gaussians, respectively. *h*^2^ is the total SNP heritability, reexpressed as 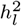 on the liability scale for binary phenotypes. Reading the plots: on the vertical axis, choose a p-value threshold for typed SNPs (SNPs with z-scores; more extreme values are further from the origin), then the horizontal axis gives the proportion, q, of typed SNPs exceeding that threshold (higher proportions are closer to the origin). See 3 Tables C1 and C2 for a comparison of numerical values between model estimates for real phenotypes and Hapgen-based simulations where the underlying distributions of simulation causal effects were given based on the real phenotype model parameters (with *σ*_0_ = 1).

**Figure 7:**
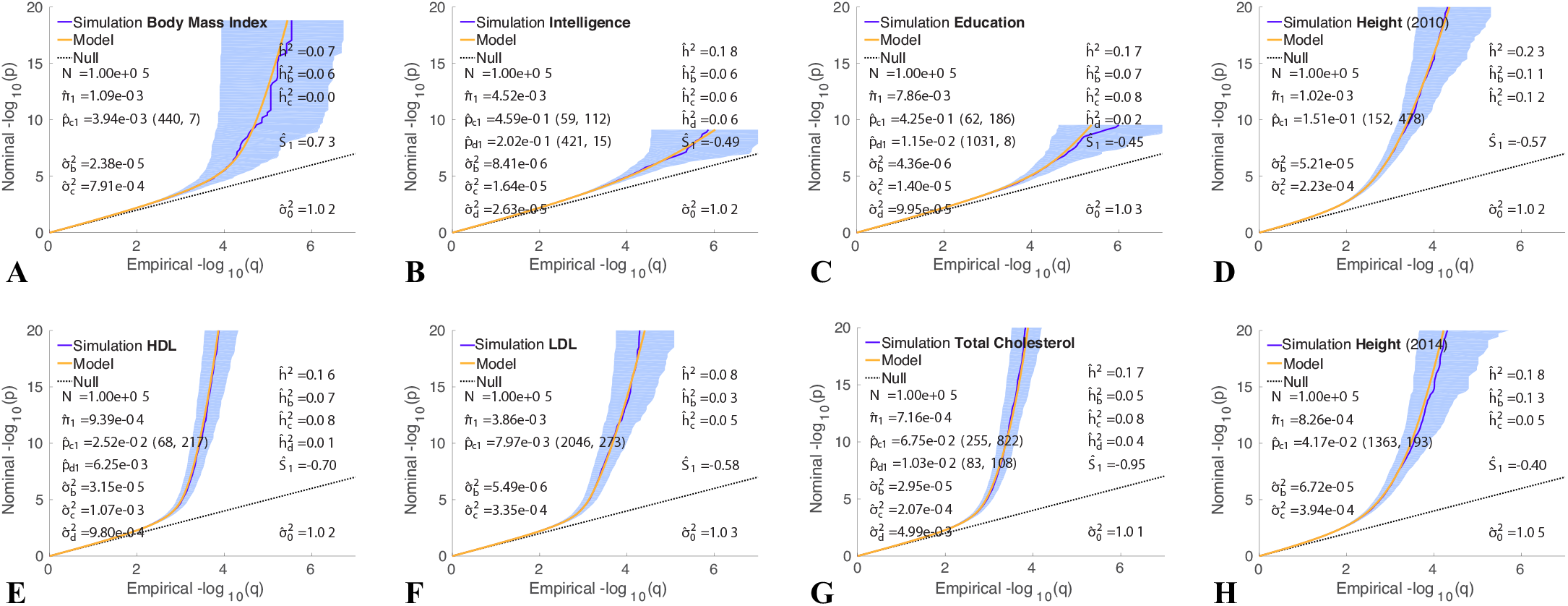
QQ plots of (pruned) z-scores for simulated quantitative phenotypes (dark blue, 95% confidence interval in light blue) with model prediction (yellow). See Figure 3. See caption to Figure 2 for further description. See 3 Tables C1 and C2 for a comparison of numerical values between model estimates for real phenotypes and Hapgen-based simulations where the underlying distributions of simulation causal effects were given based on the real phenotype model parameters (with *σ*_0_ = 1).

## 3. Discussion

We propose an extended Gaussian mixture model for the distribution of underlying SNP-level causal genetic effects in human complex phenotypes, allowing for the phenotype-specific distribution to be modulated by heterozygosity, *H*, and total LD, *L*, of the causal SNPs, and also allowing for independent distributions for large and small effects. The GWAS z-scores for the typed or imputed SNPs, in addition to having a random environmental and error contribution, arise through LD with the causal SNPs. Thus, taking the detailed LD and heterozygosity structure of the population into account by using a reference panel, we are able to model the distribution of z-scores and test the applicability of our model to human complex phenotypes.

Complex phenotypes are emergent phenomena arising from random mutations and selection pressure. Underlying causal variants come from multiple functional categories (Schork et al., 2013), and heritability is known to be enriched for some functional categories (Finucane et al., 2015; Gazal et al., 2017; Shadrin et al., 2019). Thus, it is likely that different variants will experience different evolutionary pressure either due to fitness directly or to pleiotropy with fitness related traits.

Here, we find evidence for markedly different genetic architectures across diverse complex phenotypes, where the effective polygenicity (or, equivalently, the prior probability that a SNP is causal with large effect) is a function of SNP total LD (*L*), and discoverability is multi-component and MAF dependent.

In contrast to previous work modeling the distribution of causal effects that took total LD and multiple functional annotation categories into account while implicitly assuming a polygenicity of 1 (Gazal et al., 2017), or took MAF into account while ignoring total LD dependence and different distributions for large and small effects (Zeng et al., 2018), or took independent distributions for large and small effects into account (which is related to incorporating multiple functional annotation categories) while ignoring total LD and MAF dependence, here we combine all these issues in a unified way, using an extensive underlying reference panel of ~11 million SNPs and an exact methodology using Fourier transforms to relate summary GWAS statistics to the posited underlying distribution of causal effects. We show that the distributions of all sets of phenotypic z-scores, including extreme values that are well within genome-wide significance, are accurately reproduced by the model, both at overall summary level and when broken down with respect to a 10×10 H×TLD grid even though the various phenotypic polygenicities and per causal-SNP heritabilities range over orders of magnitude. Improvement with respect to the basic model can be seen in the summary QQ plots Figs. 2 and 3, and also the H×TLD grids of subplots, such as Figure 4 for HDL (see also Supplemental Material, Figs. S15-S29). Compared with the basic mixture model, the extended model primarily improves the fits to data for GWAS SNPs with low total LD, in several cases for low heterozygosity as well, where very strong signals are evident. This improved model fit can be traced to the structure of the prior probability *p*_*c*_(*L*) and the variance for effect sizes, 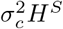, features absent in the basic model. But even when model fits are similar, the extended model fit taking selection pressure into account results in higher likelihood and lower Bayesian information criterion.

Negative selection can be expected to result in an increasing number of effects with increasing effect size at lower heterozygosity. It was found in (Gazal et al., 2017) which, in addition to analyzing total LD, modeled allele age and recombination rates – that common variants associated with complex traits are weakly deleterious to fitness, in line with an earlier model result that most of the variance in fitness comes from rare variants that have a large effect on the trait in question (Eyre-Walker, 2010). Thus, larger per-allele effect sizes for less common variants is consistent with the action of negative selection. Further, based on a model equivalent to Eq. 2 with *p*_*c*_ ≡ 1, it was argued in (Zeng et al., 2018), using forward simulations and a commonly used demographic model (Gravel et al., 2011), that negative values for the selection parameter, *S*, which leads to larger effects for rarer variants, is a signature of negative selection. Similar results were found for the related infinitesimal model (*p*_*c*_ ≡ 1 *and π*_1_ ≡ 1) in (Schoech et al., 2019).

We find negative selection parameter values for most traits, which is broadly in agreement with (Zeng et al., 2018) and (Schoech et al., 2019) with the exception of BMI, which we find can be modeled with two Gaussians with no or weakly positive (*S* ≳ 0) heterozygosity dependence, though it should be noted that the polygenicity for the larger-effects Gaussian (the “c” Gaussian with the *S* parameter) is very low, amounting to an estimate of < 100 common causal SNPs of large effect.

A similar situation (*S* ≃ 0) obtains with ALS restricted to chromosome 9. Here, the sample size is relatively low (12,577 cases), which contributes to the signal being weak, and we estimate only 7 common causal SNPs associated with the “c” Gaussian.

From twin studies, the heritability of sporadic ALS has been estimated as 0.61 (0.38 to 0.78) (Al-Chalabi et al., 2010); a clinically ascertained case series estimated the heritability to be between 0.40 and 0.45 (Wingo et al., 2011). Mutations of the chromosome 9 open reading frame 72 gene C9orf72 are implicated in both familial and sporadic ALS (Van Blitterswijk et al., 2012), but other chromosomes are also known to be involved (Chen et al., 2013). However, Supplemental Material, Figure S8, showing QQ plots accounting for all GWAS SNPs, implies that all the GWAS signal comes from chromosome 9. It should be noted that the mixture model, which is primarily focused on characterizing a distribution of undiscovered effect sizes, is designed to capture a broad-based polygenic contribution, i.e., not dominated by singular variants or effects from very large LD blocks. In implementing the model, we limit to SNPs with total LD<600 and randomly prune GWAS SNPs in LD blocks. From Figure 2 B and Figure S8, however, the few excluded GWAS SNPs clearly can not account for missing the preponderance of heritability. It is likely that the GWAS data lacks rare variants pertinent to ALS, and in addition is underpowered.

Our BMI results are not in agreement with the results of others. For the UKB (2015) data, Zeng et al. (2018) report a selection parameter *S* = −0.283 [−0.377, −0.224], *h*^2^ = 0.276, and *n_causal_* = 45*k*, while Schoech et al. (2019) report a selection parameter *α* = −0.24 [−0.38, −0.06], and *h*^2^ = 0.31. For the same GIANT 2015 dataset used here (Locke et al., 2015), Zhang et al Zhang et al. (2018) report *h*^2^ = 0.20 and *n_causal_* = 18*k*. It is not clear why our BMI results are in such disagreement with these, except to note that we have a two-component (versus single component) causal Gaussian and the majority of the heritability comes from the small-effects “b” Gaussian. For height, our selection parameter *S* = −0.46 (2014 GWAS) is in reasonable agreement with *S* = −0.422 reported in (Zeng et al., 2018), and with *α* = −0.45 reported in (Schoech et al., 2019).

Compared with our basic model results (Holland et al., 2020), we generally have found that using the extended model presented here, heritabilities show marked increases, with large contributions form the selection (“c”) Gaussian see 3, Table B1. For example, our basic-model heritability estimate for HDL was 7%, in close agreement with an earlier LDSC estimate (Speed and Balding, 2019) and the M2 model of Zhang et al. (2018) (9%), though the heritability from discovered loci was known to be 12% (Willer et al., 2013); our extended-model estimate is 19%, and the fit to the data is a dramatic improvement over the basic model, as can be seen in Figure 4 (it should be borne in mind that mixture models are designed to capture broad-based polygenic contributions to heritability, not outlier effects). However, due to the small effect sizes associated with the “b” Gaussian as a component of the extended model, for some phenotypes the overall number of causal SNPs can be considerably larger than previously estimated. The concept of effective number of causal SNPs, *M_e_*, in O’Connor et al. (2019), roughly corresponds to *n*_*c*_ + *n_d_* here, though the relationship is not precise (see also 3); e.g., for schizophrenia, *M_e_* = 15k versus *n*_*c*_ + *n_d_* = 26k. Supplemental Material, Figures S11-S14 capture the breakdown in SNP contributions to heritability as a function of heterozygosity and total linkage disequilibrium.

From Eq. 4, *β* is a weighted sum of contributions from Gaussians of different variance. Since we find *σ_b_ < σ_c_* and *S* < 0 – which increasingly magnifies the difference in variance of the two Gaussians as *H* gets smaller – we find larger E(*β*^2^) for rarer variants. Also, as *p*_*c*_(*L*) increases (from 0) with decreasing *L*, for a given *H* we also find that as *L* decreases, per causal-SNP heritability and E(*β*^2^) increase, consistent with Gazal et al. (2017). These patterns can be seen in Supplemental Material, Figures S11-S14 (second and fourth columns). The per causal-SNP contribution to heritability (second columns) is found to be more smoothly varying across common and low-frequency variants than E(*β*^2^), in broad agreement with O’Connor et al. (2019). It was also found in Gazal et al. (2017) that more recent common alleles have both lower LLD and larger per-SNP heritability (all SNPs causal); since selection has had less time to remove recent deleterious alleles, larger per-SNP heritability from SNPs with lower LLD was indicative of negative selection.

Generally, we find evidence for the existence of genetic architectures where the per causal-SNP heritability is larger for more common SNPs with lower total LD. But the trend is not uniform across phenotypes – see Supplemental Material, Figures S11-S14, second columns. The observed-scale heritability estimates given by 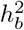 correspond to effects not experiencing much selection pressure. The new final values of *h*^2^ presented here result from a model that, compared with the basic Gaussian mixture model it is an extension of, gives better fits between data and model prediction of the summary and detailed QQ plots, and thus constitute more accurate estimates of SNP heritability. For 2010 and 2014 height GWASs, we obtain very good consistency for the model parameters and therefore heritability, despite considerable difference in apparent inflation. The 2018 height GWAS (Yengo et al., 2018) has a much larger sample size (almost three quarters of a million); the slightly different parameter estimates for it might arise due to population structure not fully captured by our model (Sohail et al., 2019; Berg et al., 2019). Our *h*^2^ estimates for height, however, remain consistently lower than other reported results (e.g., *h*^2^ = 0.33 in (Zhang et al., 2018), *h*^2^ = 0.527 in (Zeng et al., 2018), *h*^2^ = 0.61 in (Schoech et al., 2019)). For educational attainment, our heritability estimate agrees with (Zeng et al., 2018) (*h*^2^ = 0.182), despite our using a sample more than twice as large. The difference in selection parameter value, *S* = −0.335 in Zeng et al. (2018) versus *S* = −0.44 here, might partially be explained by the model differences (Zeng et al use one causal Gaussian with MAF-dependence but no LD dependence). A comparison of the total heritabilities reported here with estimates from the work of others (Zhang et al., 2018; Zeng et al., 2018; Schoech et al., 2019), in addition to our basic model alone (Holland et al., 2020), are given in 3, Table B1.

For most traits, we find strong evidence that causal SNPs with low heterozygosity have larger effect sizes (*S* < 0 in Table 1; the effect of this as an amplifier of 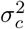 in Eq. 5 is illustrated in Supplemental Material, Figure S9) see also Supplemental Material, Figures S11-S14, fourth columns. Thus, negative selection seems to play an important role in most phenotypes-genotypes. This is also indicated by the extent of the region of finite probability for variants of large effect sizes (which are enhanced by having *S* ≲ −0.4) being relatively rare (low *H*), which will be greater for larger *p*_*c*1_, the amplitude of the prior probability for the “c” Gaussian (see Supplemental Material, Figures S3 and S4).

The “b” Gaussian in Eq. 4 or 5 does not involve a selection parameter: effect size variance is independent of MAF. Thus, causal SNPs associated with this Gaussian are likely undergoing neutral (or very weakly negative) selection. It should be noted that in all traits examined here, whether or not there is evidence of negative selection (*S* < 0), the effect size variance of the “b” Gaussian is many times smaller – sometimes by more than an order of magnitude – than that for the “c” Gaussian. Thus, it appears there are many causal variants of weak effect undergoing neutral (or very weakly negative) selection. For the nine phenotypes where the “d” Gaussian could be implemented, its variance parameter was several times larger than that of the “c” Gaussian. However, the amplitude of the prior probability for the “d” Gaussian, *p_d_*_1_, was generally much smaller than the amplitudes of the prior probabilities for the “b” or “c” Gaussians, which translated into a relatively small number of causal variants with very large effect associated with this Gaussian. (Due to lack of power, in three instances – CD, UC, and TC – *p_d_*(*L*) was treated as a constant, i.e., independent of *L*.) Interestingly, intelligence had the highest number of causal SNPs associated with this Gaussian, while the extent of total LD for associated SNPs was also liberal (*m_d_* = 561; see also Supplemental Material, Figure S5). It is possible that some of these SNPs are undergoing positive selection, but we did not find direct evidence of that.

A limitation of this work is that we do not take SNP functional categories into account. Inclusion of SNP annotation – e.g., by having a set of model parameters for each of many functional categories, and subdividing each SNP’s total LD into contributions from these categories – is important for deriving more biologically-informed interpretations of genetic effects. However, for the summary quantities estimated here, annotation is not expected to have a large impact; indeed, O’Connor et al. (2019) conclude that the accuracy of S-LD4M is not contingent on modeling annotation. Another limitation, a feature of many large GWAS, is that rare and disease-specific SNPs are not included. Thus, our analysis strictly applies only to the spectrum of relatively common SNPs. Indeed, our results point to the importance of rare variants in order to more comprehensively study the evolutionary architecture of complex phenotypes.

We find a diversity of genetic architectures across multiple human complex phenotypes where SNP total LD plays an important role in effect size distribution. In general, lower total LD SNPs are more likely to be causal with larger effects. Furthermore, for most phenotypes, while taking total LD into account, causal SNPs with lower MAF have larger effect sizes. These phenomena are consistent with models of the action of negative selection. Additionally, for all phenotypes, we find evidence of neutral selection operating on SNPs with relatively weak effect. We did not find direct evidence of positive selection. Compared with the basic Gaussian mixture model, which did not take heterozygosity or total LD into account in the distribution of effect sizes, the extended model consistently provided a much better fit to the distribution of GWAS summary statistics, thus providing more accurate estimates of genetic quantities of interest. Future work will explore SNP functional annotation categories and their differential roles in human complex phenotypes.

## Appendix A Related LDAK and LDSR models

The LDAK model of Speed et al also incorporates two predetermined fixed factors that scale the variance of the causal effect size distribution: SNP-specific local LD weightings and “information scores”, the latter a quantification of SNP quality (Speed et al., 2017). The local LD weighting (which remains implicitly dependent on MAF) is designed to scale down the effect size in the context of the infinitesimal model, to try to compensate for the effect itself being replicated, in empirical z-scores, through LD with neighboring SNPs. It should be noted that *this* problem of LD is implicitly taken care of in our basic model by: (1) using a point-normal distribution for non-causal SNPs; (2) an extensive underlying reference panel; and (3) directly modeling the effects of LD on z-scores in our PDF for GWAS summary statistics. The LDAK local LD weighting scheme is quite distinct from exploring the possibility of whether or not a SNP is causal depending on its TLD, which is the role of *π*_1_ × *p*_*c*_ in the extended model presented here (see below), or the degree to which its effect size is LD-dependent. Rather, within the infinitesimal framework, i.e., assuming a polygenicity of 1, and not explicitly taking the direct effects of LD on z-scores into account (as is done in our basic model), it unwinds the otherwise implicit problem – that SNPs with larger TLD would have inflated z-scores – by means of an assumed exponential decay function with respect to base-pair distance (such a function provides an approximation for the gross decay of long-range LD (Pritchard and Przeworski, 2001; Laird and Lange, 2010)).

In the LD Score regression model, which is another instance of the infinitesimal framework, Finucane et al. (2015) introduce an annotation-weighted LD score *l*(*j, c*) for variant *j* in LD with reference SNPs with (potentially overlapping) categorical annotation *c*, finding that different categories of SNPs are differentially enriched for heritability. Gazal et al. (2017) extend this to include continuous-valued annotations, and also introduce rank-based inverse normal transformation of the LD score – called level of LD (LLD) – which is calculated independently for different MAF bins and, in this way, unlike LD Score, is intended to be independent of MAF, with the effect size variance for SNP *j* given as a sum of terms: a baseline parameter common to all SNPs, plus one of ten parameters depending on which of ten bins the MAF of *j* is in, plus another parameter times the LLD of *j*.

Schoech et al. (2019) implement a variation of their main model, again in the infinitesimal framework, by augmenting their variance with an additional factor (1 − *τ* ·*LLD_j_*), searching over five values of the new parameter *τ* along with the original 20 values of the selection parameter in a fixed 2D grid.

Another variation on LD Score regression is “stratified LD fourth moments regression” (S-LD4M) from O’Connor et al. (2019). The authors argue that negative selection limits the contribution to heritability of SNPs with large per-allele effect (corresponding to *β* here), and that SNPs with relatively small per-allele effects can show up in GWAS as having relatively large chi-squared statistics. They introduce a new definition of polygenicity, *M_e_/M*, where *M* is the total number of SNPs and *M_e_* is the *effective* number of causal SNPs: if *f_i_* is the fractional contribution of SNP *i* to heritability 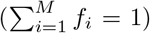, then *M_e_* can be understood as 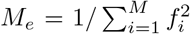. *M_e_* lies between 1 and M and will be weighted toward the number of SNPs with large contributions to heritability; the more even the distribution of *f_i_* the larger *M_e_* will be. Negative selection will not have any obvious effect on polygenicity *per se* (in the usual sense of counting up all contributing SNPs), but it will affect *M_e_*. In the absence of negative selection, the authors hypothesize that the effect-size distribution will be dominated by a relatively small number of large effect loci, resulting in a small *M_e_*. Under negative selection, critical large effects will not become common, thus limiting their contributions to heritability (smaller *f_i_* due to smaller heterozygosity), leading to a more uniform distribution of *f_i_* across all causal SNPs: the contributions to heritability of the many small effects from common alleles will not be drowned out by a small number of very large effects, thus leading to a much larger *M_e_* than in the absence of negative selection. It is in this sense – negative selection causing a flattening of the distribution of SNP contributions to heritability – that “extreme polygenicity of complex traits is explained by negative selection.”

## Appendix B Comparison of model heritability estimates

**Table B1:** Comparison of our current extended model heritability estimates (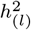, see Table 3) with estimates from: our earlier basic model (dressed with a tilde, 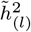) (Holland et al., 2020); the two- and three-component Gaussian models in (Zhang et al., 2018), denoted with subscript *M*2 and *M*3, respectively; the single Gaussian model with selection parameter in (Zeng et al., 2018), subscripted *MS*; and the single Gaussian infinitesimal model (polygenicity=1) with selection parameter in (Schoech et al., 2019), subscripted *α*. *M*2 and *M*3 use the HapMap3 reference panel (Consortium et al., 2010) with 1.07 million common SNPs (MAF) 0.05); *MS* uses an Affymetrix panel with 483,634 SNPs (MAF*>* 0.01) on UK Biobank data; our results are based on a 1000 Genomes Phase3 reference panel with 11 million SNPs (MAF) 0.002). SE denotes standard error. 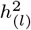 is our heritability estimate (see Table 3); those obtained from *M*2, *M*3, and *MS* are labeled 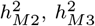, and 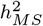, respectively. For quantitative phenotypes, values are on the observed scale; for binary phenotypes, values are on the liability scale, using the same population prevalence as used for Table 3. It is important to note that our heritability estimates are corrected for inflation, by dividing by the inflation parameter 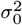; this is not done for *M*2, *M*3 or *MS*. ^*^Disease prevalences are given as a percentage in parentheses. For binary phenotypes, let 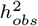 denote the heritability on the observed 0-1 scale (this is *h*^2^ in Figure 2). Let *P* denote the proportion of cases in the study: *P* = *N_cases_/*(*N_cases_* + *N_controls_*). Then the heritability on the log-odds-ratio scale reported in (Zhang et al., 2018) is 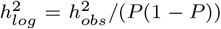. The transformation between the observed and liability scale is given by Eq. 39 in (Holland et al., 2020). ^†^N is the total sample size for quantitative traits; for qualitative traits, *N_eff_* = 4/(1*/N_cases_* + 1*/N_controls_*) – see main text.

| Phenotype* | $N_{(eff)}^\dagger$ | $h_{(l)}^2$ (SE) | $\tilde{h}_{(l)}^2$ (SE) | $h_{M2}^2$ (SE) | $h_{M3}^2$ (SE) | $h_{MS}^2$ (SE) | $h_\alpha^2$ (SE) |
| --- | --- | --- | --- | --- | --- | --- | --- |
| Schizophrenia (1.0%) | 8.07E4 | 0.31 (0.01) | 0.21 (0.001) | 0.29 (0.013) | — | — | — |
| Bipolar Disorder (0.5%) | 4.94E4 | 0.26 (0.01) | 0.16 (0.001) | 0.24 (0.027) | — | — | — |
| Crohn's Disease (0.1%) | 3.62E4 | 0.24 (0.01) | 0.18 (0.001) | 0.17 (0.021) | 0.23 (0.026) | — | — |
| Ulcerative Colitis (0.1%) | 3.65E4 | 0.18 (0.01) | 0.11 (0.001) | 0.10 (0.015) | 0.13 (0.020) | — | — |
| CAD (2011; 3.0%) | 6.57E4 | — | — | 0.07 (0.013) | 0.07 (0.012) | — | — |
| CAD (2015; 3.0%) | 1.63E5 | 0.07 (0.01) | 0.03 (0.001) | — | — | — | — |
| AD (14.0%) | 4.67E4 | 0.21 (0.10) | 0.15 (0.013) | 0.07 (0.024) | 0.10 (0.021) | — | — |
| Education (2016) | 2.94E5 | 0.18 (0.05) | 0.12 (0.001) | 0.13 (0.006) | — | — | — |
| Education (UKB) | 1.25E5 | — | — | — | — | 0.18 (0.004) | 0.15 (0.01) |
| Intelligence (2017) | 7.80E4 | — | — | 0.22 (0.015) | — | — | — |
| Intelligence (2018) | 2.63E5 | 0.18 (0.02) | 0.13 (0.001) | — | — | — | — |
| Height (2010) | 1.34E5 | 0.22 (0.01) | 0.17 (0.002) | 0.30 (0.014) | 0.32 (0.015) | — | — |
| Height (2014) | 2.53E5 | 0.21 (0.01) | 0.17 (0.001) | — | 0.33 (0.011) | — | — |
| Height (2018) | 7.08E5 | 0.28 (0.01) | 0.19 (0.001) | — | — | — | — |
| Height (UKB) | 1.26E5 | — | — | — | — | 0.53 (0.003) | 0.61 (0.00) |
| HDL | 9.43E4 | 0.19 (0.03) | 0.07 (0.000) | 0.09 (0.015) | 0.11 (0.010) | — | — |
| LDL | 8.99E4 | 0.11 (0.01) | 0.06 (0.002) | 0.08 (0.016) | 0.11 (0.011) | — | — |
| BMI (2010) | 1.24E5 | — | — | 0.20 (0.011) | 0.20 (0.010) | — | — |
| BMI (GIANT 2015) | 2.34E5 | 0.08 (0.01) | 0.07 (0.001) | — | 0.13 (0.005) | — | — |
| BMI (GIANT-UKB 2018) | 9.91E5 | 0.09 (0.01) | 0.27 | — | — | — | — |
| BMI (UKB 2015) | 1.26E5 | — | — | — | — | 0.28 (0.004) | 0.31 (0.00) |
| Total Cholesterol | 9.46E4 | 0.18 (0.08) | 0.09 (0.002) | 0.09 (0.013) | 0.12 (0.012) | — | — |

**Table B2:**
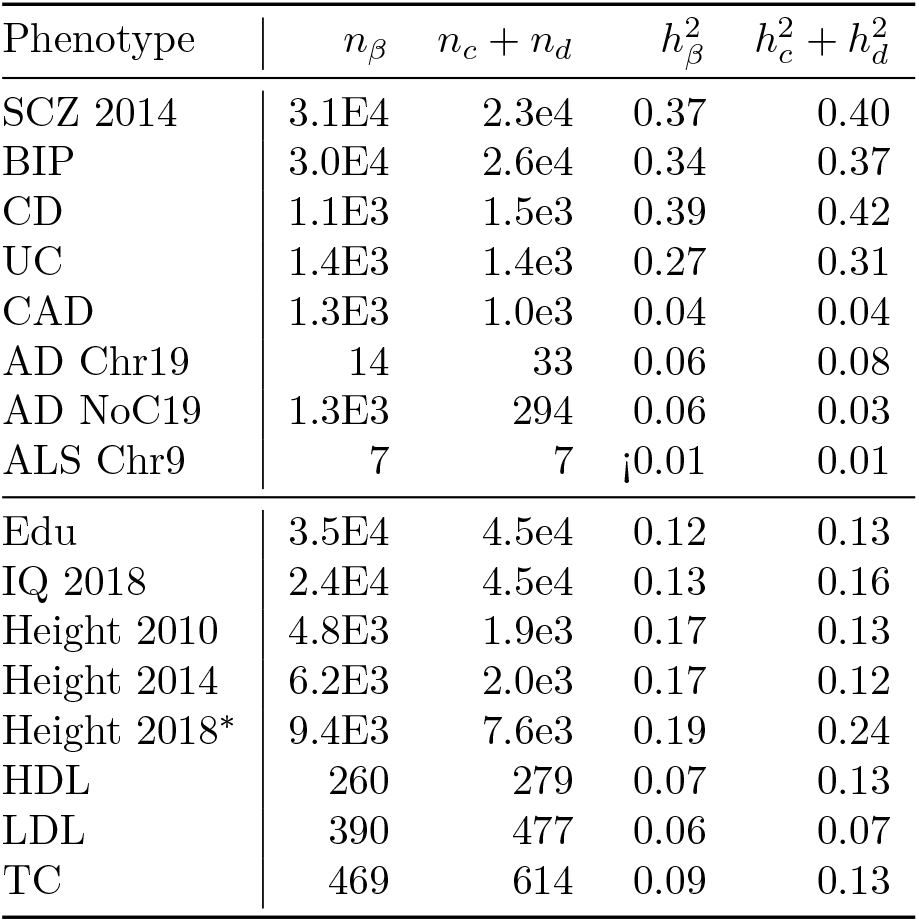
Comparison of the basic model estimates (Holland et al., 2020) of the number of causal SNPs (here denoted *n_β_*) and heritability (here denoted 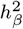) with the corresponding net contributions *n*_*c*_ + *n_d_* and 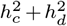 from the large effects components (“c” and “d”) of the extended model. (All disease heritabilities here are on the observed scale; see Table 3.) For BMI (GIANT 2015), it is the “b” component of the extended model that dominates, and a comparison with the basic model gives: *n_β_* = 7.5*E*3, *n_b_* = 1.7*E*4; 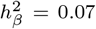, 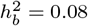 *Model C for Height 2018.

## Appendix C Comparison of model parameters for phenotypes and Hapgen-based simulations

**Table C1:** Comparison of model parameters for phenotypes and Hapgen-based simulations using the model parameters (with *σ*_0_ = 1) to produce the underlying distribution of effects. *π*_1_ is the overall proportion of the 11 million SNPs from the reference panel that are estimated to be causal. *π*_1_ × *p*_*c*_(*L* ⩾ 1) is the total prior probability multiplying the “c” Gaussian, which has variance 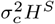, where *H* is the reference SNP heterozygosity. Note that *p*_*c*_(*L*) is just a sigmoidal curve, and can be characterized quite generally by three parameters: the value *p*_*c*1_ ≡ *p*_*c*_(1) at *L* = 1; the total LD value *L* = *m_c_* at the mid point of the transition, i.e., *p*_*c*_(*m_c_*) = *p*_*c*1_/2 (see the middle gray dashed lines in Figure 1, which shows examples of *π*_1_ times the function *p*_*c*_(*L*)); and the width *w_c_* of the transition, defined as the distance (in *L*) between where the curve falls to 95% and 5% of *p*_*c*1_ (distance between the flanking red dashed lines in Figure 1). The “d” Gaussian is similarly defined (but without the *H^S^* dependence in the variance). Note that for AD Chr19, AD NoC19, and ALS Chr9, *π*_1_ is the fraction of reference SNPs on chromosome 19, on the autosome excluding chromosome 19, and on chromosome 9, respectively. See Figures 6 and 7, and Supplemental Material, Figures S1 and S2.

| Phenotype | $\pi_1$ | $\sigma_b^2$ | $\sigma_c^2$ | $S$ | $\pi_1 p_{c1}$ | $m_c$ | $w_c$ | $\sigma_d^2$ | $\pi_1 p_{d1}$ | $m_d$ | $w_d$ | $\sigma_0^2$ |
| --- | --- | --- | --- | --- | --- | --- | --- | --- | --- | --- | --- | --- |
| SCZ 2014 | 5.28e-2 | 1.4e-6 | 3.6e-5 | -0.52 | 3.7e-3 | 87 | 352 | 1.4e-4 | 3.7e-4 | 519 | 7 | 1.07 |
| Simulation | 1.05e-2 | 2.9e-6 | 3.2e-5 | -0.42 | 2.6e-3 | 59 | 174 | 1.1e-4 | 6.8e-3 | 559 | 119 | 1.08 |
| BIP | 5.91e-2 | 1.2e-6 | 4.5e-5 | -0.40 | 4.0e-3 | 102 | 414 | — | — | — | — | 1.01 |
| Simulation | 6.86e-3 | 1.6e-5 | 3.9e-5 | -0.40 | 5.1e-3 | 64 | 162 | — | — | — | — | 1.07 |
| CD | 9.55e-4 | 5.0e-5 | 5.5e-4 | -0.64 | 1.9e-4 | 176 | 604 | 7.6e-2, | 2.0e-5 | — | — | 1.14 |
| Simulation | 6.29e-4 | 1.1e-4 | 4.4e-4 | -0.84 | 1.8e-4 | 127 | 409 | 2.6e-2, | 9.8e-3 | 20 | 50 | 1.05 |
| UC | 1.16e-3 | 3.6e-5 | 4.0e-4 | -0.67 | 1.9e-4 | 173 | 627 | 8.0e-2 | 1.6e-5 | — | — | 1.12 |
| Simulation | 9.50e-4 | 5.5e-5 | 4.0e-4 | -0.65 | 1.8e-4 | 67 | 214 | 6.0e-4 | 9.5e-3 | — | — | 1.04 |
| CAD | 1.88e-3 | 1.1e-5 | 9.2e-5 | -0.51 | 1.3e-4 | 171 | 683 | 5.3e-3 | 2.5e-5 | 102 | 7 | 0.92 |
| Simulation | 4.01e-3 | 5.0e-6 | 3.2e-3 | -0.33 | 6.4e-6 | 45 | 143 | 1.8e-4 | 2.4e-5 | — | — | 1.02 |
| AD Chr19 | 4.34e-4 | 1.0e-4 | 6.1e-3 | -0.57 | 3.2e-4 | 35 | 89 | — | — | — | — | 1.09 |
| Simulation | 3.88e-4 | 7.4e-4 | 1.8e-3 | -0.60 | 3.9e-4 | — | — | — | — | — | — | 1.11 |
| AD NoC19 | 1.05e-3 | 1.8e-5 | 2.6e-4 | -0.52 | 3.0e-5 | 264 | 6 | — | — | — | — | 1.04 |
| Simulation | 2.52e-3 | 8.8e-6 | 9.8e-5 | -0.78 | 5.3e-5 | 708 | 757 | — | — | — | — | 1.00 |
| ALS Chr9 | 1.12e-2 | 7.3e-6 | 3.9e-3 | -0.01 | 2.0e-5 | 106 | 6 | — | — | — | — | 0.99 |
| Simulation | 5.23e-3 | 2.0e-5 | 1.2e-2 | 1.00 | 8.4e-6 | 144 | 7 | — | — | — | — | 1.03 |
| Edu | 1.43e-2 | 1.7e-6 | 7.8e-6 | -0.44 | 6.4e-3 | 111 | 339 | 8.5e-5 | 2.8e-3 | 441 | 7 | 0.94 |
| Simulation | 7.86e-3 | 4.4e-6 | 1.4e-5 | -0.45 | 3.4e-3 | 62 | 186 | 1.0e-4 | 4.7e-3 | 1031 | 8 | 1.03 |
| IQ 2018 | 1.27e-2 | 7.5e-7 | 6.2e-6 | -0.51 | 4.8e-3 | 122 | 309 | 3.6e-5 | 3.0e-2 | 561 | 6 | 1.17 |
| Simulation | 4.52e-3 | 8.4e-6 | 1.6e-5 | -0.49 | 2.1e-3 | 59 | 112 | 2.6e-5 | 9.2e-2 | 421 | 15 | 1.02 |
| Height 2010 | 1.02e-3 | 4.1e-5 | 2.0e-4 | -0.44 | 2.0e-4 | 322 | 1243 | — | — | — | — | 0.90 |
| Simulation | 1.02e-3 | 5.2e-5 | 2.2e-4 | -0.57 | 1.5e-4 | 152 | 478 | — | — | — | — | 1.02 |
| Height 2014 | 1.15e-3 | 3.7e-5 | 1.6e-4 | -0.46 | 2.4e-4 | 242 | 929 | — | — | — | — | 1.57 |
| Simulation | 8.26e-4 | 6.7e-5 | 3.9e-4 | -0.40 | 3.5e-5 | 1363 | 193 | — | — | — | — | 1.05 |
| HDL | 2.54e-3 | 1.1e-5 | 4.5e-4 | -0.79 | 3.6e-5 | 143 | 599 | 2.2e-2 | 1.5e-5 | 66 | 7 | 0.91 |
| Simulation | 9.39e-4 | 3.1e-5 | 1.1e-3 | -0.70 | 2.3e-5 | 68 | 217 | 9.8e-4 | 1.6e-4 | — | — | 1.02 |
| LDL | 5.84e-3 | 3.3e-6 | 2.4e-4 | -0.52 | 5.1e-5 | 336 | 1417 | 7.3e-3 | 1.9e-6 | 346 | 6 | 0.92 |
| Simulation | 3.86e-3 | 5.5e-6 | 3.3e-4 | -0.58 | 3.1e-5 | 2046 | 273 | — | — | — | — | 1.03 |
| BMI GIANT 2015 | 1.54e-3 | 2.2e-5 | 4.5e-4 | 0.00 | 6.8e-6 | 288 | 12 | — | — | — | — | 0.85 |
| Simulation | 1.09e-3 | 2.4e-5 | 7.9e-4 | 0.73 | 4.3e-6 | 440 | 7 | — | — | — | — | 1.02 |
| TC | 1.15e-3 | 1.7e-5 | 6.2e-4 | -0.97 | 2.4e-5 | 140 | 583 | 2.9e-4 | 7.1e-4 | — | — | 0.92 |
| Simulation | 7.16e-4 | 2.9e-5 | 2.1e-4 | -0.95 | 4.8e-5 | 255 | 822 | 5.0e-3 | 6.7e-4 | 83 | 108 | 1.01 |

**Table C2:**
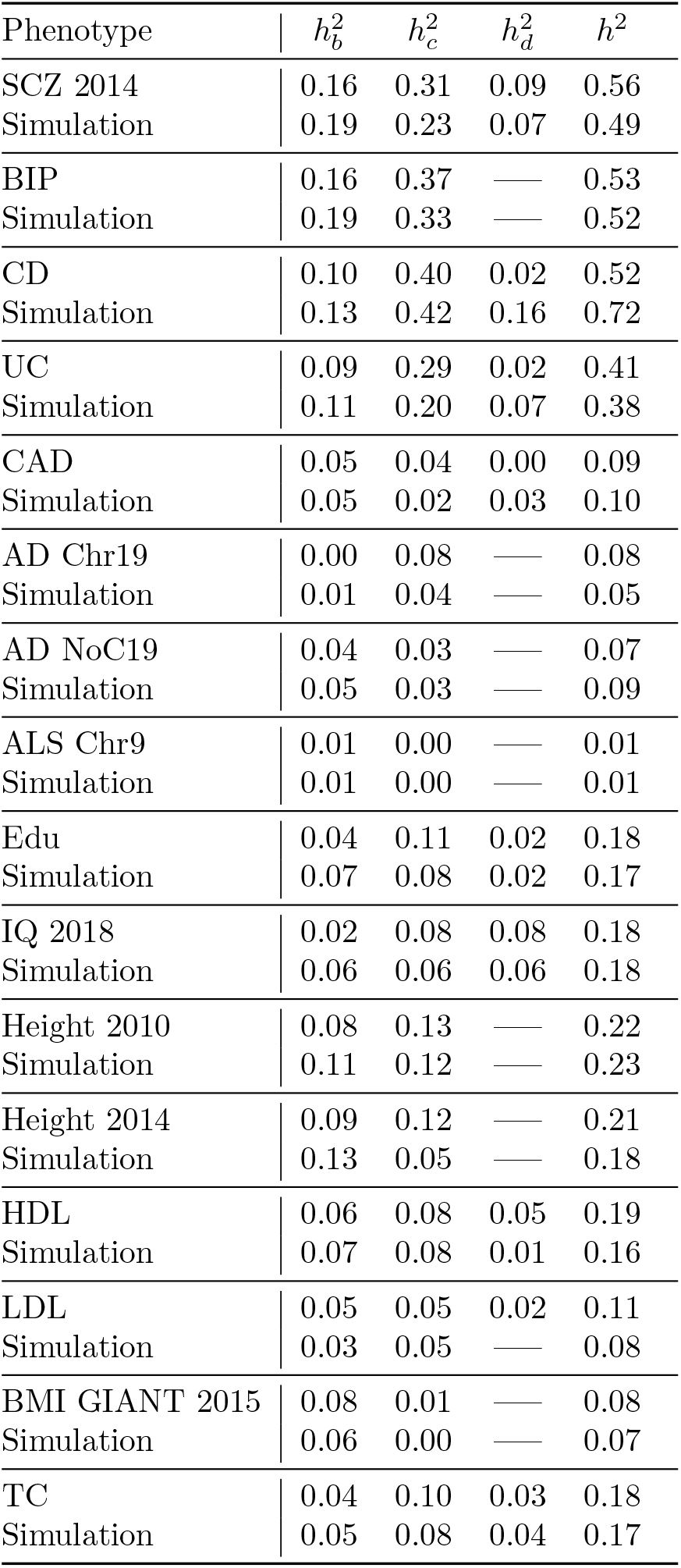
Comparison of model heritabilities for phenotypes and Hapgen-based simulations using the model parameters (with *σ*_0_ = 1) to produce the underlying distribution of effects. *h*^2^ is the total additive SNP heritability on the observed scale. 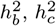, and 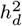 are theheritabilities associated with the “b”, “c”, and “d” Gaussians, respectively. See Figures 6 and 7, and Supplemental Material, Figures S1 and S2.

## Appendix D Bayesian information criterion (BIC) and model validity

**Table D1:** Bayesian information criterion (BIC) and model validity. The danger in adding extra parameters to a model is that it will over-fit the data. For a given model with *k* parameters and *n* degrees of freedom (number of independent z-scores), the BIC value is defined as 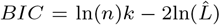, where 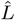 is the estimated likelihood of the data given the parameters. All else equal, models with lower *BIC* are preferred. Here we show BIC values for three models: the 3-parameter model B with only the “b” Gaussian (*π*_1_*, σ_b_, σ*_0_); the 8-parameter model C with both the “b” with “c” Gaussians (Eq. 4); and the 12-parameter model D with “b”, “c” and “d” Gaussians (Eq. 5). Since our cost function returns the product over heterozygosity-total LD bins of the log likelihood probabilities, i.e., the product over H-L elements *cost_HL_* = ln(pdf(*z*-score data in H-L bin | model params)), the overall 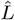 is calculated as minus the cost for approximately independent elements. For a more conservative estimate of *BIC*, we calculate this using aggressive pruning – selecting typed SNPs that are approximately independent, by setting a threshold for LD blocks at *r*^2^ = 0.1 and randomly selecting a typed SNP to represent that block, then averaged over 10 iterations. *C_flag_* and *D_flag_* indicate whether the increases in model complexity (relative to model B) are valid for the models C and D. †for model D applied to LDL, only 11 parameters can be considered for BIC to indicate an improvement over model C, i.e., *w_d_* is ignored as a parameter but rather treated as a fixed quantity giving a sharp yet smooth transition to 0 for the *p_d_*(*L*) function for large *L*.

| Phenotype | B (3) | C (8) | D (12) | B-C | C-D | $C_{flag}$ | $D_{flag}$ |
| --- | --- | --- | --- | --- | --- | --- | --- |
| SCZ 2014 | 345385 | 342862 | 342668 | 2523 | 194 | 1 | 1 |
| BIP | 315359 | 313187 | — | 2172 | — | 1 | 0 |
| CD | 337448 | 336225 | 336000 | 1223 | 225 | 1 | 1 |
| UC | 323298 | 322182 | 322071 | 1116 | 111 | 1 | 1 |
| CAD | 114633 | 113643 | 113496 | 989 | 147 | 1 | 1 |
| AD Chr19 | 104047 | 103606 | — | 441 | — | 1 | 0 |
| AD NoC19 | 285119 | 284942 | — | 177 | — | 1 | 0 |
| ALS Chr9 | 121378 | 121271 | — | 107 | — | 1 | 0 |
| Edu | 336485 | 333281 | 332820 | 3204 | 460 | 1 | 1 |
| IQ 2018 | 339400 | 338556 | 338392 | 844 | 164 | 1 | 1 |
| Height 2010 | 293877 | 292768 | 292778 | 1109 | -10 | 1 | 0 |
| Height 2014 | 402975 | 400877 | 400904 | 2098 | -27 | 1 | 0 |
| Height 2018 | 568229 | 562841 | 561496 | 5388 | 13454 | 1 | 1 |
| HDL | 288164 | 281404 | 281321 | 6760 | 83 | 1 | 1 |
| LDL<sup>†</sup> | 273299 | 270769 | 270760 | 2530 | 9 | 1 | 1 |
| BMI GIANT 2015 | 10885 | 10171 | — | 714 | — | 1 | 0 |
| BMI GIANT-UKB 2018 | 47507 | 37914 | — | 9593 | — | 1 | 0 |
| TC | 291141 | 288852 | 288738 | 2289 | 114 | 1 | 1 |
| TG | 274454 | 272227 | 272115 | 2227 | 112 | 1 | 1 |

## Appendix E 95% confidence intervals for model parameters

**Table E1:**
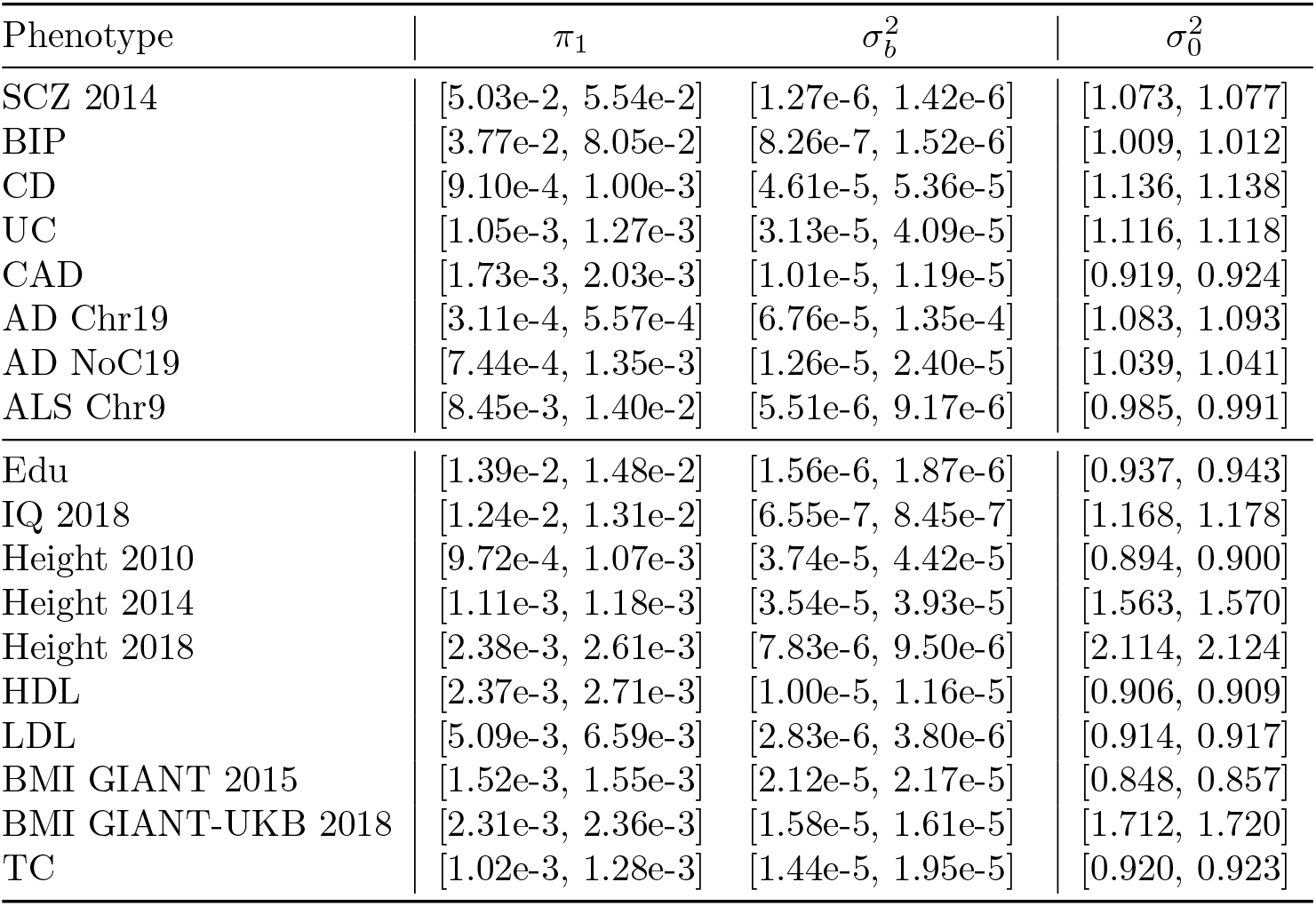
95% confidence intervals for model B parameters: *π*_1_, 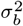 and 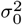. See Table 1. These confidence intervals are likely underestimates (too narrow), due to linkage disequilibrium and a consequent over-counting of the number of degrees of freedom (independent z-scores).

| Phenotype | $\pi_1$ | $\sigma_b^2$ | $\sigma_0^2$ |
| --- | --- | --- | --- |
| SCZ 2014 | [5.03e-2, 5.54e-2] | [1.27e-6, 1.42e-6] | [1.073, 1.077] |
| BIP | [3.77e-2, 8.05e-2] | [8.26e-7, 1.52e-6] | [1.009, 1.012] |
| CD | [9.10e-4, 1.00e-3] | [4.61e-5, 5.36e-5] | [1.136, 1.138] |
| UC | [1.05e-3, 1.27e-3] | [3.13e-5, 4.09e-5] | [1.116, 1.118] |
| CAD | [1.73e-3, 2.03e-3] | [1.01e-5, 1.19e-5] | [0.919, 0.924] |
| AD Chr19 | [3.11e-4, 5.57e-4] | [6.76e-5, 1.35e-4] | [1.083, 1.093] |
| AD NoC19 | [7.44e-4, 1.35e-3] | [1.26e-5, 2.40e-5] | [1.039, 1.041] |
| ALS Chr9 | [8.45e-3, 1.40e-2] | [5.51e-6, 9.17e-6] | [0.985, 0.991] |
| Edu | [1.39e-2, 1.48e-2] | [1.56e-6, 1.87e-6] | [0.937, 0.943] |
| IQ 2018 | [1.24e-2, 1.31e-2] | [6.55e-7, 8.45e-7] | [1.168, 1.178] |
| Height 2010 | [9.72e-4, 1.07e-3] | [3.74e-5, 4.42e-5] | [0.894, 0.900] |
| Height 2014 | [1.11e-3, 1.18e-3] | [3.54e-5, 3.93e-5] | [1.563, 1.570] |
| Height 2018 | [2.38e-3, 2.61e-3] | [7.83e-6, 9.50e-6] | [2.114, 2.124] |
| HDL | [2.37e-3, 2.71e-3] | [1.00e-5, 1.16e-5] | [0.906, 0.909] |
| LDL | [5.09e-3, 6.59e-3] | [2.83e-6, 3.80e-6] | [0.914, 0.917] |
| BMI GIANT 2015 | [1.52e-3, 1.55e-3] | [2.12e-5, 2.17e-5] | [0.848, 0.857] |
| BMI GIANT-UKB 2018 | [2.31e-3, 2.36e-3] | [1.58e-5, 1.61e-5] | [1.712, 1.720] |
| TC | [1.02e-3, 1.28e-3] | [1.44e-5, 1.95e-5] | [0.920, 0.923] |

**Table E2:** 95% confidence intervals for additional parameters (i.e., in addition to the three model B parameters) included in full model C: 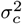, *S*, *p*_*c*1_, *m_c_*, and *w_c_*. See Table 1. These confidence intervals are likely underestimates (too narrow), due to linkage disequilibrium and a consequent over-counting of the number of degrees of freedom (independent z-scores).

| Phenotype | $\sigma_c^2$ | $S$ | $p_{c1}$ | $m_c$ | $w_c$ |
| --- | --- | --- | --- | --- | --- |
| SCZ 2014 | [3.43e-5, 3.73e-5] | [-0.55, -0.50] | [0.054, 0.088] | [86, 88] | [335, 369] |
| BIP | [4.32e-5, 4.66e-5] | [-0.43, -0.38] | [6.6e-2, 6.9e-2] | [100, 103] | [386, 441] |
| CD | [5.35e-4, 5.67e-4] | [-0.65, -0.62] | [0.19, 0.22] | [168, 181] | [575, 627] |
| UC | [3.91e-4, 4.17e-4] | [-0.68, -0.65] | [0.14, 0.18] | [165, 181] | [594, 658] |
| CAD | [8.73e-5, 9.73e-5] | [-0.53, -0.49] | [4.4e-2, 9.3e-2] | [165, 177] | [635, 732] |
| AD Chr19 | [5.17e-3, 7.05e-3] | [-0.66, -0.47] | [0.66, 0.81] | [28, 41] | [80, 97] |
| AD NoC19 | [2.31e-4, 2.87e-4] | [-0.57, -0.47] | [2.2e-2, 3.6e-2] | [263, 265] | — |
| ALS Chr9 | [2.39e-3, 5.45e-3] | [-0.18, 0.15] | [1.2e-3, 2.4e-3] | [104, 107] | — |
| Edu | [7.50e-6, 8.07e-6] | [-0.47, -0.40] | [0.31, 0.60] | [94, 127] | [276, 402] |
| IQ 2018 | [5.79e-6, 6.56e-6] | [-0.55, -0.47] | [0.36, 0.40] | [117, 122] | [294, 309] |
| Height 2010 | [1.87e-4, 2.05e-4] | [-0.47, -0.40] | [0.19, 0.21] | [315, 329] | [1137, 1348] |
| Height 2014 | [1.57e-4, 1.71e-4] | [-0.49, -0.43] | [0.2, 0.22] | [238, 246] | [872, 986] |
| Height 2018 | [8.66e-5, 9.08e-5] | [-0.45, -0.42] | [0.36, 0.38] | [206, 214] | [717, 760] |
| HDL | [4.08e-4, 4.91e-4] | [-0.85, -0.73] | [5.0e-3, 2.3e-2] | [142, 144] | [559, 639] |
| LDL | [2.10e-4, 2.72e-4] | [-0.59, -0.45] | [7.9e-3, 9.6e-3] | [336, 336] | [1401, 1433] |
| BMI GIANT 2015 | [4.47e-4, 4.60e-4] | [-0.01, 0.00] | [4.3e-3, 4.4e-3] | — | — |
| BMI GIANT-UKB 2018 | [2.96e-4, 3.05e-4] | [0.10, 0.11] | [3.7e-3, 3.8e-3] | — | — |
| TC | [5.56e-4, 6.74e-4] | [-1.04, -0.90] | [1.1e-2, 3.1e-2] | [139, 141] | [538, 627] |

**Table E3:** 95% confidence intervals for additional parameters (i.e., in addition to the eight model C parameters) included in full model D: 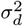, *p_d_*_1_, *m_d_*. See Table 1. These confidence intervals are likely underestimates (too narrow), due to linkage disequilibrium and a consequent over-counting of the number of degrees of freedom (independent z-scores).

| Phenotype | $\sigma_d^2$ | $p_{d1}$ | $m_d$ |
| --- | --- | --- | --- |
| SCZ 2014 | [1.33e-4, 1.42e-4] | [4.2e-4, 4.5e-4] | [512, 525] |
| BIP | — | — | — |
| CD | [6.96e-2, 8.24e-2] | [3.8e-4, 4.4e-4] | — |
| UC | [6.86e-2, 9.20e-2] | [2.7e-5, 3.6e-5] | — |
| CAD | [4.51e-3, 6.07e-3] | [3.8e-5, 5.1e-5] | [101, 104] |
| AD Chr19 | — | — | — |
| AD NoC19 | — | — | — |
| ALS Chr9 | — | — | — |
| Edu | [8.19e-5, 8.75e-5] | [3.4e-3, 3.7e-3] | [442, 448] |
| IQ 2018 | [3.55e-5, 3.66e-5] | [5.8e-2, 6.0e-2] | [563, 568] |
| Height 2010 | — | — | — |
| Height 2014 | — | — | — |
| Height 2018 | — | — | — |
| HDL | [1.88e-2, 2.56e-2] | [1.8e-5, 2.2e-5] | [63, 69] |
| LDL | [6.32e-3, 8.29e-3] | [2.0e-6, 2.5e-6] | [339, 352] |
| BMI GIANT 2015 | — | — | — |
| BMI GIANT-UKB 2018 | — | — | — |
| TC | [2.67e-4, 3.14e-4] | [7.9e-4, 9.3e-4] | — |

## Appendix F 95% confidence intervals for number of causal SNPs and heritability

**Table F1:** 95% confidence intervals for numbers of causal SNPs. *n_causal_* is the total number of causal SNPs (from the 11 million in the reference panel); *n_b_*, *n*_*c*_, and *n_d_* are the numbers associated with the “b”, “c”, and “d” Gaussians, respectively. See Table 2. These confidence intervals are likely underestimates (too narrow), due to linkage disequilibrium and a consequent over-counting of the number of degrees of freedom (independent z-scores).

| Phenotype | $n_b$ | $n_c$ | $n_d$ | $n_{causal}$ |
| --- | --- | --- | --- | --- |
| SCZ 2014 | [5.29e5, 5.83e5] | [1.73e4, 2.86e4] | [2.81e3, 3.11e3] | [5.54e5, 6.10e5] |
| BIP | [3.99e5, 8.52e5] | [1.66e4, 3.55e4] | — | [4.25e5, 8.78e5] |
| CD | [8.57e3, 9.44e3] | [1.43e3, 1.61e3] | [1, 2] | [1.01e4, 1.10e4] |
| UC | [1.02e4, 1.25e4] | [1.27e3, 1.60e3] | [1, 2] | [1.16e4, 1.39e4] |
| CAD | [1.81e4, 2.12e4] | [814, 1.20e3] | [4, 5] | [1.91e4, 2.23e4] |
| AD Chr19 | [50, 110] | [17, 50] | — | [67, 159] |
| AD NoC19 | [7.75e3, 1.41e4] | [1.72, 417] | — | [7.92e3, 1.46e4] |
| ALS Chr9 | [3.98e3, 6.58e3] | [4, 10] | — | [3.98e3, 6.59e3] |
| Edu | [1.02e5, 1.23e5] | [3.57e4, 5.29e4] | [888, 1.02e3] | [1.39e5, 1.77e5] |
| IQ 2018 | [9.19e4, 9.92e4] | [3.23e4, 3.61e4] | [1.03e4, 1.12e4] | [1.34e5, 1.46e5] |
| Height 2010 | [8.90e3, 9.91e3] | [1.66e3, 2.07e3] | — | [1.07e4, 1.18e4] |
| Height 2014 | [1.02e4, 1.09e4] | [1.85e3, 2.21e3] | — | [1.22e4, 1.30e4] |
| Height 2018 | [1.89e4, 2.08e4] | [7.20e3, 8.09e3] | — | [2.65e4, 2.86e4] |
| HDL | [2.58e4, 2.95e4] | [178, 350] | [13, 16] | [2.61e4, 2.98e4] |
| LDL | [5.56e4, 7.21e4] | [388, 539] | [12, 16] | [5.61e4, 7.26e4] |
| BMI GIANT 2015 | [1.67e4, 1.70e4] | [67, 69] | — | [1.68e4, 1.71e4] |
| BMI GIANT-UKB 2018 | [2.54e4, 2.59e4] | [87, 90] | — | [2.55e4, 2.60e4] |
| TC | [1.07e4, 1.34e4] | [132, 227] | [383, 485] | [1.13e4, 1.41e4] |

**Table F2:** 95% confidence intervals for heritabilities: *h*^2^ is the total additive SNP heritability, reexpressed on the liability scale as 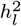 for the qualitative traits (upper section). 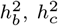, and 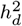 are the heritabilities associated with the “b”, “c”, and “d” Gaussians, respectively. See Table 3. These confidence intervals are likely underestimates (too narrow), due to linkage disequilibrium and a consequent over-counting of the number of degrees of freedom (independent z-scores).

| Phenotype | $h_b^2$ | $h_c^2$ | $h_d^2$ | $h^2$ | $h_l^2$ |
| --- | --- | --- | --- | --- | --- |
| SCZ 2014 | [0.16, 0.17] | [0.27, 0.35] | [0.08, 0.09] | [0.52, 0.60] | [0.29, 0.34] |
| BIP | [0.15, 0.17] | [0.34, 0.39] | — | [0.50, 0.56] | [0.25, 0.28] |
| CD | [0.10, 0.10] | [0.36, 0.44] | [0.02, 0.02] | [0.48, 0.56] | [0.22, 0.26] |
| UC | [0.09, 0.09] | [0.26, 0.33] | [0.02, 0.03] | [0.37, 0.44] | [0.17, 0.20] |
| CAD | [0.05, 0.05] | [0.03, 0.04] | [0.00, 0.00] | [0.08, 0.10] | [0.07, 0.08] |
| AD Chr19 | [0.00, 0.00] | [0.00, 0.22] | — | [0.00, 0.22] | [0.00, 0.30] |
| AD NoC19 | [0.02, 0.07] | [0.01, 0.06] | — | [0.02, 0.13] | [0.03, 0.17] |
| ALS Chr9 | [0.00, 0.01] | [0.00, 0.01] | — | [0.01, 0.02] | [0.00, 0.01] |
| Edu | [0.03, 0.06] | [0.02, 0.21] | [0.02, 0.02] | [0.07, 0.28] | — |
| IQ 2018 | [0.01, 0.02] | [0.05, 0.11] | [0.08, 0.09] | [0.14, 0.22] | — |
| Height 2010 | [0.08, 0.09] | [0.12, 0.14] | — | [0.20, 0.23] | — |
| Height 2014 | [0.08, 0.09] | [0.11, 0.13] | — | [0.20, 0.22] | — |
| Height 2018 | [0.04, 0.04] | [0.23, 0.25] | — | [0.27, 0.29] | — |
| HDL | [0.06, 0.06] | [0.01, 0.14] | [0.04, 0.06] | [0.13, 0.25] | — |
| LDL | [0.04, 0.05] | [0.03, 0.06] | [0.02, 0.02] | [0.10, 0.13] | — |
| BMI GIANT 2015 | [0.08, 0.08] | [0.01, 0.01] | — | [0.08, 0.09] | — |
| BMI GIANT-UKB 2018 | [0.09, 0.09] | [0.01, 0.01] | — | [0.09, 0.09] | — |
| TC | [0.04, 0.05] | [0.00, 0.27] | [0.02, 0.03] | [0.01, 0.34] | — |

## Appendix G Parameters for *p*_*c*_(*L*) and *p_d_*(*L*)

**Table G1:** Parameters for *p*_*c*_(*L*) and *p_d_*(*L*), the sigmoid function *y*(*x*) given in Eq. 3. The plots in Figure 1 and Supplemental Material, Figures S3-S5, are for *L* = *x* ⩾ 1. Thus, for example, *p*_*c*_(*L*) = *y*(*L*) = *y_cmax_/*(1 + exp((*L* − *x_cmid_*)*/x_cwidth_*)). For BIP, CD, UC, and TC, *p_d_* is constant: *p_d_* = *y_dmax_*.

| Phenotype | $y_{cmax}$ | $x_{cmid}$ | $x_{cwidth}$ | $y_{dmax}$ | $x_{dmid}$ | $x_{dwidth}$ |
| --- | --- | --- | --- | --- | --- | --- |
| SCZ 2014 | 0.8422 | -276.5 | 116.1 | 0.0052 | 518.0 | 1.0 |
| BIP | 1 | -360.2 | 137.4 | — | — | — |
| CD | 0.5 | -69.8 | 177.3 | 0.0001 | — | — |
| UC | 0.5 | -147.8 | 190.8 | 0.0001 | — | — |
| CAD | 0.552 | -433.4 | 222.8 | 0.000357 | 101.6 | 1.00 |
| AD Chr19 | 1 | 22.8 | 21.5 | — | — | — |
| AD NoC19 | 0.0289 | 264.0 | 1.0 | — | — | — |
| ALS Chr9 | 0.00182 | 105.2 | 1.0 | — | — | — |
| Edu | 0.804 | 25.2 | 91.7 | 0.00626 | 441.0 | 1.08 |
| IQ 2018 | 0.5 | 84.5 | 73.8 | 0.0779 | 561.0 | 1.00 |
| Height 2010 | 1 | -540.8 | 394.7 | — | — | — |
| Height 2014 | 1 | -393.7 | 294.4 | — | — | — |
| Height 2018 | 1 | -116.9 | 219.9 | — | — | — |
| HDL | 0.738 | -795.0 | 202.0 | 0.00106 | 65.6 | 1.02 |
| LDL | 0.876 | -2202.9 | 479.6 | 0.000224 | 345.5 | 1.00 |
| BMI GIANT 2015 | 0.00439 | 288.0 | 1.9 | — | — | — |
| BMI GIANT-UKB 2018 | 0.0038 | 267.1 | 1.2 | — | — | — |
| TC | 0.845 | -716.2 | 196.1 | 0.0342 | — | — |

## References

Al-Chalabi, A., Fang, F., Hanby, M. F., Leigh, P. N., Shaw, C. E., Ye, W., Rijsdijk, F., 2010. An estimate of amyotrophic lateral sclerosis heritability using twin data. Journal of Neurology, Neurosurgery & Psychiatry 81 (12), 1324–1326.

Alzheimer’s Association, 2018. 2018 alzheimer’s disease facts and figures. Alzheimer’s & Dementia 14 (3), 367–429.

Berg, J. J., Harpak, A., Sinnott-Armstrong, N., Joergensen, A. M., Mostafavi, H., Field, Y., Boyle, E. A., Zhang, X., Racimo, F., Pritchard, J. K., et al., 2019. Reduced signal for polygenic adaptation of height in uk biobank. eLife 8, e39725.

Bulik-Sullivan, B. K., Loh, P.-R, Finucane, H. K., Ripke, S., Yang, J., Patterson, N., Daly, M. J., Price, A. L., Neale, B. M., of the Psychiatric Genomics Consortium, S. W. G., et al., 2015. Ld score regression distinguishes confounding from polygenicity in genome-wide association studies. Nature genetics 47 (3), 291–295.

Burisch, J., Jess, T., Martinato, M., Lakatos, P. L., ECCO-EpiCom, 2013. The burden of inflammatory bowel disease in europe. Journal of Crohn’s and Colitis 7 (4), 322–337.

Chen, S., Sayana, P., Zhang, X., Le, W., 2013. Genetics of amyotrophic lateral sclerosis: an update. Molecular neurodegeneration 8 (1), 28.

Consortium, . G. P., et al., 2012. An integrated map of genetic variation from 1,092 human genomes. Nature 491 (7422), 56–65.

Consortium, . G. P., et al., 2015. A global reference for human genetic variation. Nature 526 (7571), 68–74.

Consortium, I. H. ., et al., 2010. Integrating common and rare genetic variation in diverse human populations. Nature 467 (7311), 52.

de Lange, K. M., Moutsianas, L., Lee, J. C., Lamb, C. A., Luo, Y., Kennedy, N. A., Jostins, L., Rice, D. L., Gutierrez-Achury, J., Ji, S.-G, et al., 2017. Genome-wide association study implicates immune activation of multiple integrin genes in inflammatory bowel disease. Nature genetics 49 (2), 256.

Devlin, B., Roeder, K., Dec 1999. Genomic control for association studies. Biometrics 55 (4), 997–1004.

Erbe, M., Hayes, B., Matukumalli, L., Goswami, S., Bowman, P., Reich, C., Mason, B., Goddard, M., 2012. Improving accuracy of genomic predictions within and between dairy cattle breeds with imputed high-density single nucleotide polymorphism panels. Journal of dairy science 95 (7), 4114–4129.

Eyre-Walker, A., 2010. Genetic architecture of a complex trait and its implications for fitness and genome-wide association studies. Proceedings of the National Academy of Sciences 107 (suppl 1), 1752–1756.

Fay, J. C., Wyckoff, G. J., Wu, C.-I, 2001. Positive and negative selection on the human genome. Genetics 158 (3), 1227–1234.

Finucane, H. K., Bulik-Sullivan, B., Gusev, A., Trynka, G., Reshef, Y., Loh, P.-R, Anttila, V., Xu, H., Zang, C., Farh, K., et al., 2015. Partitioning heritability by functional annotation using genome-wide association summary statistics. Nature genetics 47 (11), 1228.

Frei, O., Holland, D., Smeland, O. B., Shadrin, A. A., Fan, C. C., Maeland, S., OConnell, K. S., Wang, Y., Djurovic, S., Thompson, W. K., et al., 2019. Bivariate causal mixture model quantifies polygenic overlap between complex traits beyond genetic correlation. Nature communications 10 (1), 1–11.

Gazal, S., Finucane, H. K., Furlotte, N. A., Loh, P.-R, Palamara, P. F., Liu, X., Schoech, A., Bulik-Sullivan, B., Neale, B. M., Gusev, A., et al., 2017. Linkage disequilibrium–dependent architecture of human complex traits shows action of negative selection. Nature genetics 49 (10), 1421.

George, E. I., McCulloch, R. E., 1993. Variable selection via gibbs sampling. Journal of the American Statistical Association 88 (423), 881–889.

Gravel, S., Henn, B. M., Gutenkunst, R. N., Indap, A. R., Marth, G. T., Clark, A. G., Yu, F., Gibbs, R. A., Bustamante, C. D., Project, . G., et al., 2011. Demographic history and rare allele sharing among human populations. Proceedings of the National Academy of Sciences 108 (29), 11983–11988.

Hibar, D. P., Stein, J. L., Renteria, M. E., Arias-Vasquez, A., Desrivi`eres, S., Jahanshad, N., Toro, R., Wittfeld, K., Abramovic, L., Andersson, M., et al., 2015. Common genetic variants influence human subcortical brain structures. Nature.

Holland, D., 2019a. GWAS-Causal-Effects-Extended-Model. https://github.com/dominicholland/GWAS_causalEffects_extendedModel.

Holland, D., 2019b. GWAS-Causal-Effects-Model. https://github.com/dominicholland/GWAS-Causal-Effects-Model.

Holland, D., Frei, O., Desikan, R., Fan, C.-C, Shadrin, A. A., Smeland, O. B., Sundar, V., Thompson, P., Andreassen, O. A., Dale, A. M., 2020. Beyond snp heritability: Polygenicity and discoverability of phenotypes estimated with a univariate gaussian mixture model. PLOS Genetics 16 (5), e1008612. URL https://doi.org/10.1371/journal.pgen.1008612

Jansen, I., Savage, J., Watanabe, K., Bryois, J., Williams, D., Steinberg, S., Sealock, J., Karlsson, I., Hagg, S., Athanasiu, L., et al., 2018. Genetic meta-analysis identifies 10 novel loci and functional pathways for alzheimer’s disease risk. bioRxiv, 258533.

Laird, N. M., Lange, C., 2010. The fundamentals of modern statistical genetics. Springer Science & Business Media.

Lambert, J.-C, Ibrahim-Verbaas, C. A., Harold, D., Naj, A. C., Sims, R., Bellenguez, C., Jun, G., DeStefano, A. L., Bis, J. C., Beecham, G. W., et al., 2013. Meta-analysis of 74,046 individuals identifies 11 new susceptibility loci for alzheimer’s disease. Nature genetics 45 (12), 1452–1458.

Lee, S. H., Yang, J., Chen, G.-B, Ripke, S., Stahl, E. A., Hultman, C. M., Sklar, P., Visscher, P. M., Sullivan, P. F., Goddard, M. E., et al., 2013. Estimation of snp heritability from dense genotype data. The American Journal of Human Genetics 93 (6), 1151–1155.

Li, N., Stephens, M., 2003. Modeling linkage disequilibrium and identifying recombination hotspots using single-nucleotide polymorphism data. Genetics 165 (4), 2213–2233.

Locke, A. E., Kahali, B., Berndt, S. I., Justice, A. E., Pers, T. H., Day, F. R., Powell, C., Vedantam, S., Buchkovich, M. L., Yang, J., et al., 2015. Genetic studies of body mass index yield new insights for obesity biology. Nature 518 (7538), 197.

Mehta, P., Kaye, W., Raymond, J. e. a., 2018. Prevalence of amyotrophic lateral sclerosis 2014 united states. MMWR Morb Mortal Wkly Rep 67, 216–218.

Merikangas, K. R., Jin, R., He, J.-P, Kessler, R. C., Lee, S., Sampson, N. A., Viana, M. C., Andrade, L. H., Hu, C., Karam, E. G., et al., 2011. Prevalence and correlates of bipolar spectrum disorder in the world mental health survey initiative. Archives of general psychiatry 68 (3), 241–251.

Nikpay, M., Goel, A., Won, H.-H, Hall, L. M., Willenborg, C., Kanoni, S., Saleheen, D., Kyriakou, T., Nelson, C. P., Hopewell, J. C., et al., 2015. A comprehensive 1000 genomes–based genomewide association meta-analysis of coronary artery disease. Nature genetics 47 (10), 1121.

O’Connor, L. J., Schoech, A. P., Hormozdiari, F., Gazal, S., Patterson, N., Price, A. L., 2019. Extreme polygenicity of complex traits is explained by negative selection. The American Journal of Human Genetics 105 (3), 456–476.

Okbay, A., Beauchamp, J. P., Fontana, M. A., Lee, J. J., Pers, T. H., Rietveld, C. A., Turley, P., Chen, G.-B, Emilsson, V., Meddens, S. F. W., et al., 2016. Genome-wide association study identifies 74 loci associated with educational attainment. Nature 533 (7604), 539–542.

Park, J. H., Gail, M. H., Weinberg, C. R., Carroll, R. J., Chung, C., Wang, Z., Chanock, S. J., Fraumeni, J. F., Chatterjee, N., Nov 2011. Distribution of allele frequencies and effect sizes and their interrelationships for common genetic susceptibility variants. Proc. Natl. Acad. Sci. U.S.A. 108 (44), 18026–18031.

Plassman, B. L., Langa, K. M., Fisher, G. G., Heeringa, S. G., Weir, R., Ofstedal, M. B., Burke, J. R., Hurd, M. D., Potter, G. G., Rodgers, W. L., et al., 2007. Prevalence of dementia in the united states: the aging, demographics, and memory study. Neuroepidemiology 29 (1-2), 125–132.

Pritchard, J. K., Cox, N. J., 2002. The allelic architecture of human disease genes: common disease–common variant or not? Human molecular genetics 11 (20), 2417–2423.

Pritchard, J. K., Przeworski, M., 2001. Linkage disequilibrium in humans: models and data. The American Journal of Human Genetics 69 (1), 1–14.

Sanchis-Gomar, F., Perez-Quilis, C., Leischik, R., Lucia, A., 2016. Epidemiology of coronary heart disease and acute coronary syndrome. Annals of translational medicine 4 (13).

Savage, J. E., Jansen, P. R., Stringer, S., Watanabe, K., Bryois, J., De Leeuw, C. A., Nagel, M., Awasthi, S., Barr, P. B., Coleman, J. R., et al., 2018. Genome-wide association meta-analysis in 269,867 individuals identifies new genetic and functional links to intelligence. Nature genetics 50 (7), 912–919.

Schizophrenia Working Group of the PGC, Jul 2014. Biological insights from 108 schizophrenia-associated genetic loci. Nature 511 (7510), 421–427.

Schoech, A. P., Jordan, D. M., Loh, P.-R, Gazal, S., O’Connor, L. J., Balick, D. J., Palamara, P. F., Finucane, H. K., Sunyaev, S. R., Price, A. L., 2019. Quantification of frequency-dependent genetic architectures in 25 uk biobank traits reveals action of negative selection. Nature communications 10 (1), 790.

Schork, A. J., Thompson, W. K., Pham, P., Torkamani, A., Roddey, C., Sullivan, P. F., Kelsoe, J. R., O’Donovan, M. C., Furberg, H., Schork, N. J., et al., 2013. All snps are not created equal: genome-wide association studies reveal a consistent pattern of enrichment among functionally annotated snps. PLoS genetics 9 (4), e1003449.

Shadrin, A. A., Frei, O., Smeland, O. B., Bettella, F., OConnell, S., Gani, O., Bahrami, S., Uggen, T. K., Djurovic, S., Holland, D., et al., 2019. Annotation-informed causal mixture modeling (ai-mixer) reveals phenotype-specific differences in polygenicity and effect size distribution across functional annotation categories. bioRxiv, 772202.

Sniekers, S., Stringer, S., Watanabe, K., Jansen, P. R., Coleman, J. R., Krapohl, E., Taskesen, E., Hammerschlag, A. R., Okbay, A., Zabaneh, D., et al., 2017. Genome-wide association meta-analysis of 78,308 individuals identifies new loci and genes influencing human intelligence. Nature genetics 49 (7), 1107.

Sohail, M., Maier, R. M., Ganna, A., Bloemendal, A., Martin, A. R., Turchin, M. C., Chiang, C. W., Hirschhorn, J., Daly, M. J., Patterson, N., et al., 2019. Polygenic adaptation on height is overestimated due to uncorrected stratification in genome-wide association studies. eLife 8, e39702.

Speed, D., Balding, D. J., 2019. Sumher better estimates the snp heritability of complex traits from summary statistics. Nature genetics 51 (2), 277.

Speed, D., Cai, N., Johnson, M. R., Nejentsev, S., Balding, D. J., Consortium, U., et al., 2017. Reevaluation of snp heritability in complex human traits. Nature genetics 49 (7), 986.

Speed, D., Hemani, G., Johnson, M. R., Balding, D. J., 2012. Improved heritability estimation from genome-wide snps. The American Journal of Human Genetics 91 (6), 1011–1021.

Spencer, C. C., Su, Z., Donnelly, P., Marchini, J., 2009. Designing genome-wide association studies: sample size, power, imputation, and the choice of genotyping chip. PLoS Genet 5 (5), e1000477.

Stahl, E. A., Breen, G., Forstner, A. J., McQuillin, A., Ripke, S., Trubetskoy, V., Mattheisen, M., Wang, Y., Coleman, J. R., Gaspar, H. A., et al., 2019. Genome-wide association study identifies 30 loci associated with bipolar disorder. Nature genetics, 1.

Su, Z., Marchini, J., Donnelly, P., 2011. Hapgen2: simulation of multiple disease snps. Bioinformatics 27 (16), 2304–2305.

Sveinbjornsson, G., Albrechtsen, A., Zink, F., Gudjonsson, S. A., Oddson, A., Másson, G., Holm, H., Kong, A., Thorsteinsdottir, U., Sulem, P., et al., 2016. Weighting sequence variants based on their annotation increases power of whole-genome association studies. Nature genetics.

Van Blitterswijk, M., DeJesus-Hernandez, M., Rademakers, R., 2012. How do c9orf72 repeat expansions cause amyotrophic lateral sclerosis and frontotemporal dementia: can we learn from other non-coding repeat expansion disorders? Current opinion in neurology 25 (6), 689–700.

Van Rheenen, W., Shatunov, A., Dekker, A. M., McLaughlin, R. L., Diekstra, F. P., Pulit, S. L., Van Der Spek, R. A., Võsa, U., De Jong, S., Robinson, M. R., et al., 2016. Genome-wide association analyses identify new risk variants and the genetic architecture of amyotrophic lateral sclerosis. Nature genetics 48 (9), 1043.

Willer, C. J., Schmidt, E. M., Sengupta, S., Peloso, G. M., Gustafsson, S., Kanoni, S., Ganna, A., Chen, J., Buchkovich, M. L., Mora, S., et al., 2013. Discovery and refinement of loci associated with lipid levels. Nature genetics 45 (11), 1274.

Wingo, T. S., Cutler, D. J., Yarab, N., Kelly, C. M., Glass, J. D., 2011. The heritability of amyotrophic lateral sclerosis in a clinically ascertained united states research registry. PLoS One 6 (11), e27985.

Wood, A. R., Esko, T., Yang, J., Vedantam, S., Pers, T. H., Gustafsson, S., Chu, A. Y., Estrada, K., Luan, J., Kutalik, Z., et al., 2014. Defining the role of common variation in the genomic and biological architecture of adult human height. Nature genetics 46 (11), 1173–1186.

Wray, N. R., Ripke, S., Mattheisen, M., Trzaskowski, M., Byrne, E. M., Abdellaoui, A., Adams, M. J., Agerbo, E., Air, T. M., Andlauer, T. M., et al., 2018. Genome-wide association analyses identify 44 risk variants and refine the genetic architecture of major depression. Nature genetics 50 (5), 668.

Yang, J., Benyamin, B., McEvoy, B. P., Gordon, S., Henders, A. K., Nyholt, D. R., Madden, P. A., Heath, A. C., Martin, N. G., Montgomery, G. W., Goddard, M. E., Visscher, P. M., Jul 2010. Common SNPs explain a large proportion of the heritability for human height. Nat. Genet. 42 (7), 565–569.

Yengo, L., Sidorenko, J., Kemper, K. E., Zheng, Z., Wood, A. R., Weedon, M. N., Frayling, T. M., Hirschhorn, J., Yang, J., Visscher, P. M., et al., 2018. Meta-analysis of genome-wide association studies for height and body mass index in 700000 individuals of european ancestry. Human molecular genetics 27 (20), 3641–3649.

Zeng, J., Vlaming, R., Wu, Y., Robinson, M. R., Lloyd-Jones, L. R., Yengo, L., Yap, C. X., Xue, A., Sidorenko, J., McRae, A. F., et al., 2018. Signatures of negative selection in the genetic architecture of human complex traits. Nature genetics 50 (5), 746.

Zhang, Y., Qi, G., Park, J.-H, Chatterjee, N., 2018. Estimation of complex effect-size distributions using summary-level statistics from genome-wide association studies across 32 complex traits. Nature genetics 50 (9), 1318.

Zhou, X., Carbonetto, P., Stephens, M., 2013. Polygenic modeling with bayesian sparse linear mixed models. PLoS genetics 9 (2), e1003264.

Zhu, X., Stephens, M., 2017. Bayesian large-scale multiple regression with summary statistics from genome-wide association studies. The annals of applied statistics 11 (3), 1561.

