## Supplementary figures. for "Linkage Disequilibrium and Heterozygosity Modulate the Genetic Architecture of Human Complex Phenotypes"

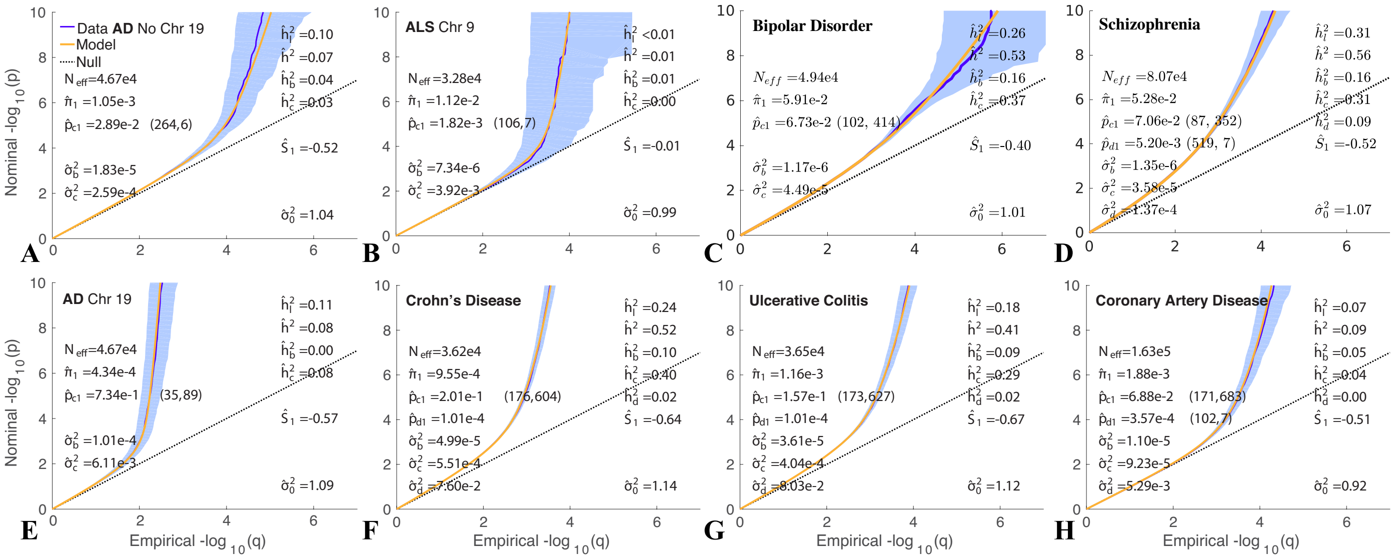


Figure S1: Blow-up of QQ plots of z-scores and model fits for qualitative phenotypes in Figure 2, focusing in on y-axis range $0 \leqslant -\log_{10}(p) \leqslant 10$.

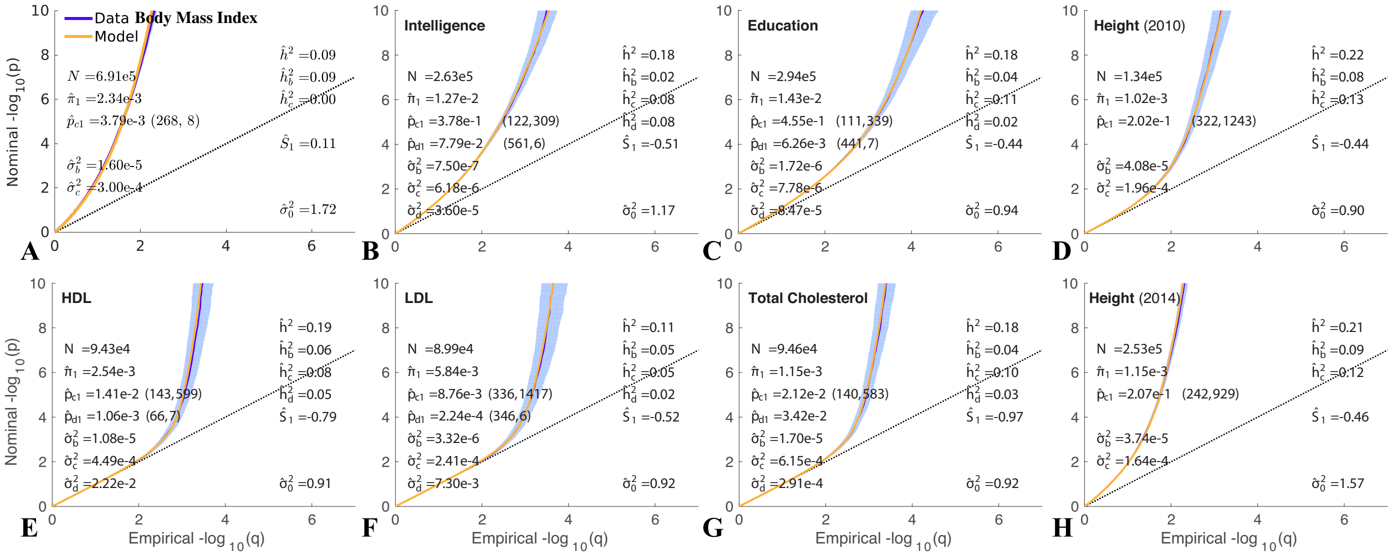


Figure S2: Blow-up of QQ plots of z-scores and model fits for quantitative phenotypes in Figure 3, focusing in on y-axis range $0 \leqslant -\log_{10}(p) \leqslant 10$.

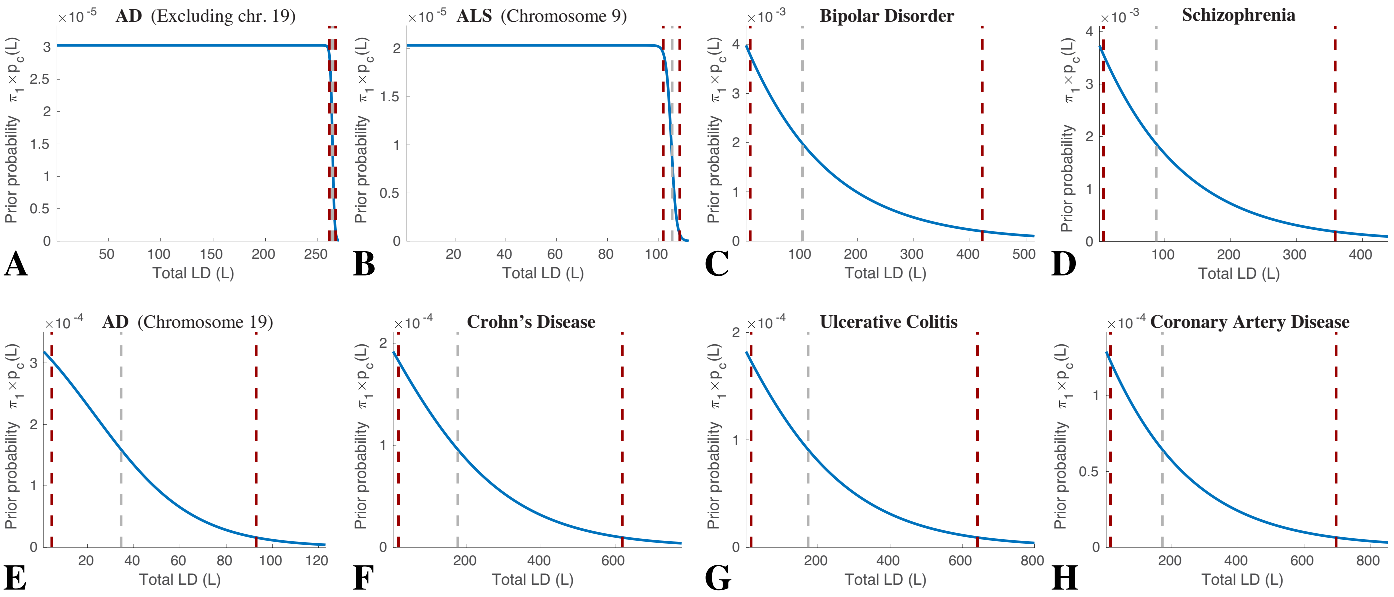


Figure S3: $\pi_1 \times p_c(L)$ for case-control phenotypes. See Eq. 5, Table 1, and Figure 2.

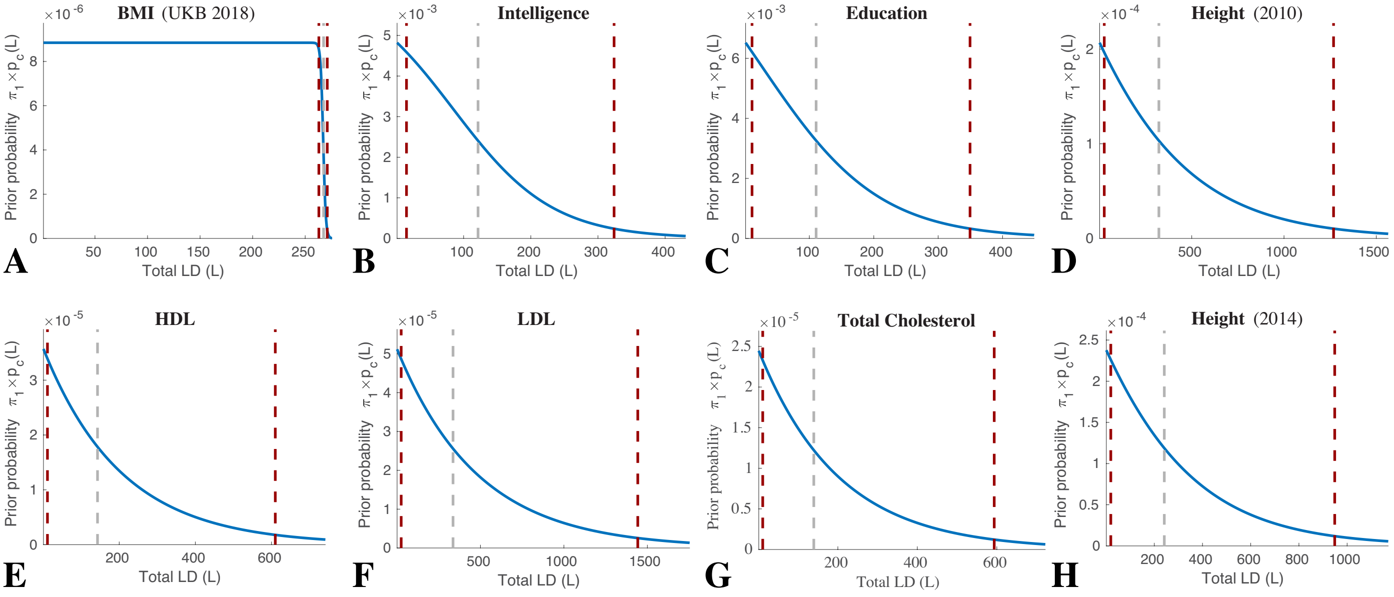


Figure S4: $\pi_1 \times p_c(L)$ for quantitative phenotypes. See Eq. 5, Table 1, and Figure 3.

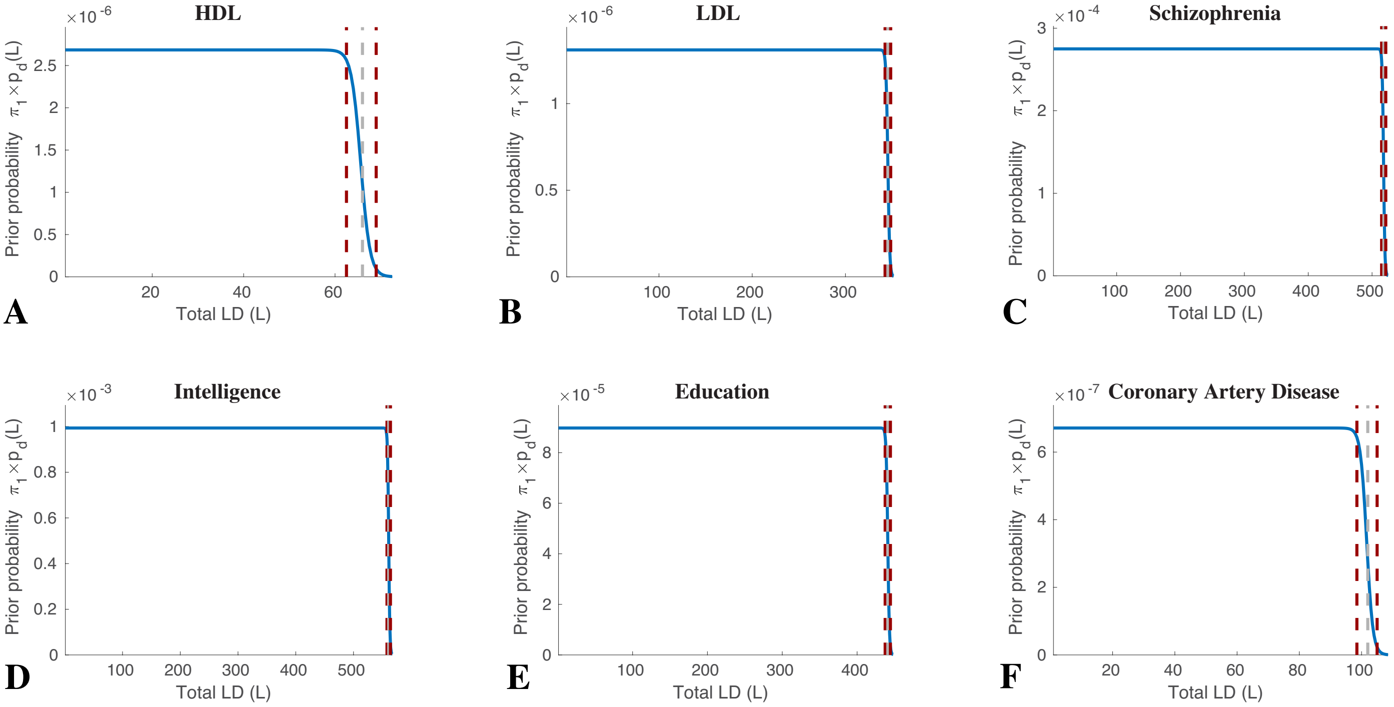


Figure S5: $\pi_1 \times p_d(L)$ for phenotypes. See Eq. 5, Table 1, and Figures 2 and 3. For CD, UC, and TC, $p_d(L)$ was a constant ($=p_d(1)$).

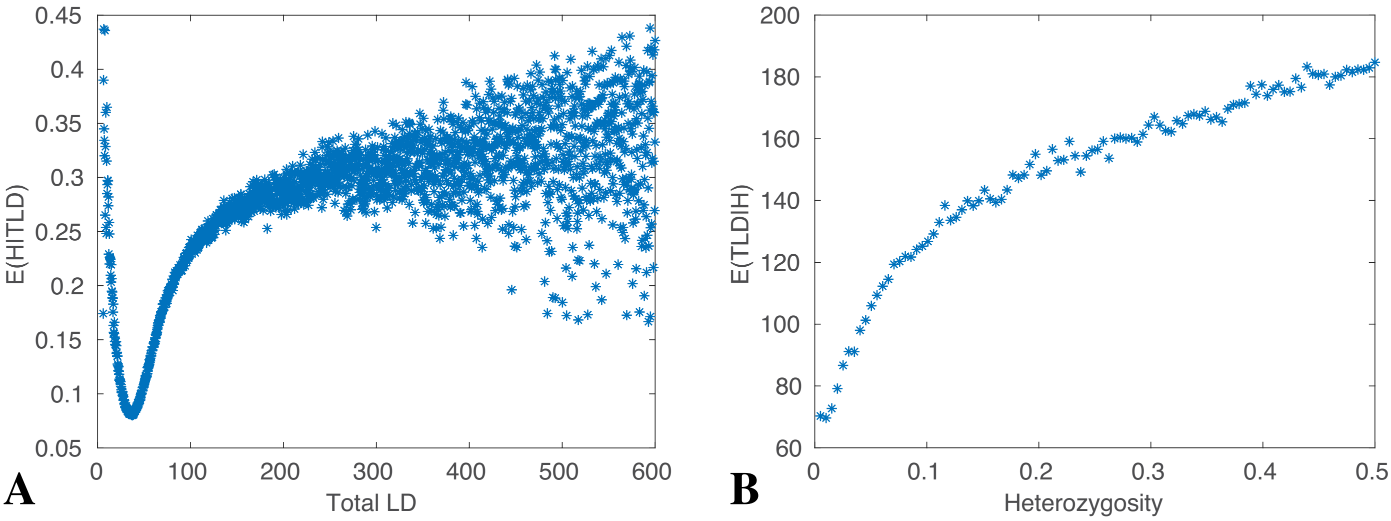


Figure S6: (A) Mean value of heterozygosity for given total LD (SNPs were binned based on TLD and the mean TLD for each bin plotted on the x-axis; the corresponding mean heterozygosity for SNPs in each bin was then plotted on the y-axis). (B) Mean value of total LD for given heterozygosity, H (SNPs were binned based on heterozygosity and the mean H for each bin plotted on the x-axis; the corresponding mean TLD for SNPs in each bin was then plotted on the y-axis). TLD and H calculated from 1000 Genomes phase 3 reference panel (Consortium et al., 2012, 2015).

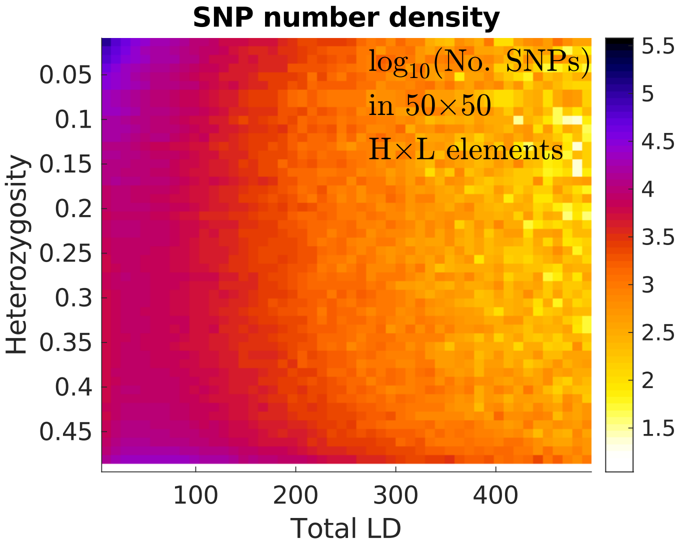


Figure S7: Number density of SNPs in the reference panel. Note that heterozygosity increases toward the bottom.

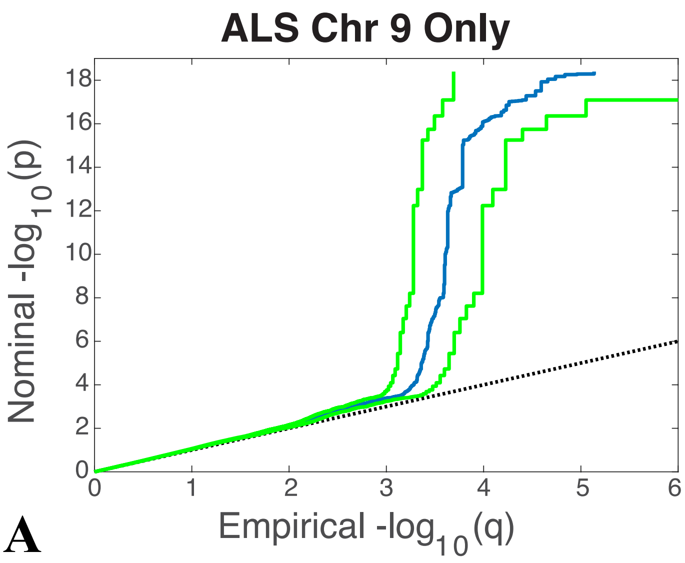


**A**

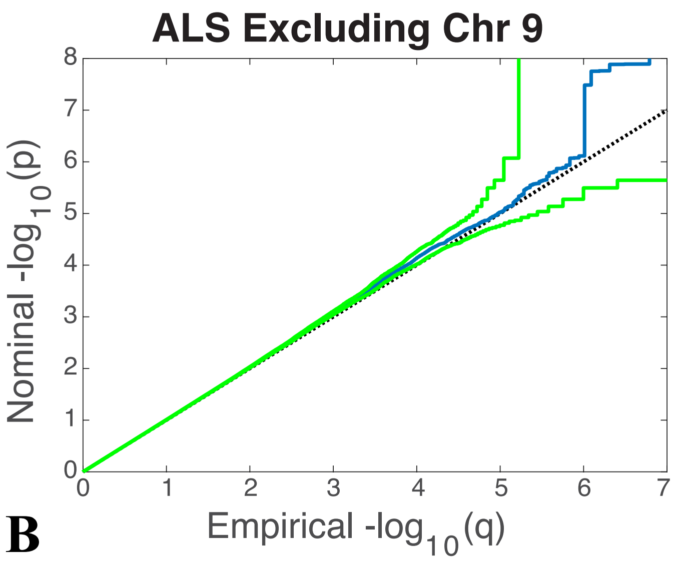


**B**

Figure S8: (A) QQ plot for amyotrophic lateral sclerosis (ALS) from chromosome 9 only, and (B) QQ plot from autosomal chromosomes excluding chromosome 9. Sample data in blue, 95% confidence intervals for population in green. All the significant GWAS signal is from chromosome 9. For these plots, there is no random pruning of SNPs, and total LD is not restricted to $< 600$. Compare Figure 2 B.

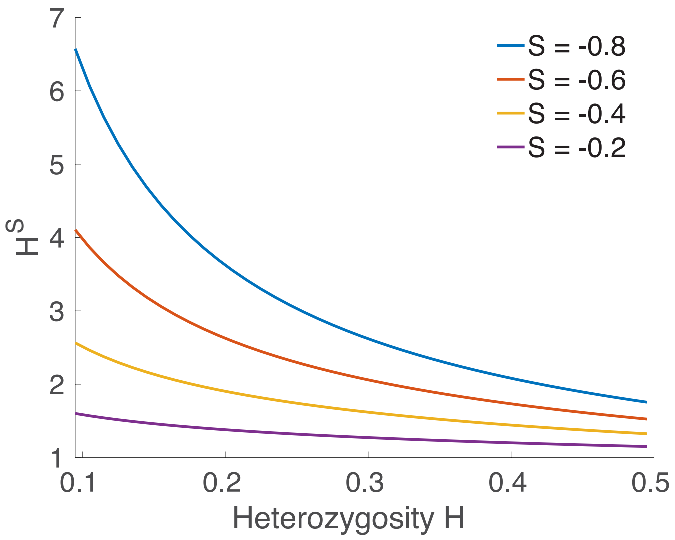


Figure S9: Examples of the factor $H^S$ multiplying $\sigma_c^2$, for $S = -0.8$, $-0.6$, , $-0.4$, and $-0.2$. See Eq. 5, Table 1, and Figures 2 and 3.

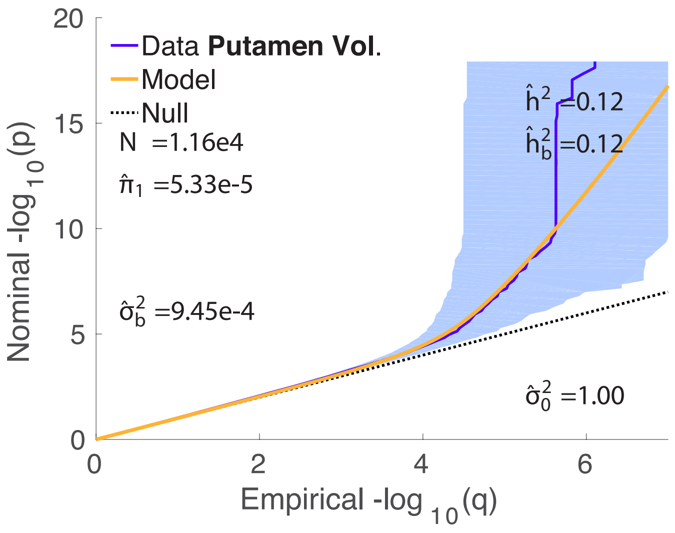


Figure S10: QQ plot for putamen volume (normalized by intracranial volume, $N = 11{,}598$ (Hibar et al., 2015)) and model fit. The 8-parameter C model gave a higher BIC (5777) value than the 3-parameter B model (5745), so we restrict analysis to the B model in this case – the same as in our earlier work (Holland et al., 2020). See Supplemental Material, Figure S30.

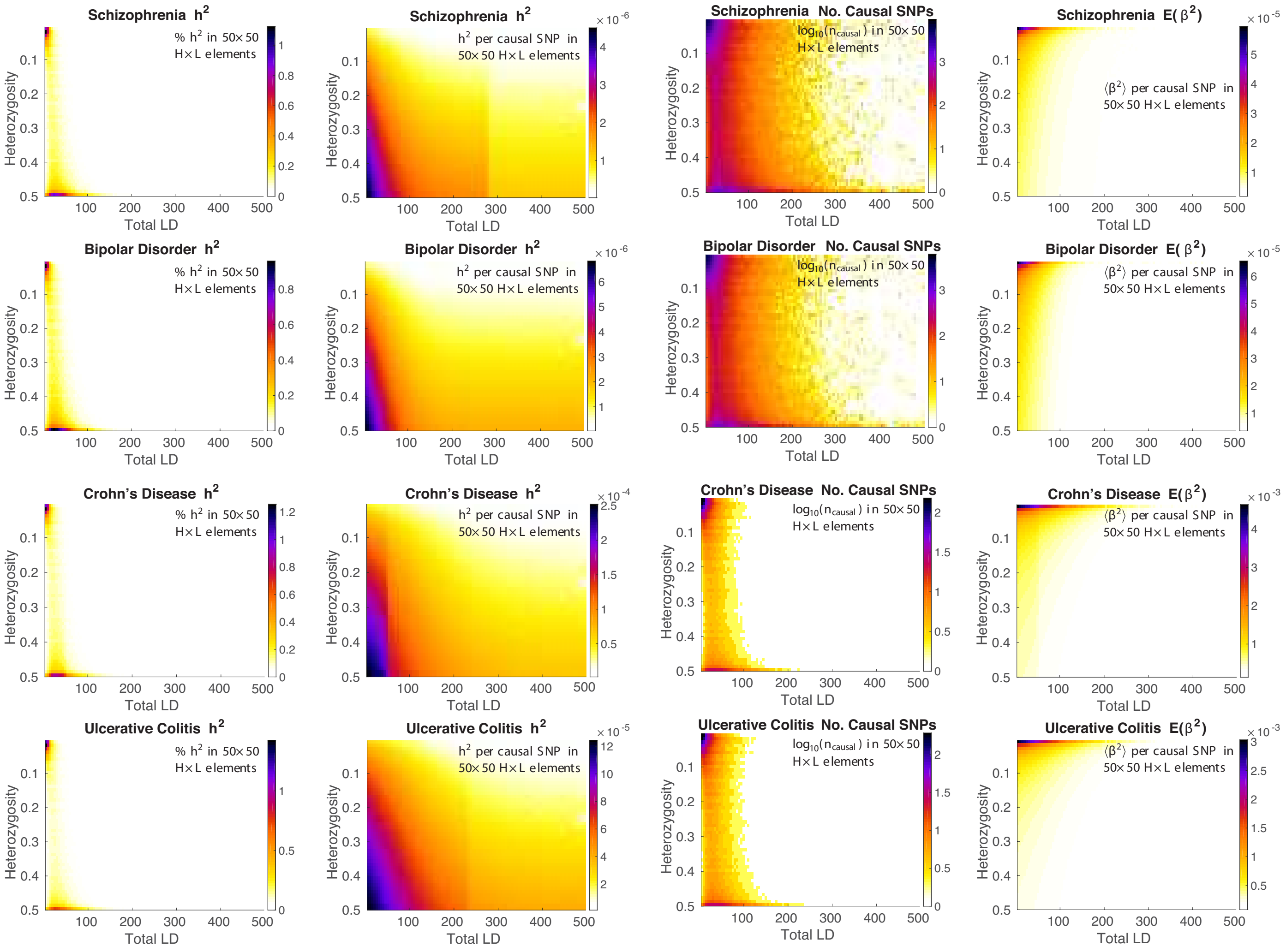


Figure S11: Model results in the top-to-bottom rows, respectively, for schizophrenia, bipolar disorder, Crohn's disease, and ulcerative colitis. The reference panel SNPs are binned with respect to both heterozygosity ($H$) and total LD ($L$) in a $50 \times 50$ grid for $0.02 \leqslant H \leqslant 0.5$ and $1 \leqslant L \leqslant 500$. Shown left to right are model estimates of: the percentage of heritability in each grid element; for each element, the average heritability per causal-SNP in the element; $\log_{10}$ of the number of causal SNPs in each element; and the expected $\beta^2$ for the element-wise causal SNPs. Note that $H$ increases from top to bottom.

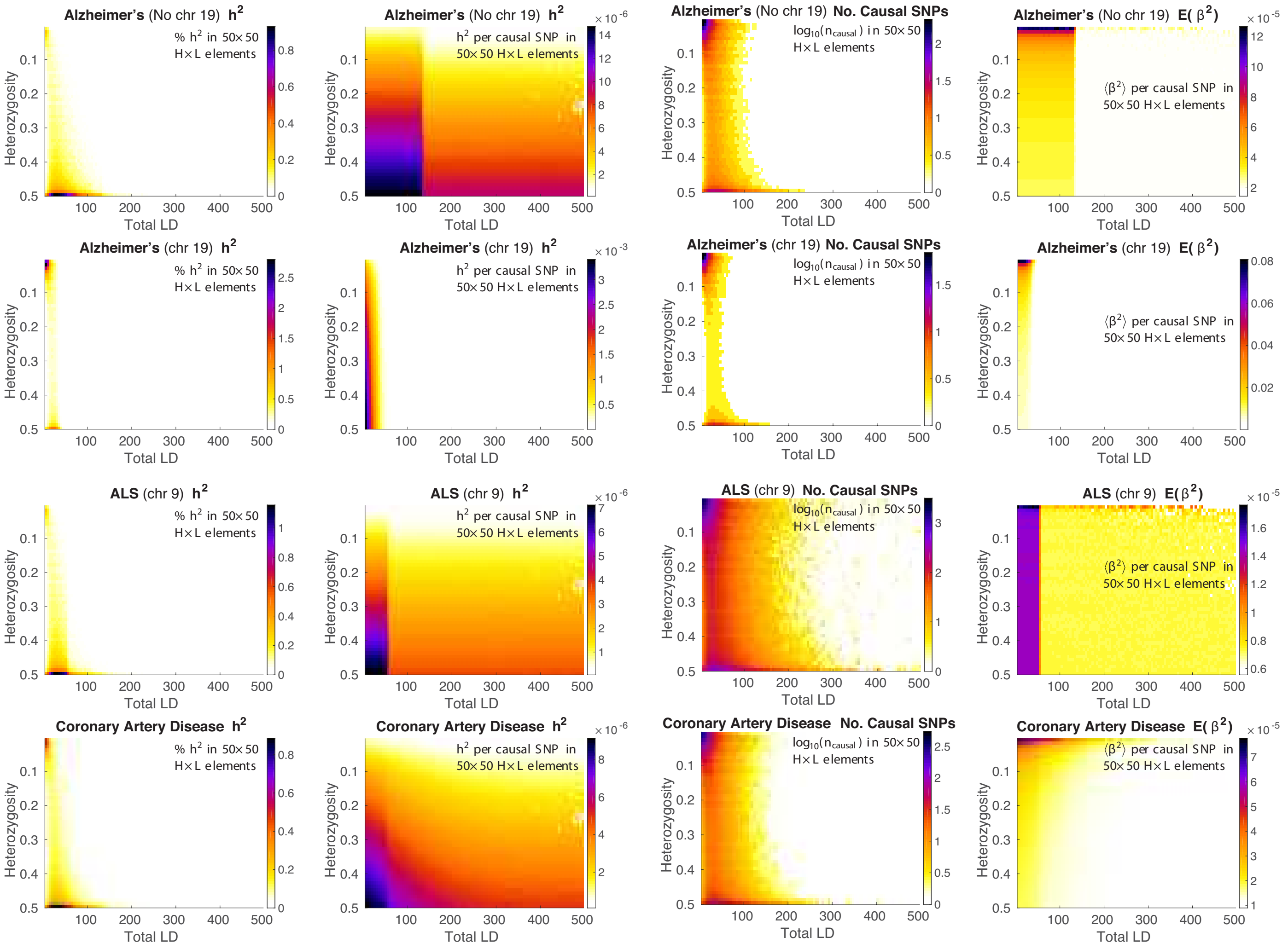


Figure S12: Model results in the top-to-bottom rows, respectively, for Alzheimer's disease (excluding chromosome 19), Alzheimer's disease (chromosome 19 only), amyotrophic lateral sclerosis (ALS), and coronary artery disease. See Supplemental Material, Figure S11 for details.

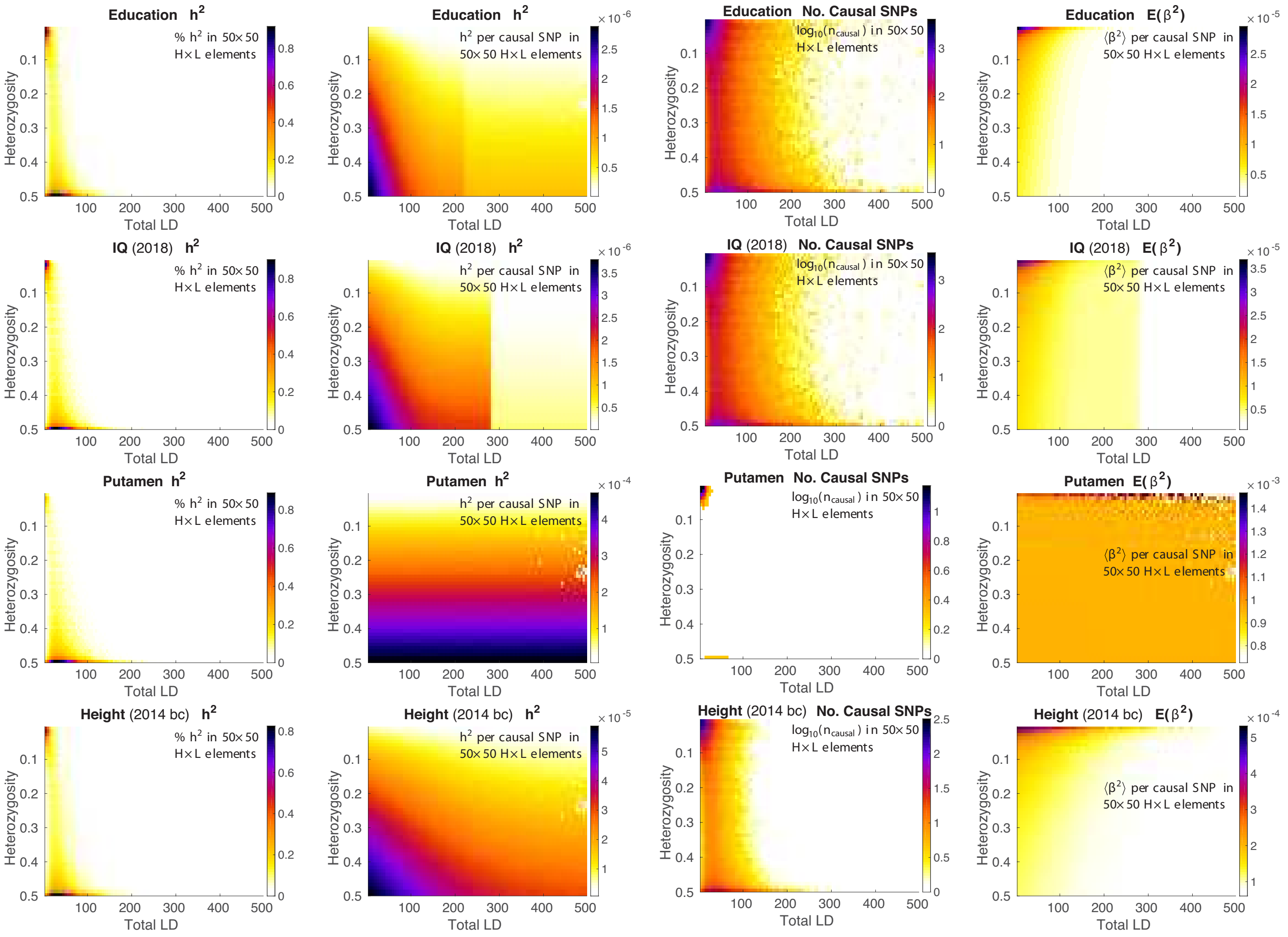


Figure S13: Model results in the top-to-bottom rows, respectively, for education, IQ, putamen volume (see also Supplemental Material, Figure S10), and height (2014). See Supplemental Material, Figure S11 for details.

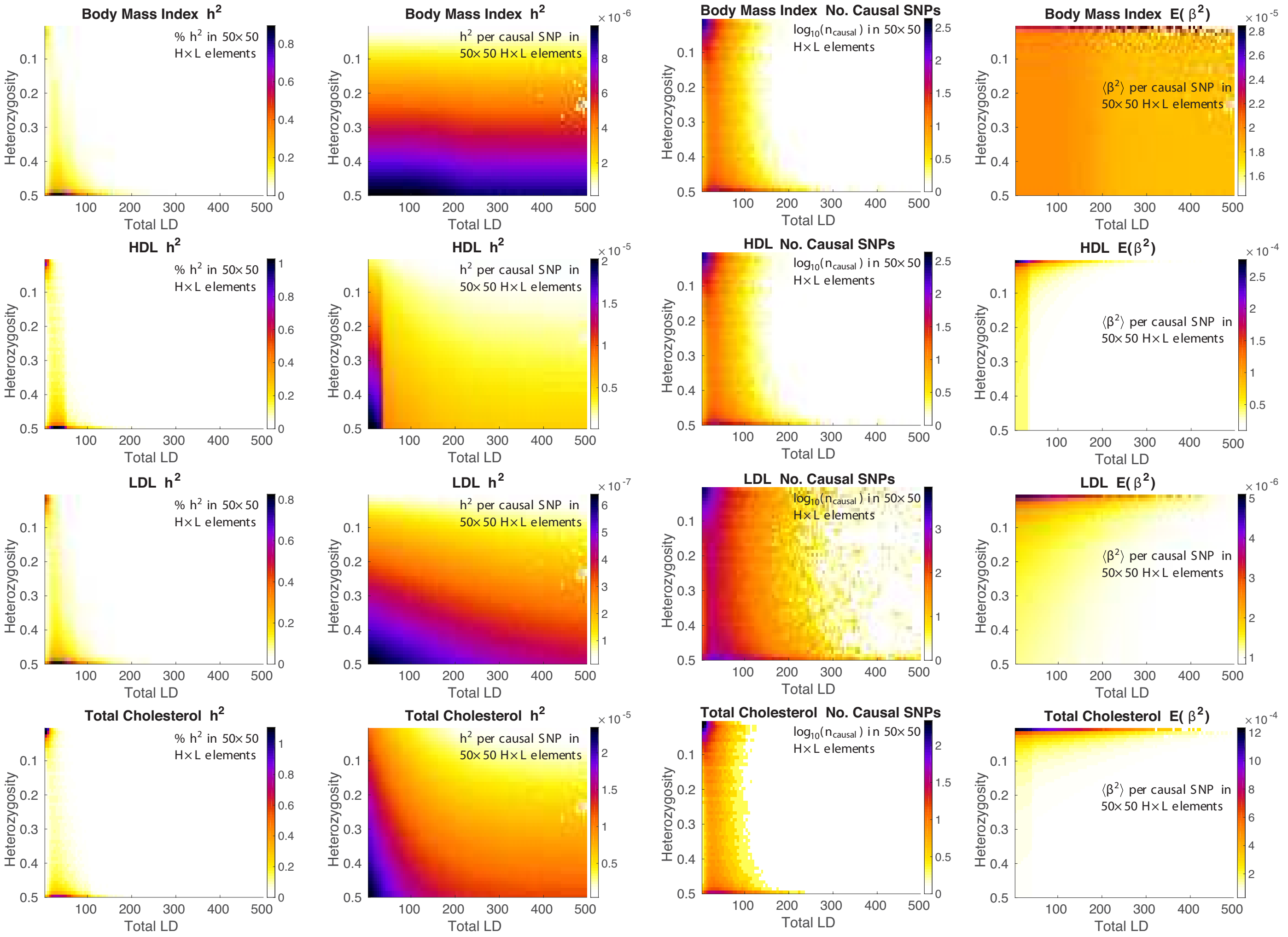


Figure S14: Model results in the top-to-bottom rows, respectively, for body mass index (BMI 2015), HDL, LDL, and total cholesterol. See Supplemental Material, Figure S11 for details.

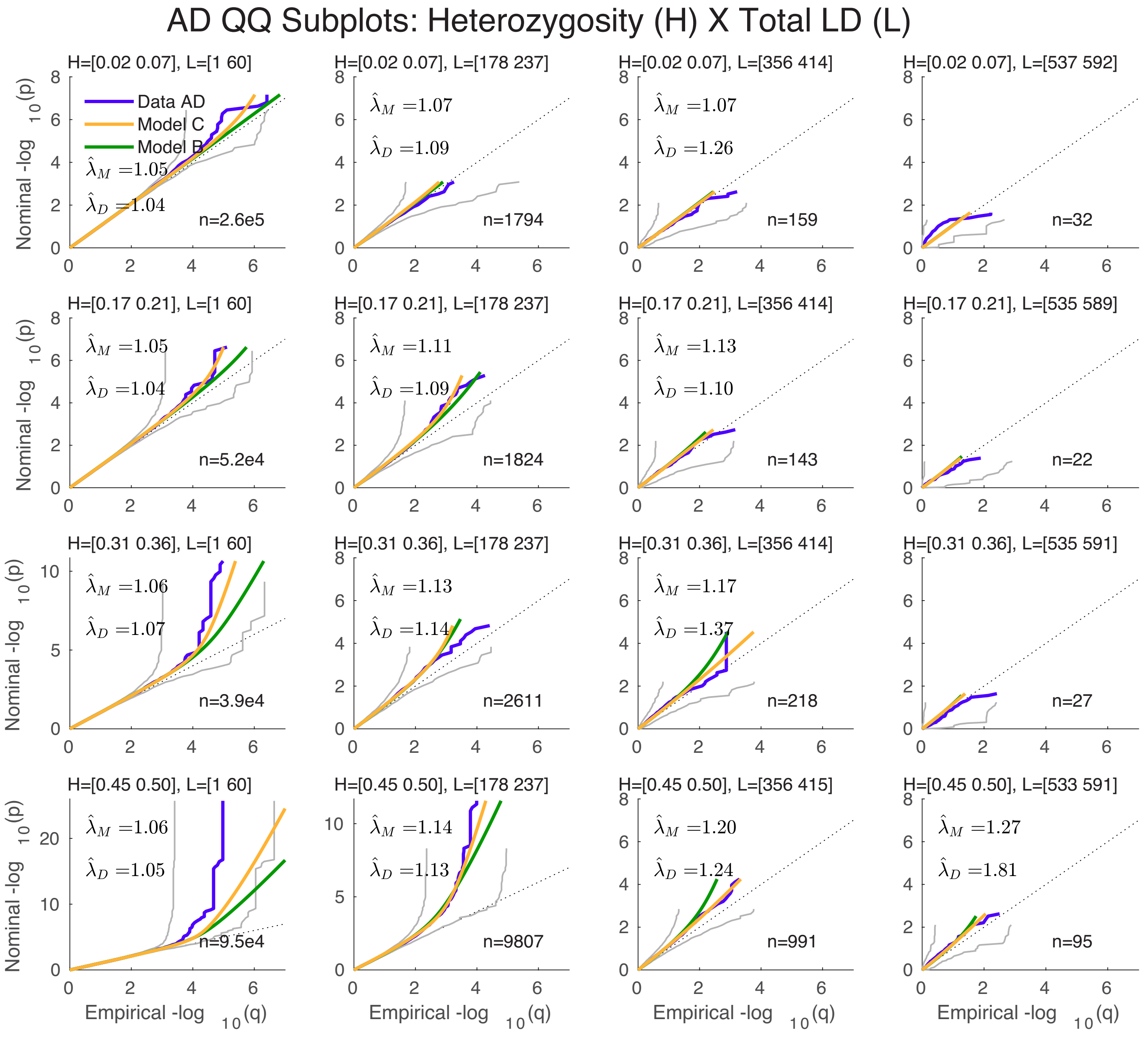


Figure S15: A 4×4 heterozygosity×TLD grid of QQ plots for Alzheimer's disease, excluding chromosome 19 (H increasing top to bottom, TLD increasing left to right, taking every second subplot from a full 10×10 grid). $n$ is the number of SNPs in each grid element. See Figure 2 (A) and Supplemental Material, Figure S16.

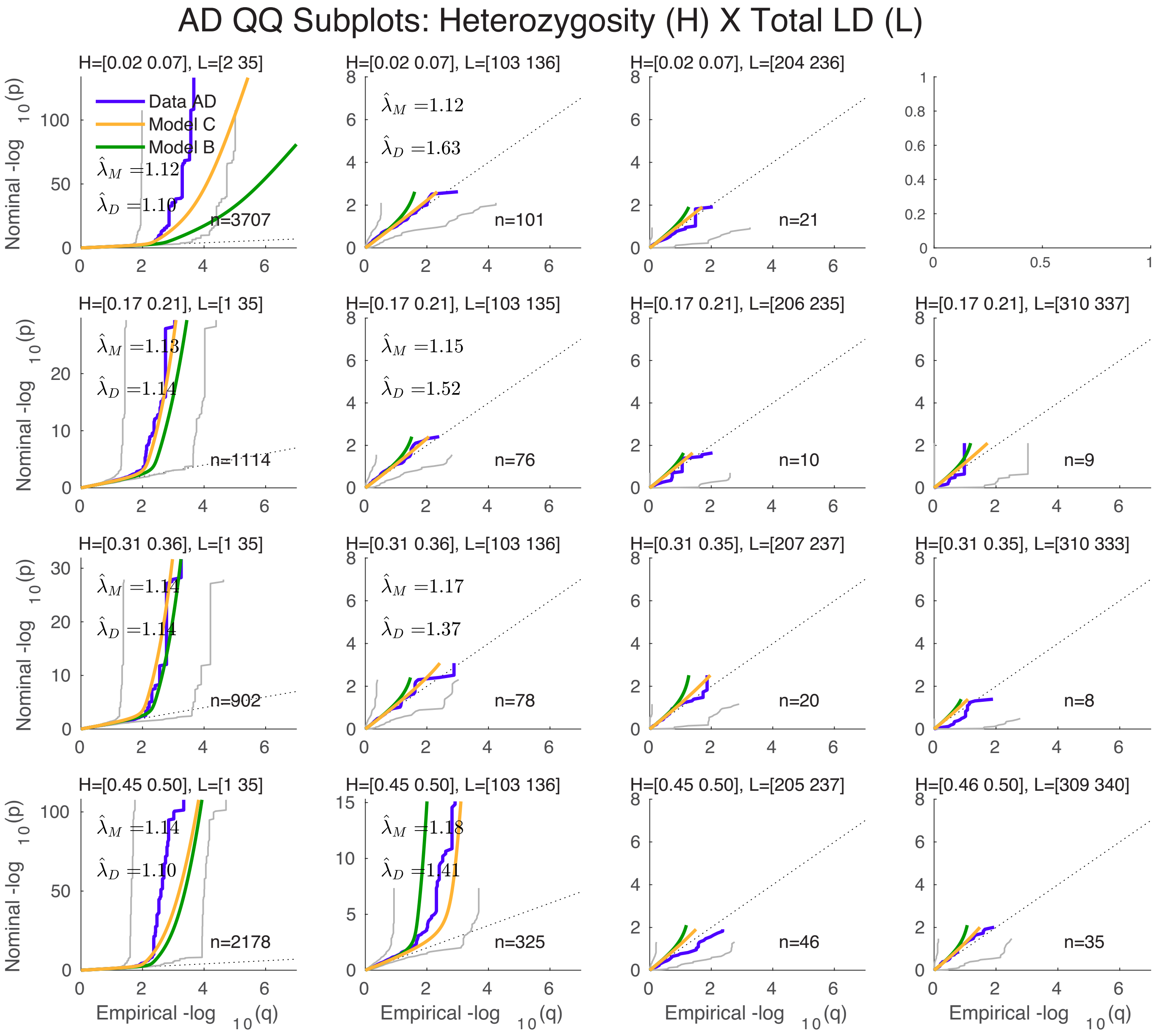


Figure S16: A 4×4 heterozygosity×TLD grid of QQ plots for Alzheimer's disease, restricted to chromosome 19. See Figure 2 (E) and Supplemental Material, Figure S15.

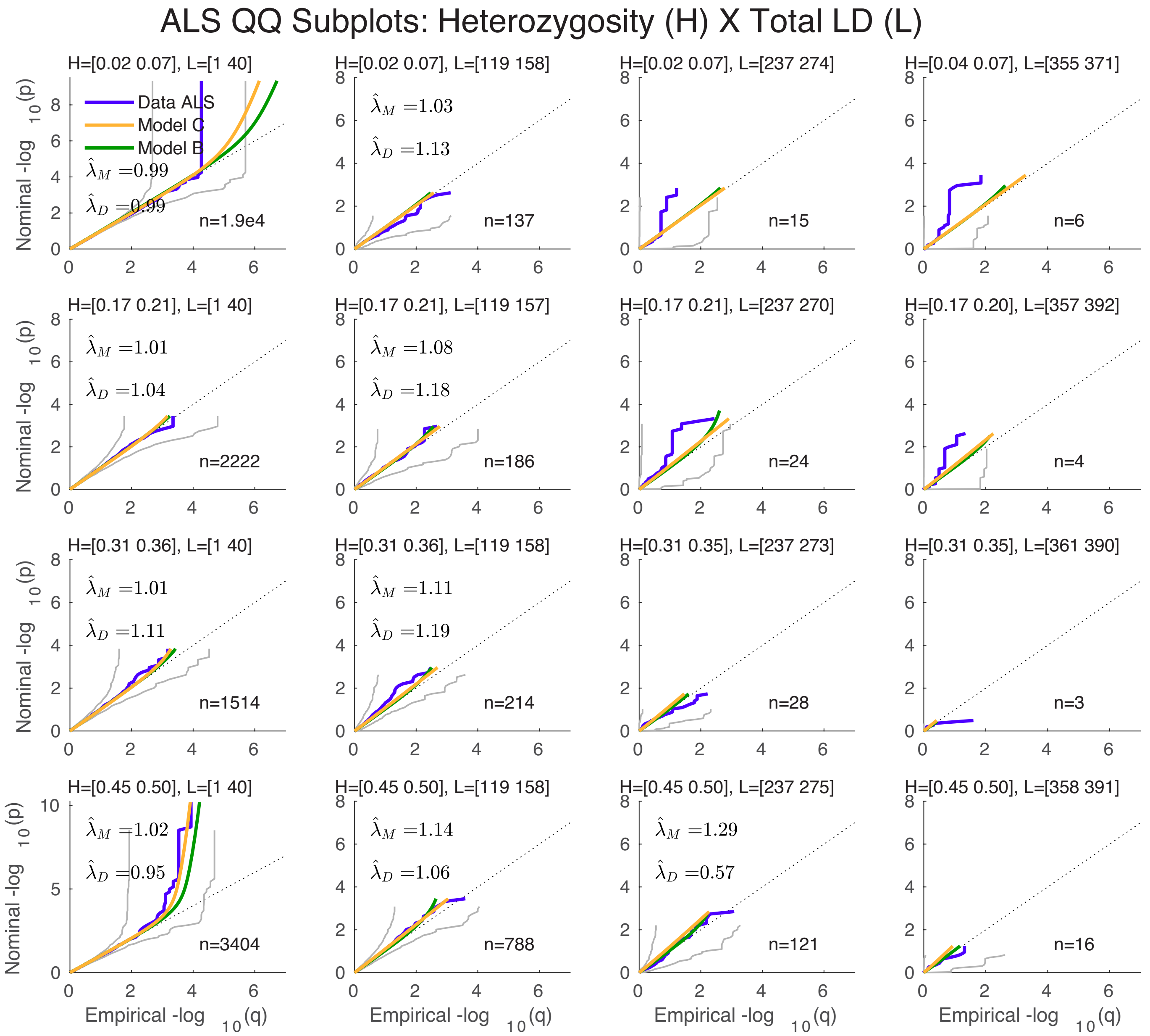


Figure S17: A 4×4 heterozygosity×TLD grid of QQ plots for amyotrophic lateral sclerosis, restricted to chromosome 9. See Figure 2 (B).

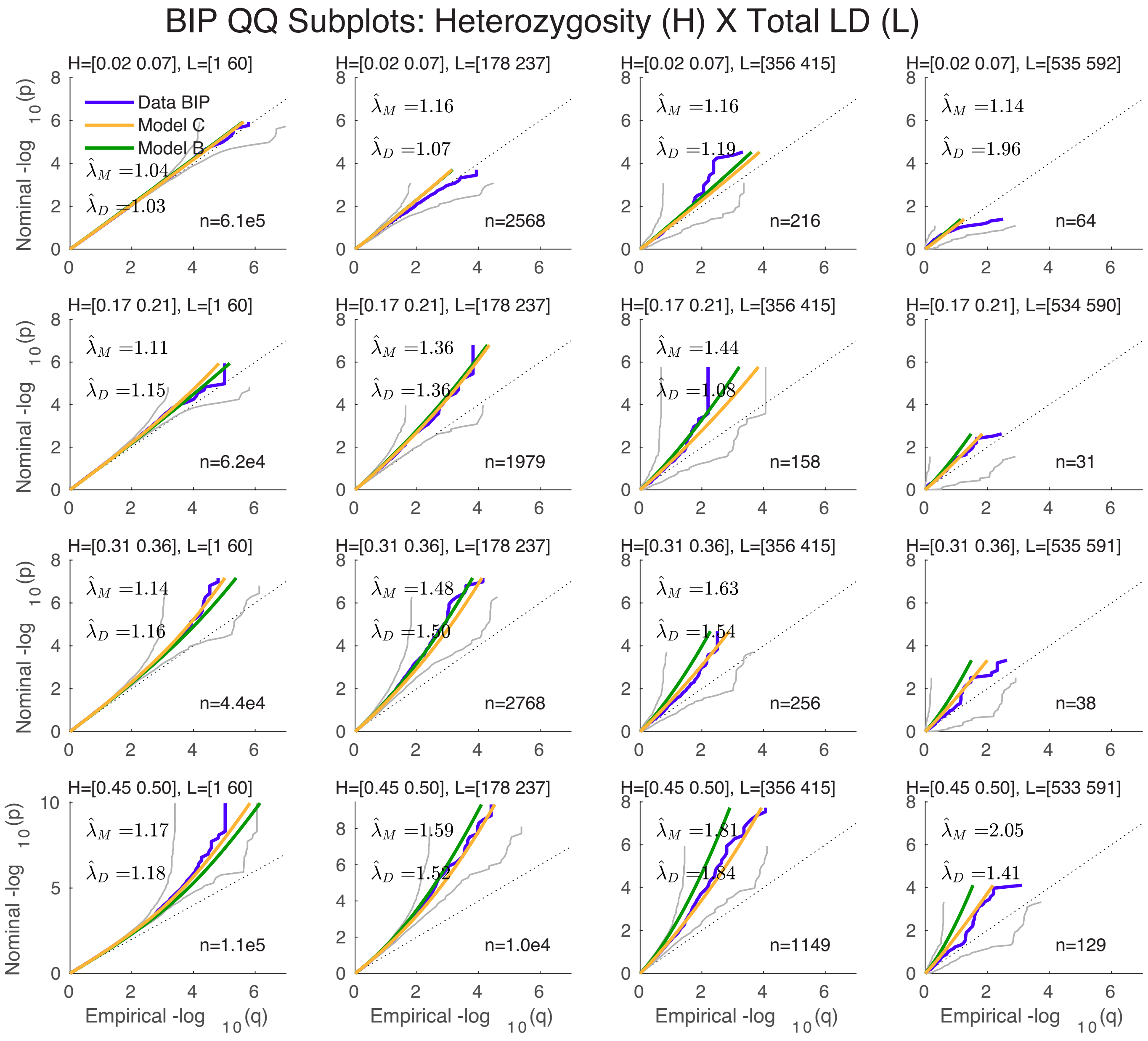


Figure S18: A 4×4 heterozygosity×TLD grid of QQ plots for bipolar disorder. See Figure 2 (C).

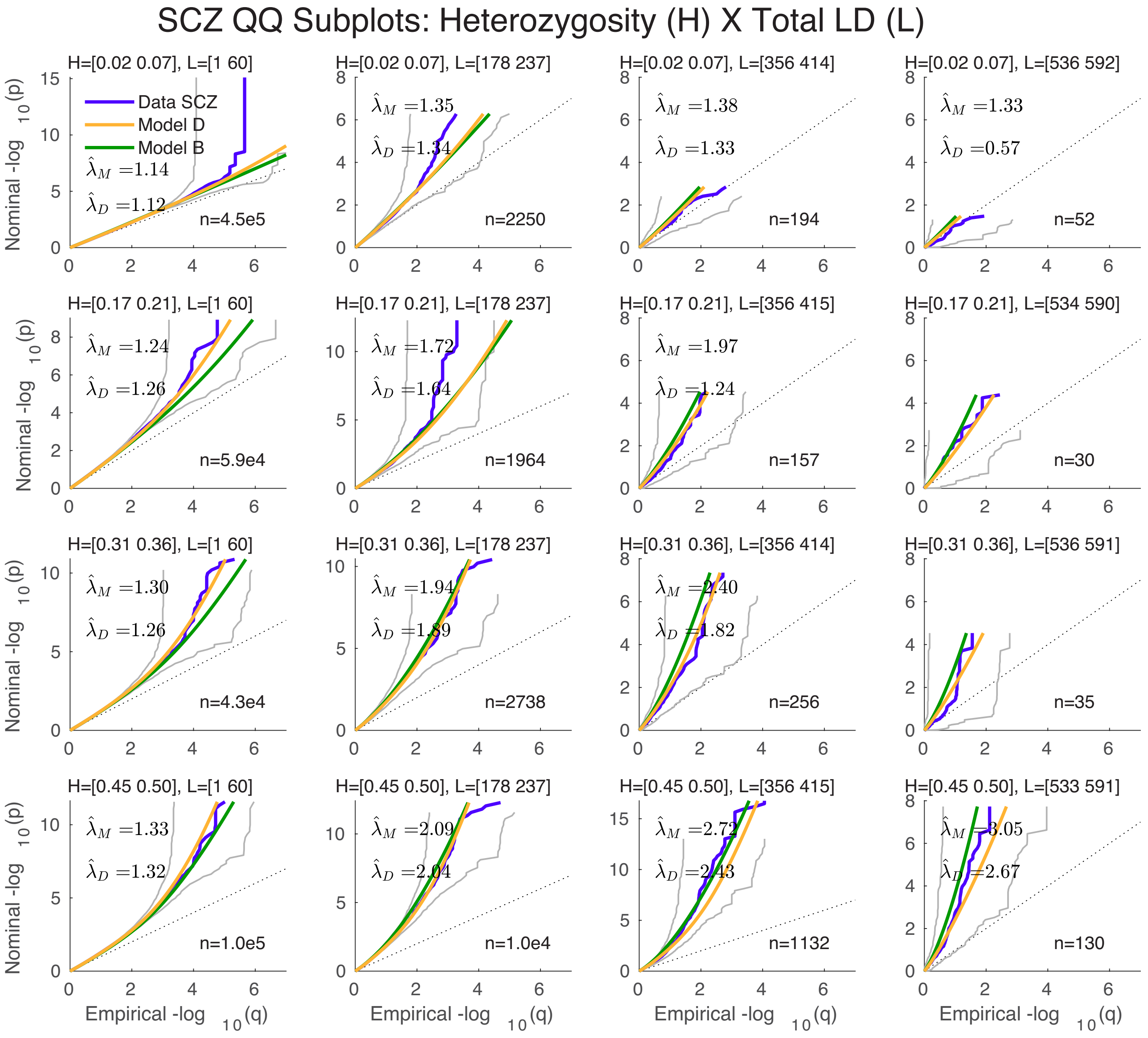


Figure S19: A 4×4 heterozygosity×TLD grid of QQ plots for schizophrenia (2014). See Figure 2 (D).

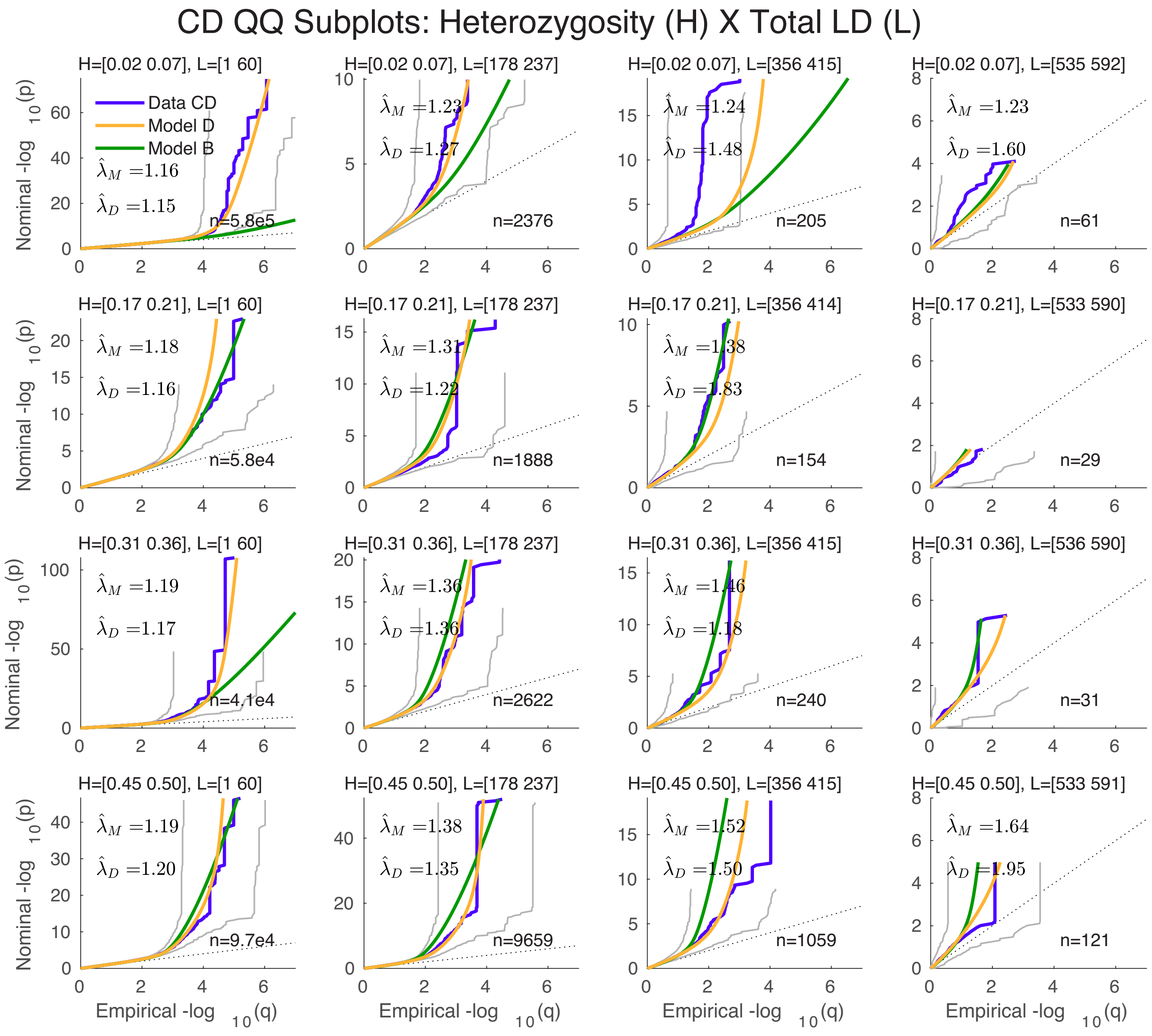


Figure S20: A 4×4 heterozygosity×TLD grid of QQ plots for Crohn's disease. See Figure 2 (F).

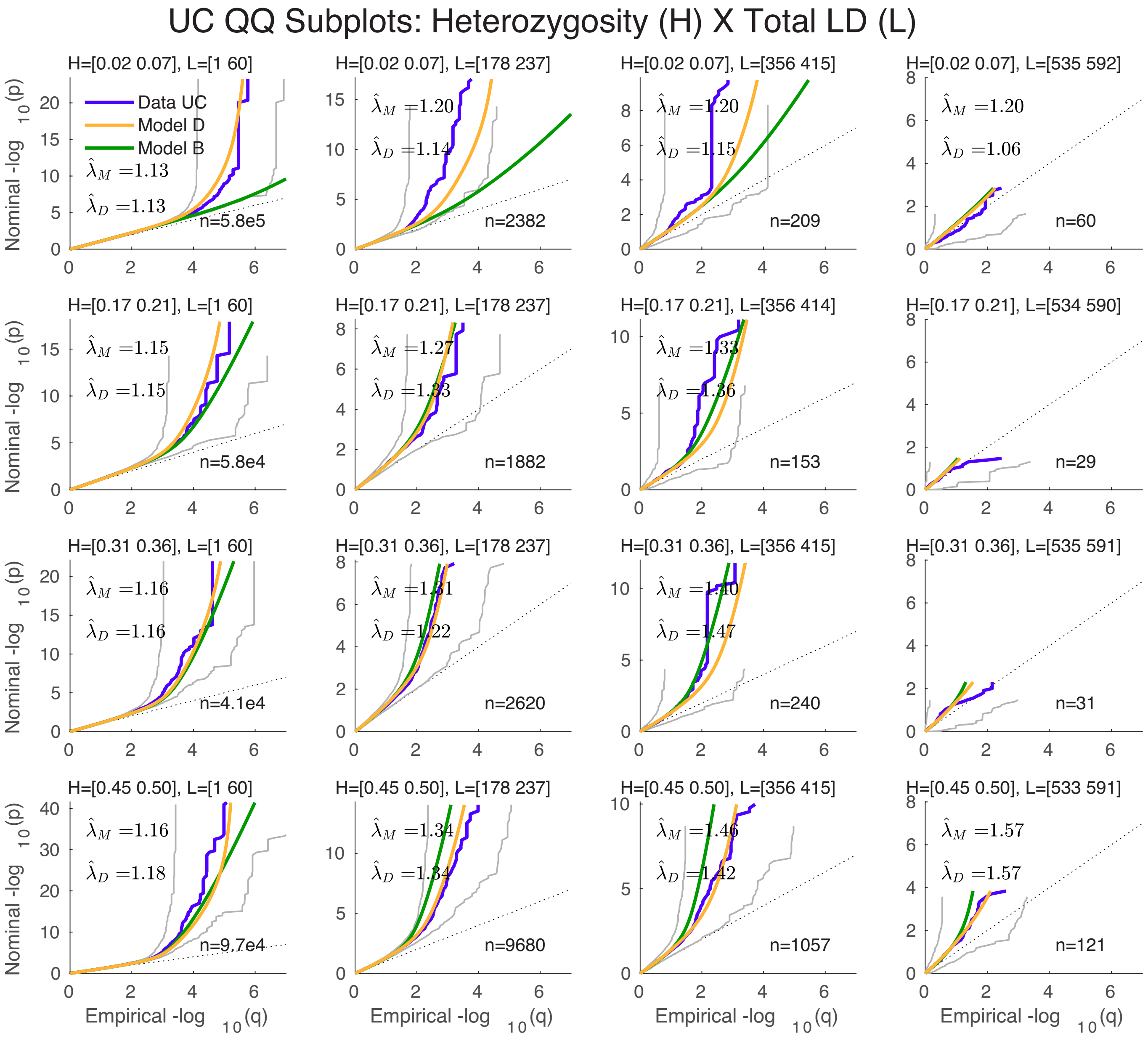


Figure S21: A 4×4 heterozygosity×TLD grid of QQ plots for ulcerative colitis. See Figure 2 (G).

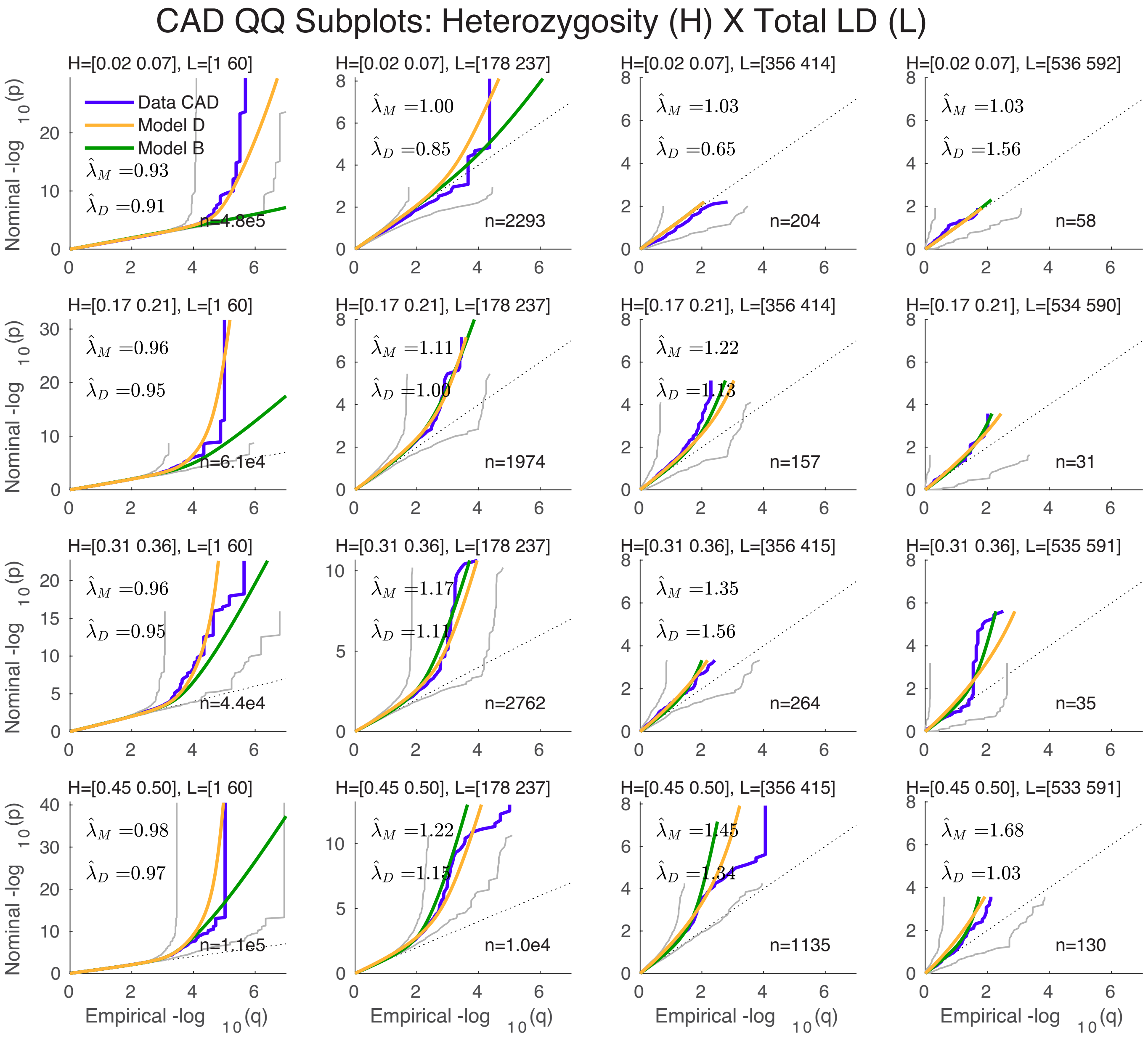


Figure S22: A 4×4 heterozygosity×TLD grid of QQ plots for coronary artery disease. See Figure 2 (H).

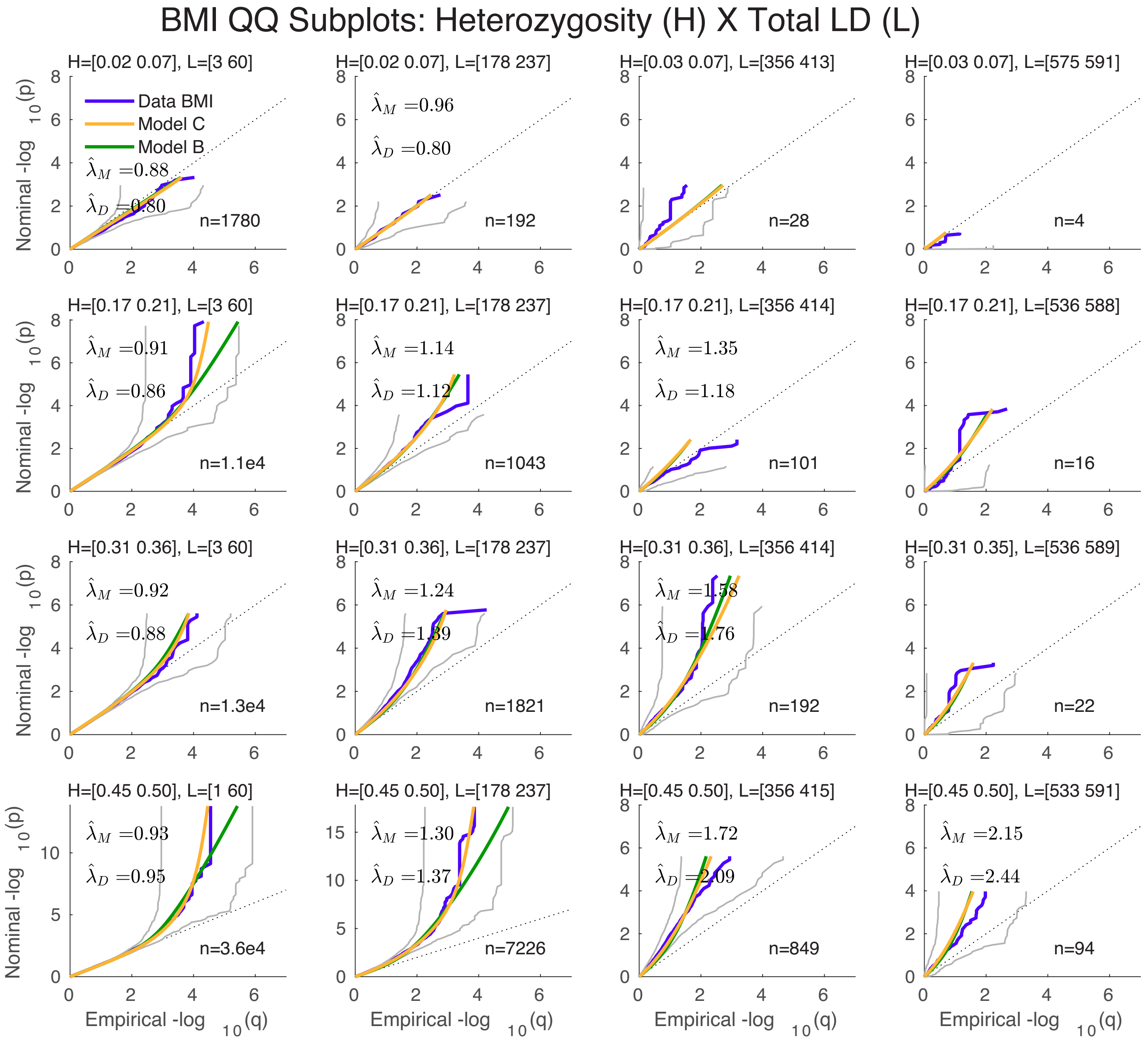


Figure S23: A 4×4 heterozygosity×TLD grid of QQ plots for body mass index (GIANT 2015 (Locke et al., 2015)). See Figure 3 (A), and Supplemental Material, Figures S24 and S37.

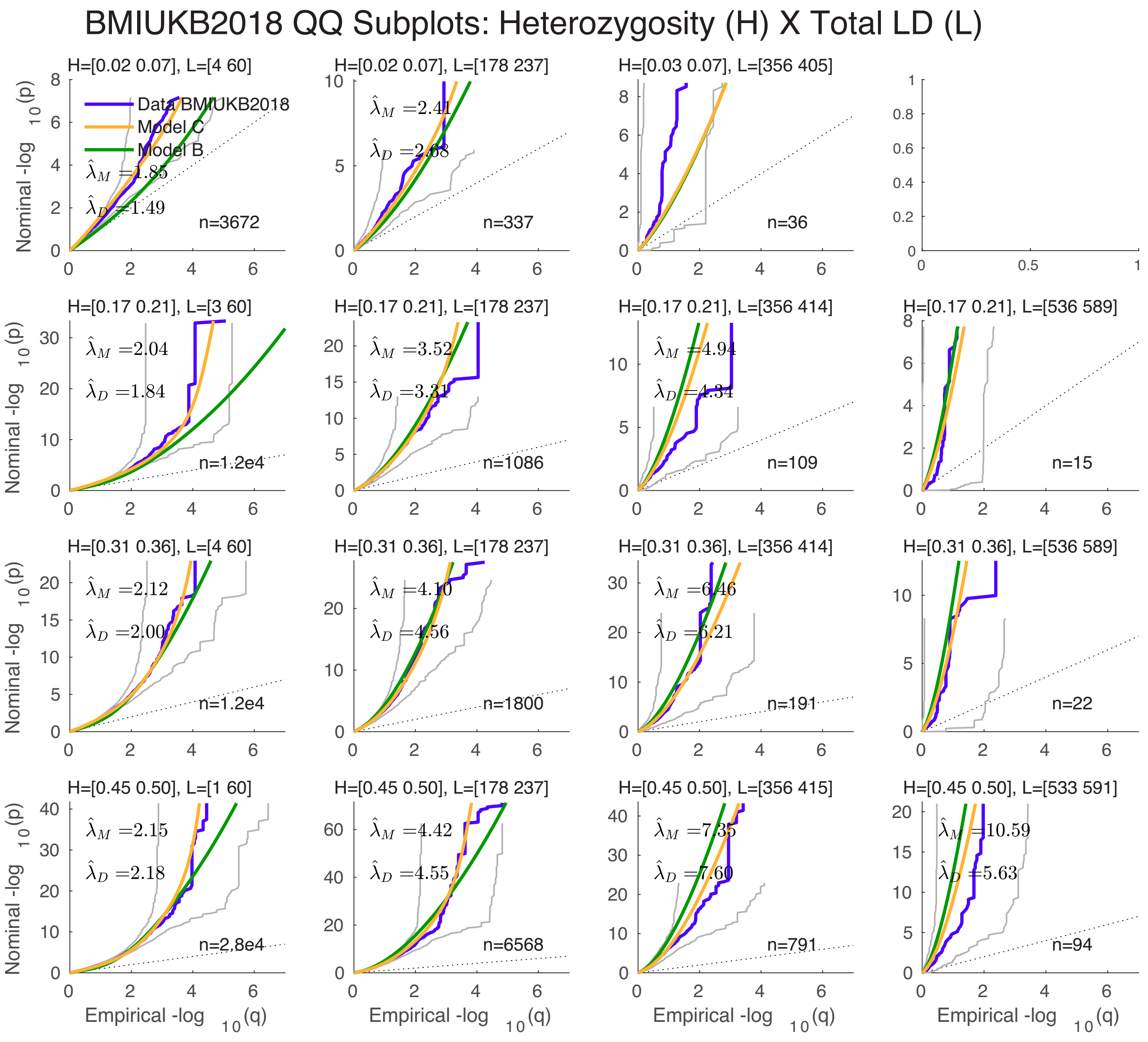


Figure S24: A 4×4 heterozygosity×TLD grid of QQ plots for body mass index (GIANT-UKB 2018 (Yengo et al., 2018)). See Figure 3 (A), and Supplemental Material, Figure S23 and S37.

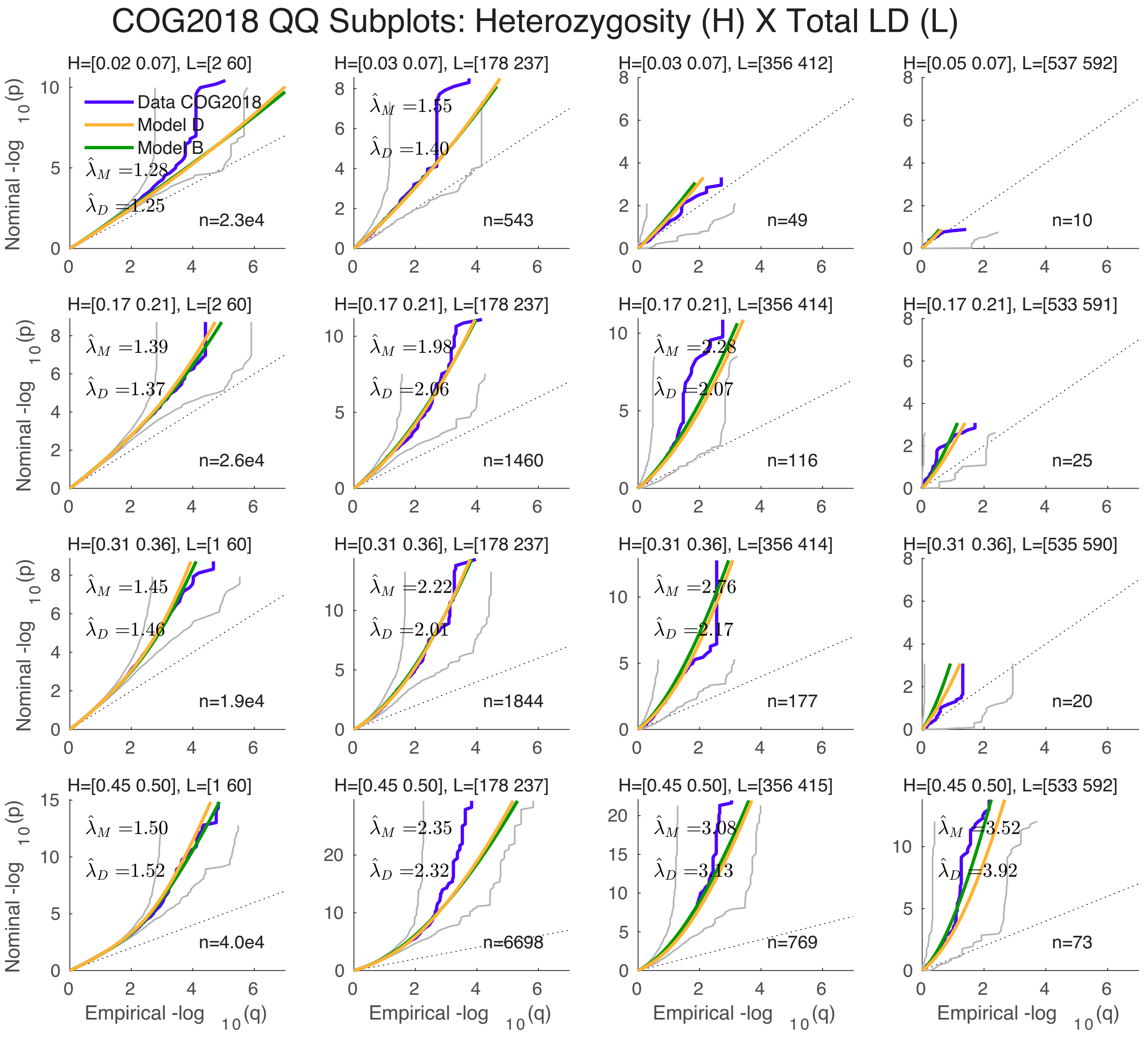


Figure S25: A 4×4 heterozygosity×TLD grid of QQ plots for intelligence. See Figure 3 (B).

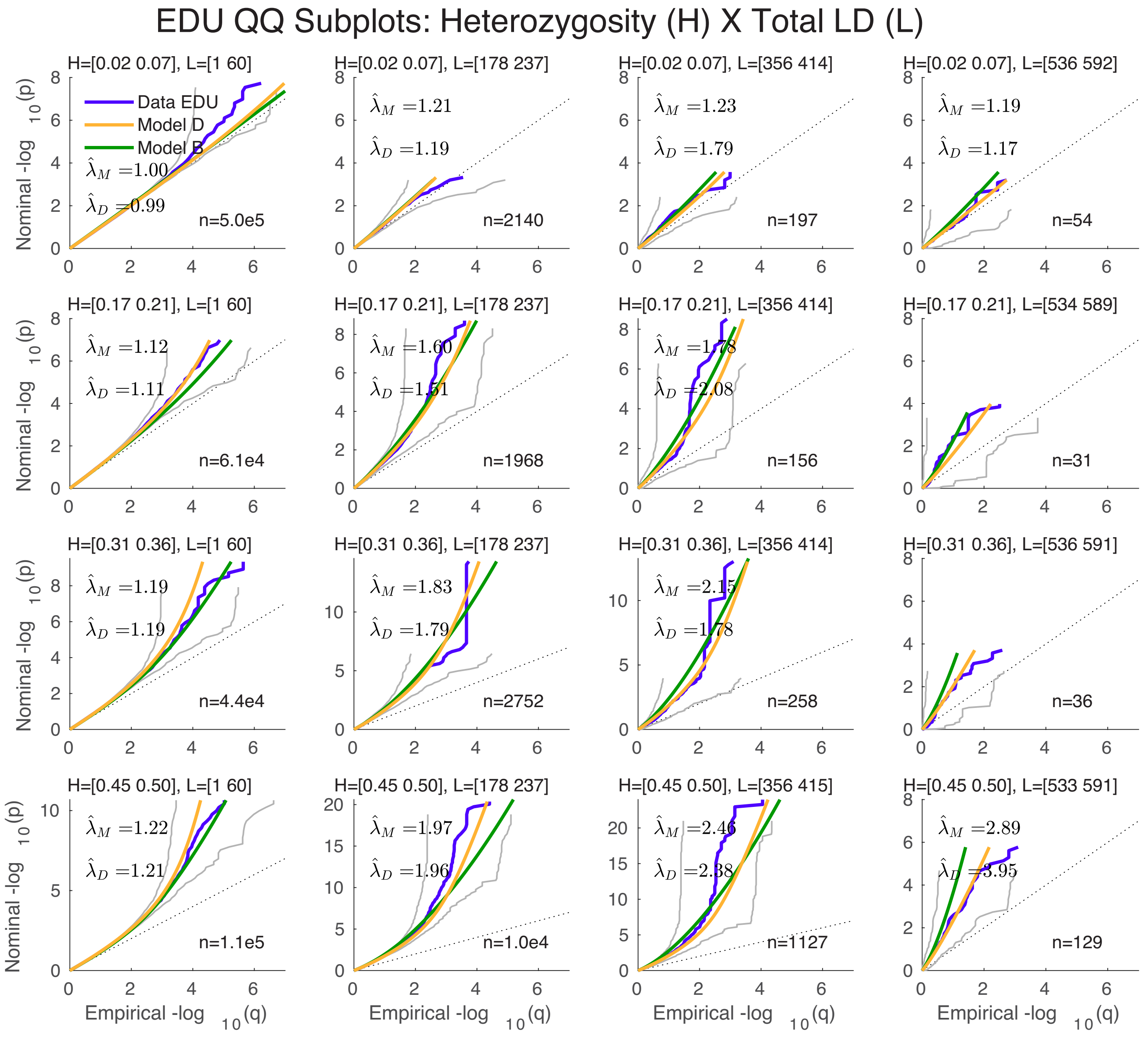


Figure S26: A 4×4 heterozygosity×TLD grid of QQ plots for years of education. See Figure 3 (C).

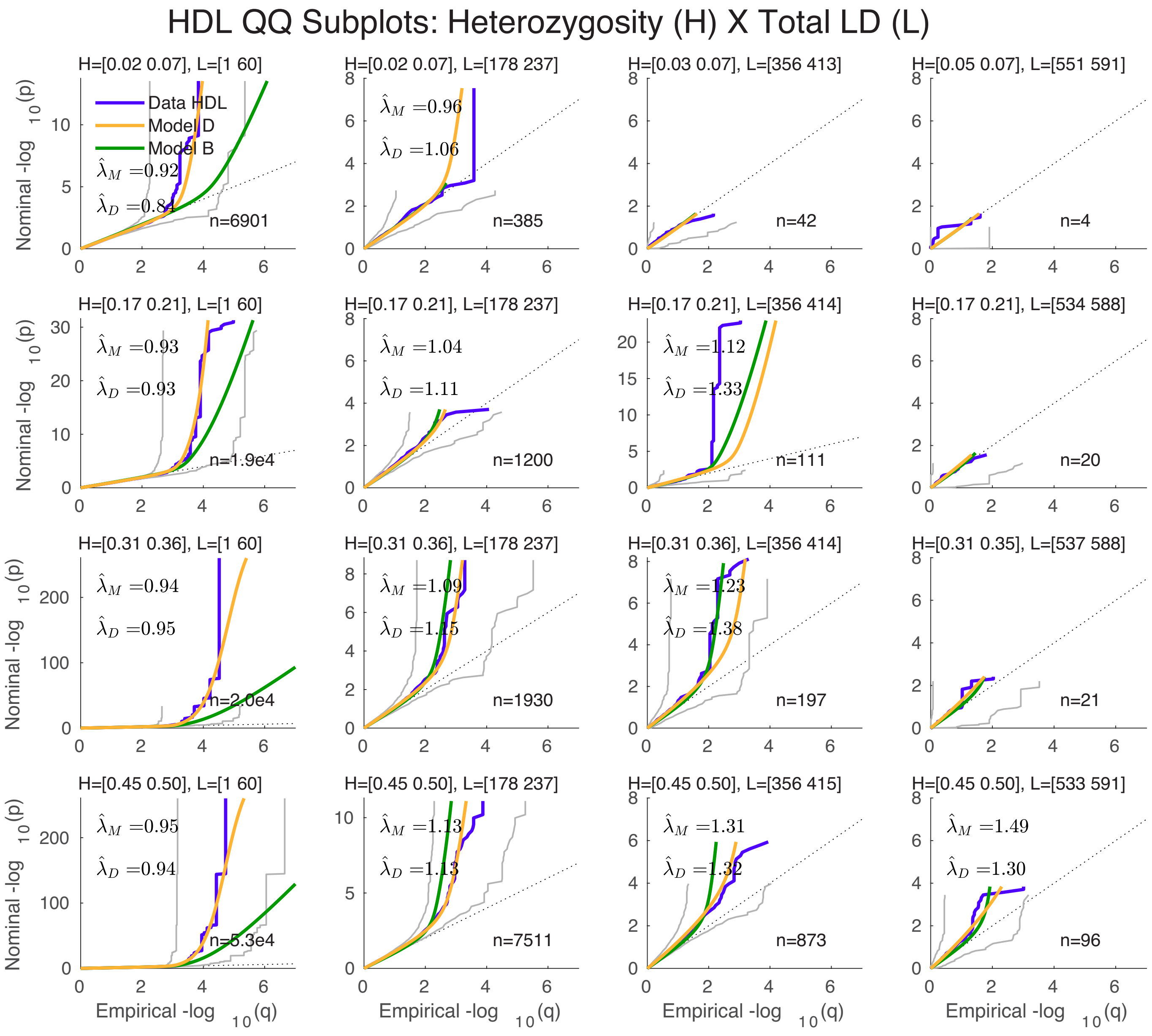


Figure S27: A 4×4 heterozygosity×TLD grid of QQ plots for high density lipoprotein. See Figure 3 (E).

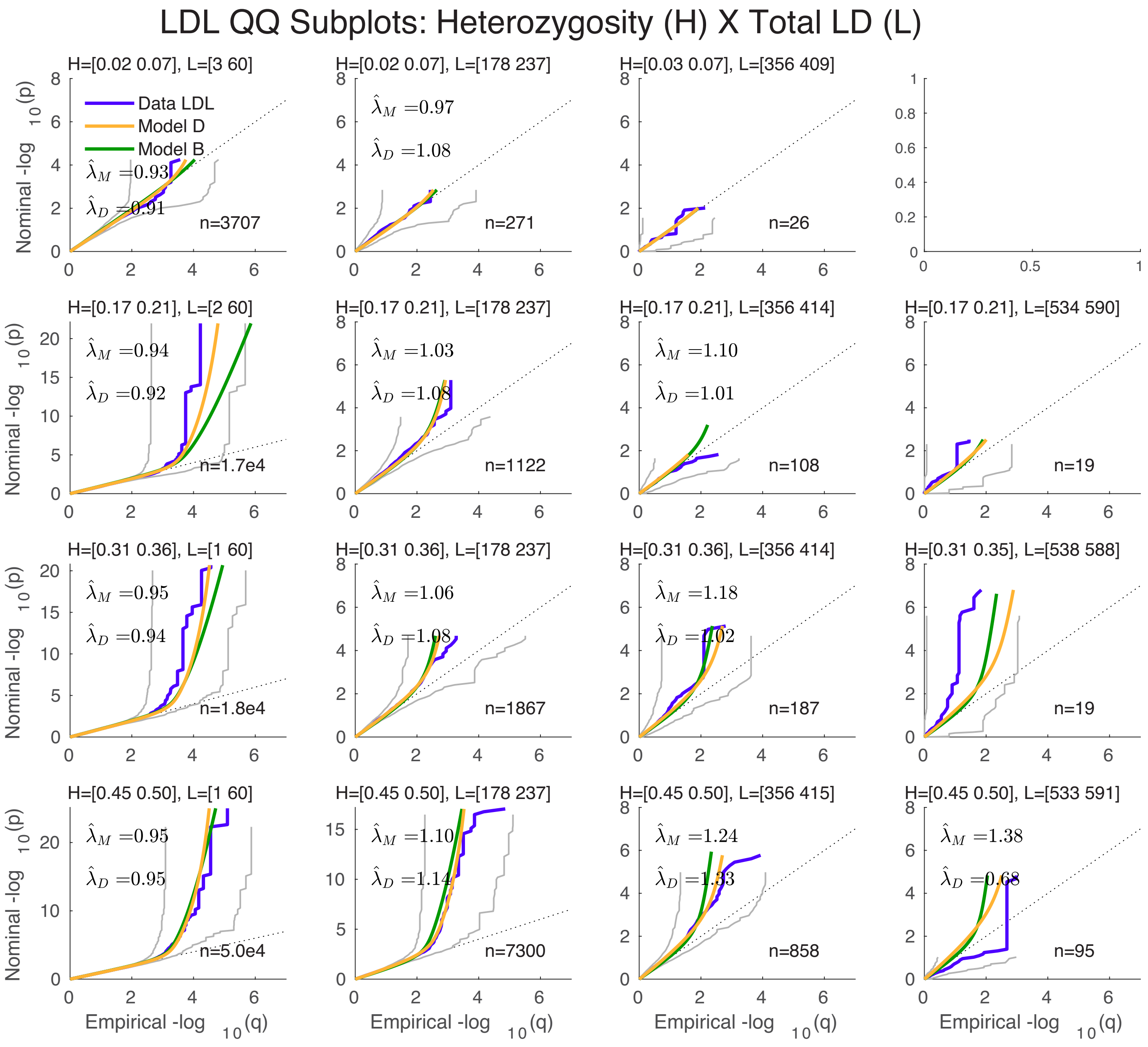


Figure S28: A 4×4 heterozygosity×TLD grid of QQ plots for low density lipoprotein. See Figure 3 (F).

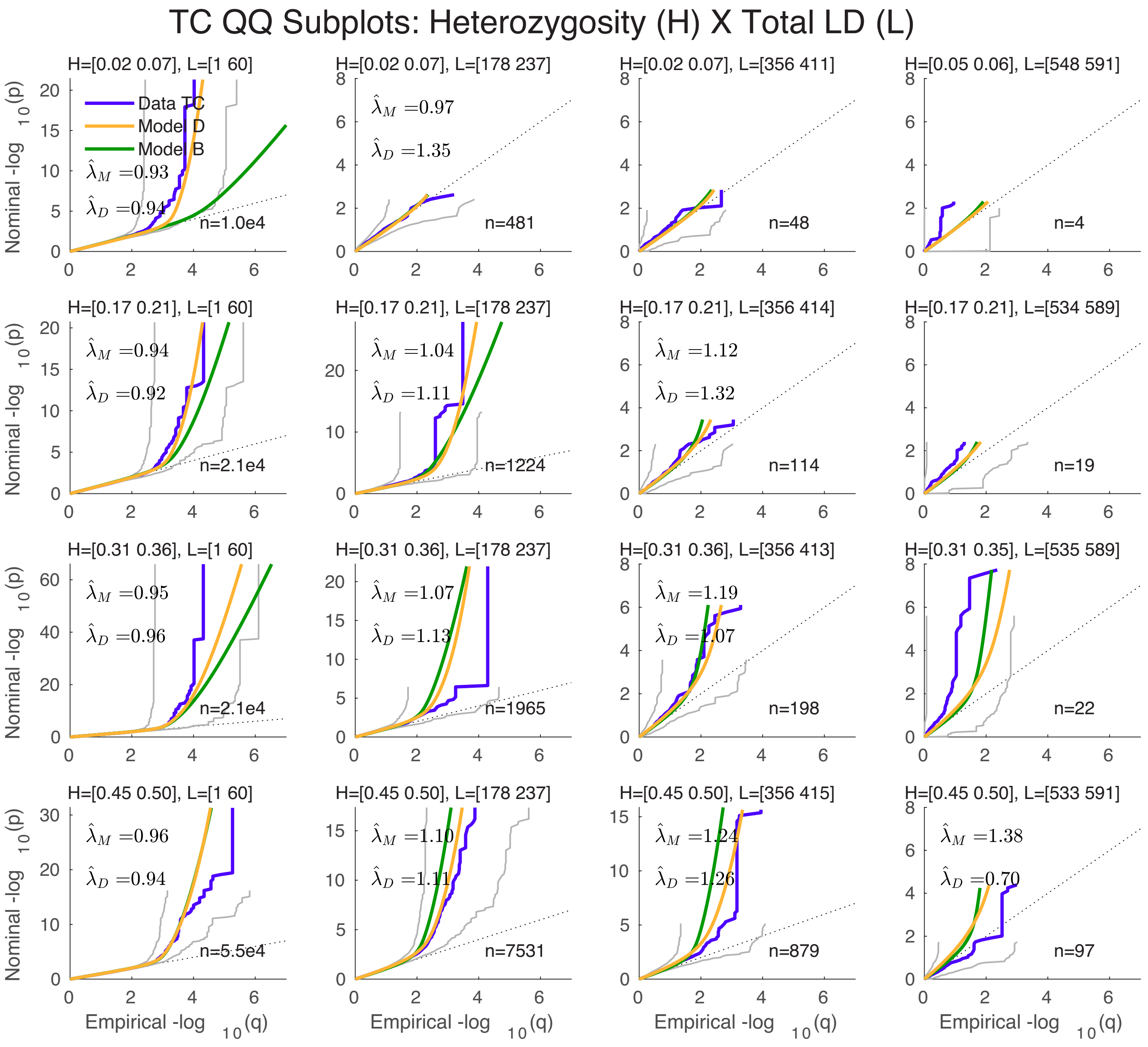


Figure S29: A 4×4 heterozygosity×TLD grid of QQ plots for total cholesterol. See Figure 3 (H).

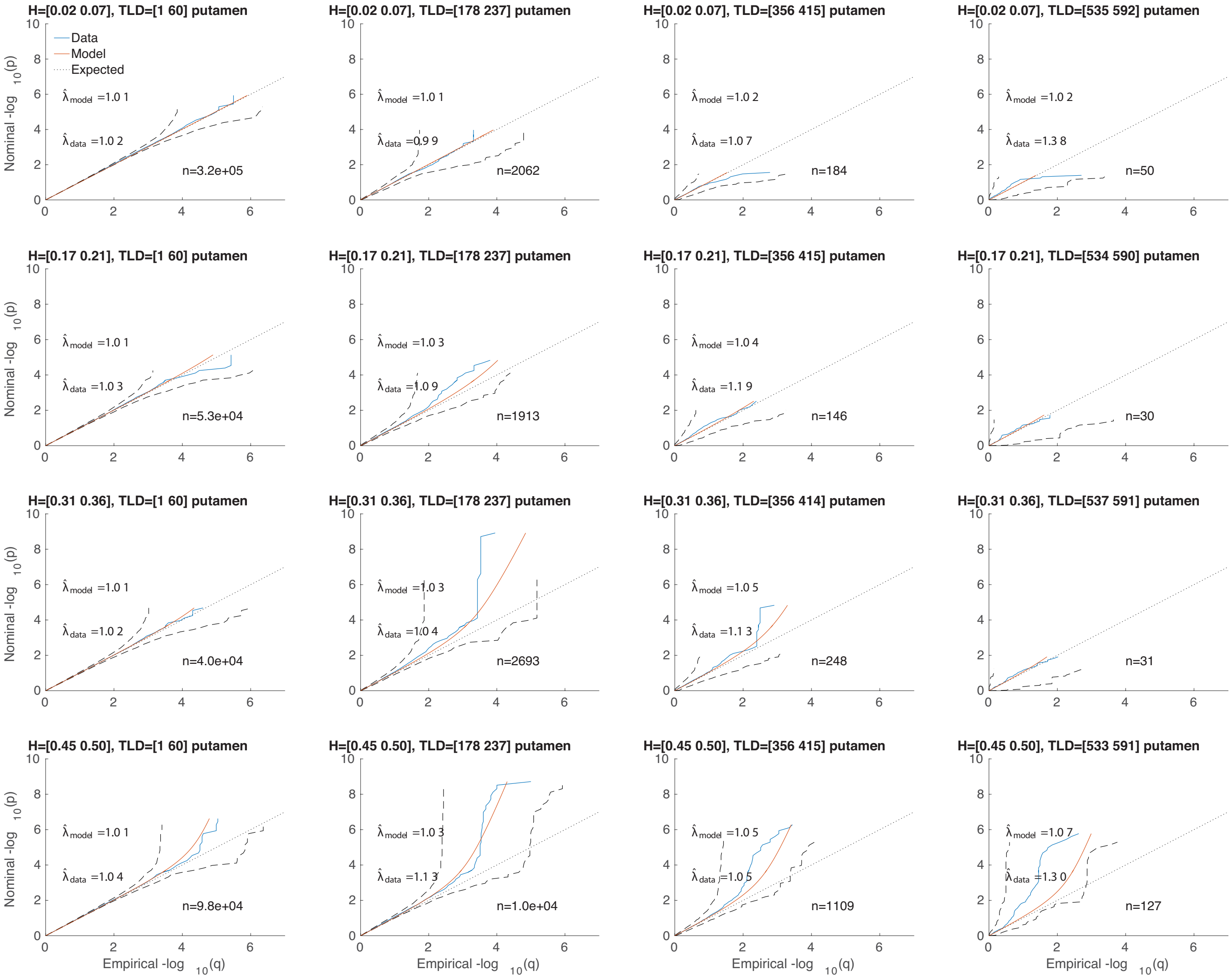


Figure S30: A 4×4 heterozygosity×TLD grid of QQ plots for putamen volume. See Supplemental Material, Figure S10.

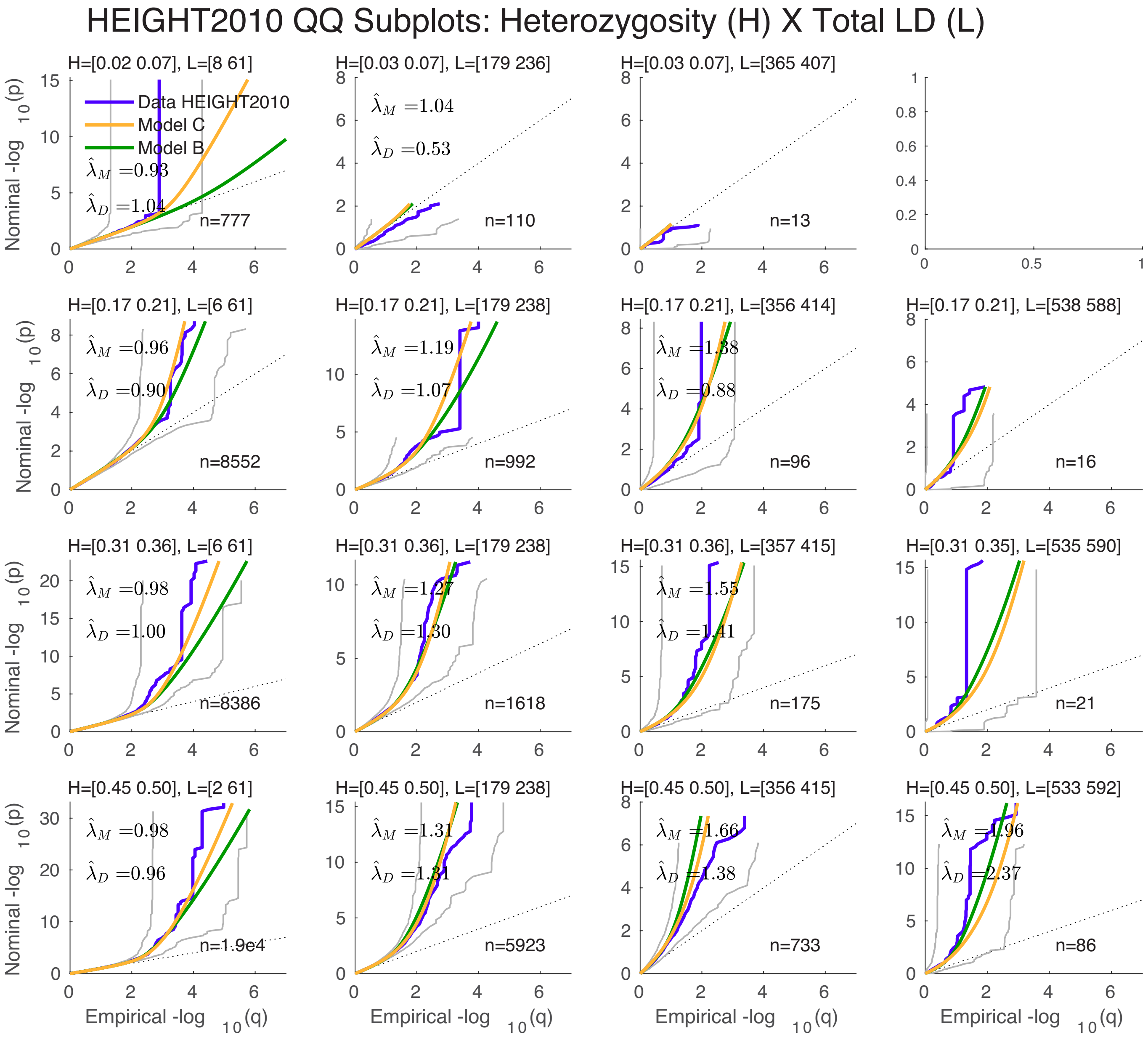


Figure S31: A 4×4 heterozygosity×TLD grid of QQ plots for height (2010). C.f. Figure 3 (G) and Supplemental Material, Figures S32 and S33.

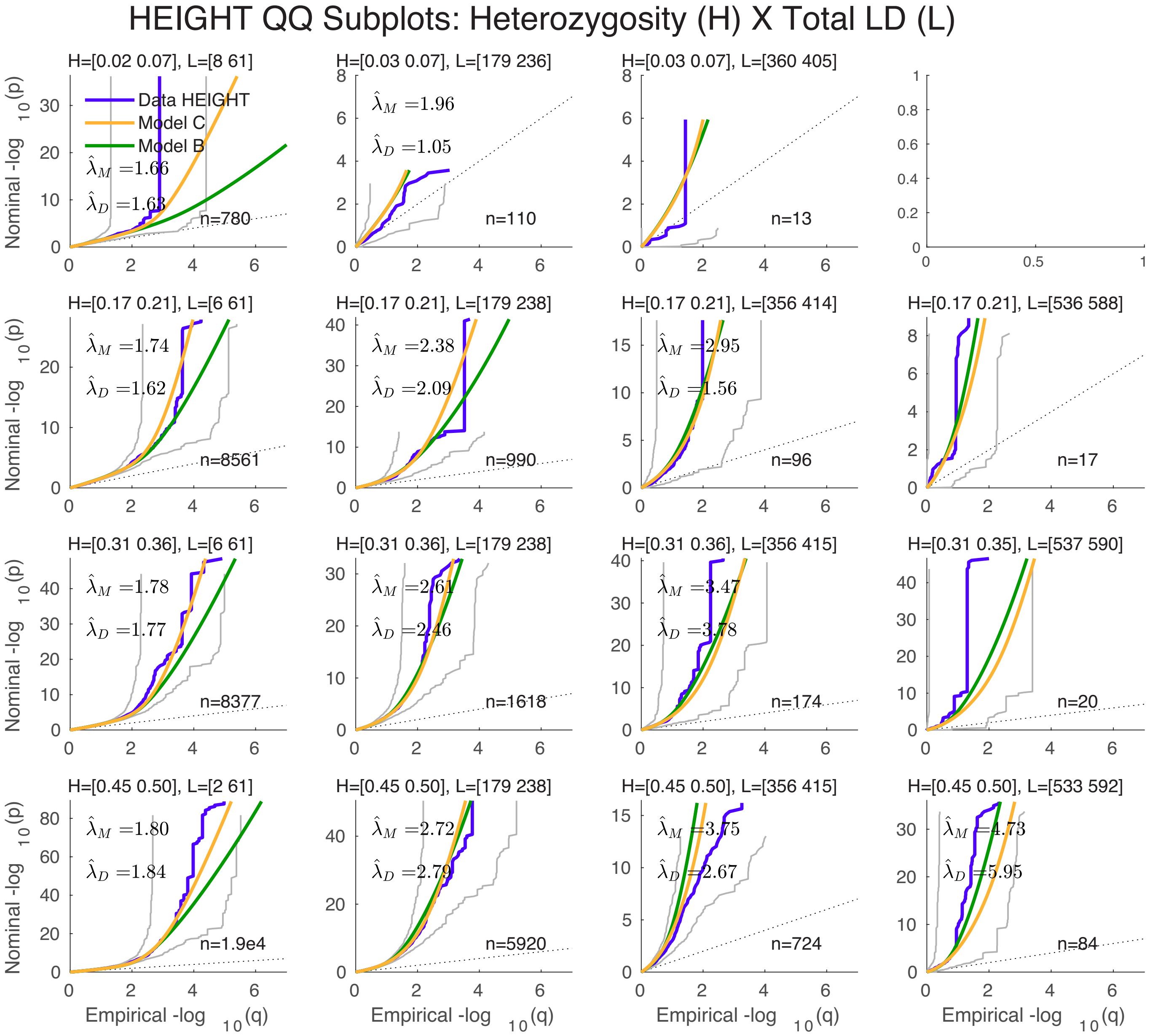


Figure S32: A 4×4 heterozygosity×TLD grid of QQ plots for height (2014). See Figures 3 (G) and Supplemental Material, Figures S31 and S33.

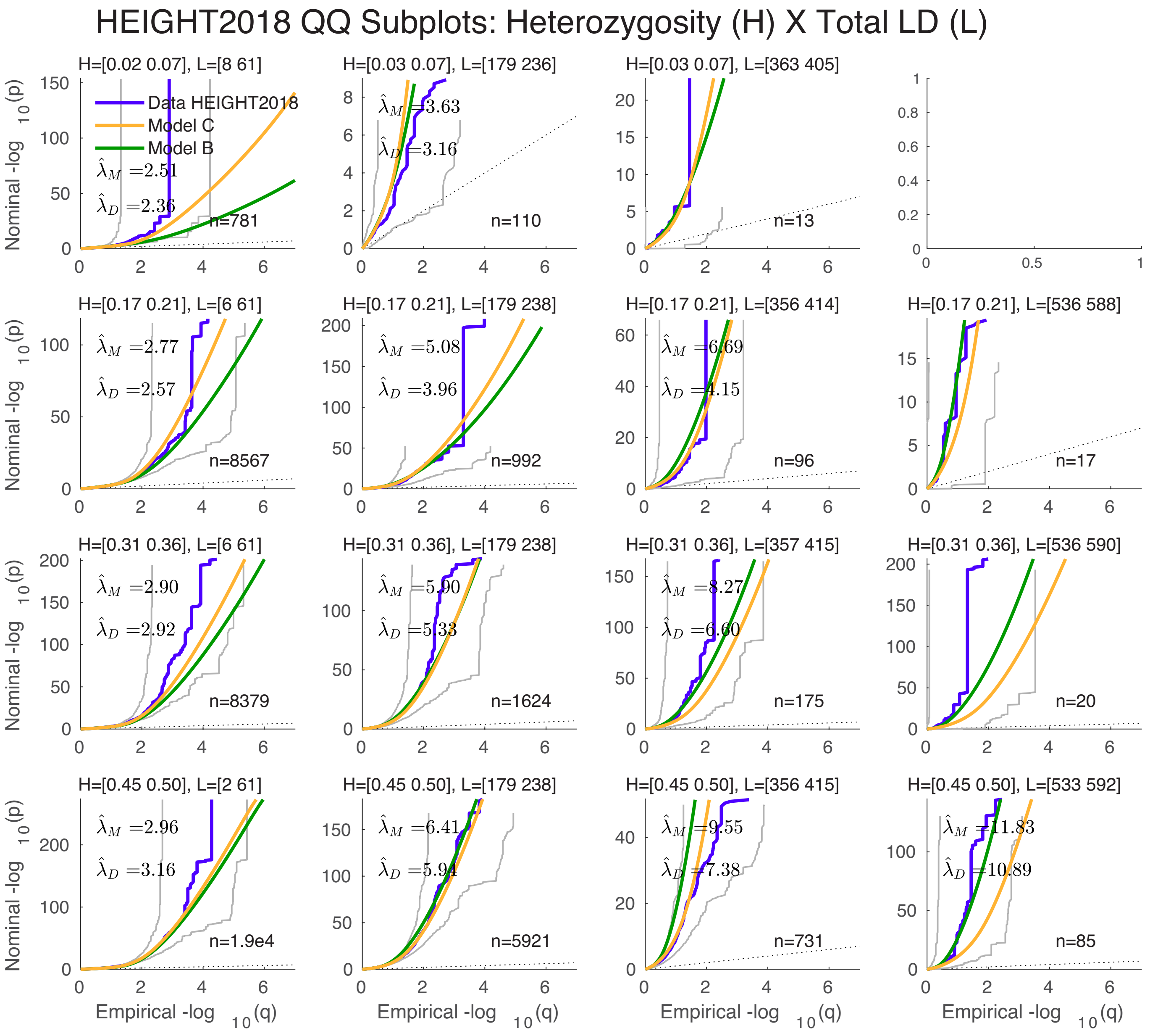


Figure S33: A 4×4 heterozygosity×TLD grid of QQ plots for height (2018 (Yengo et al., 2018)); fit is from model C. C.f. Figure 3 (G) and Supplemental Material, Figures S32 and S33.

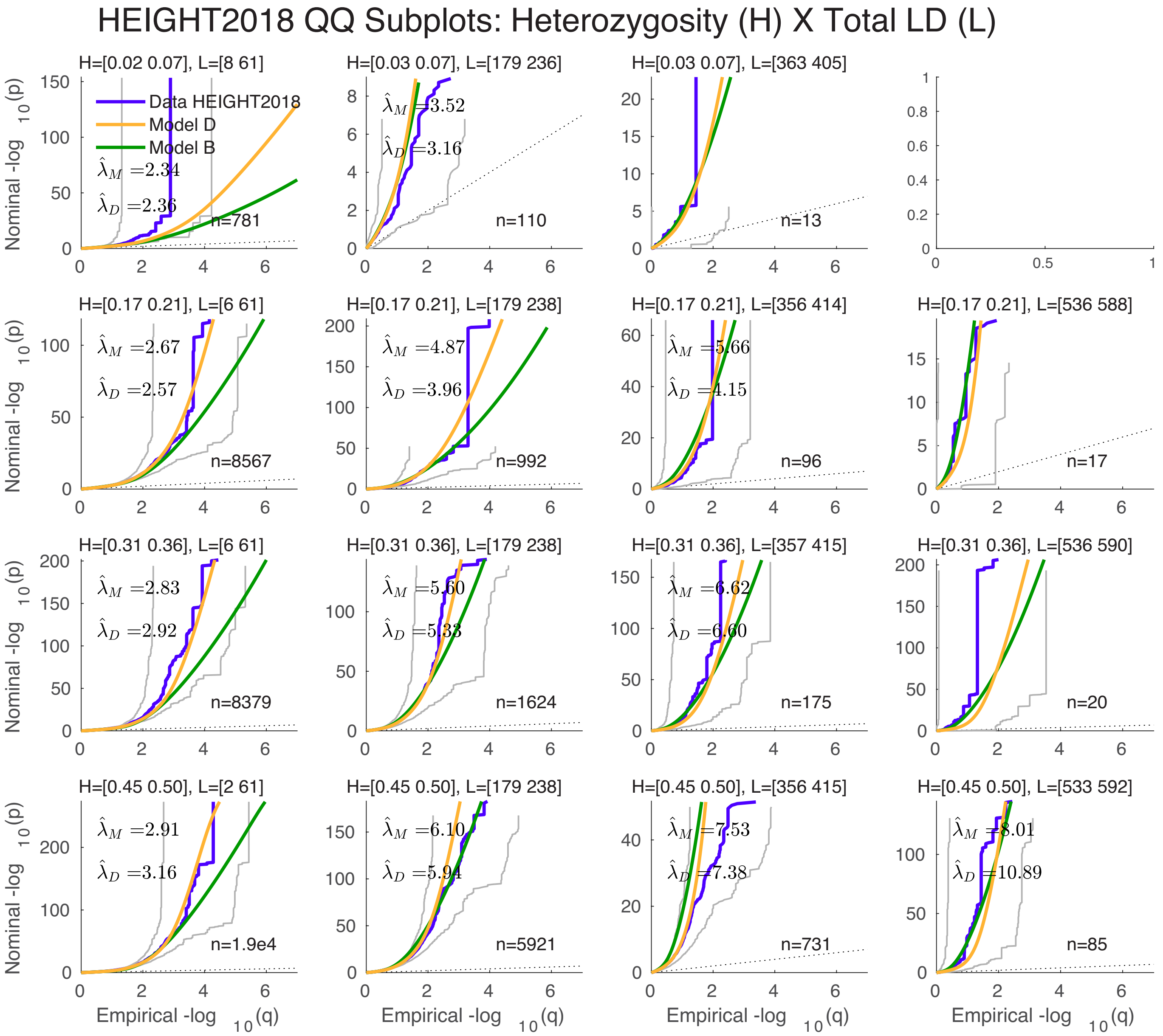


Figure S34: A 4×4 heterozygosity×TLD grid of QQ plots for height (2018 (Yengo et al., 2018)); fit is from model D. See Supplemental Material, Figure S33.

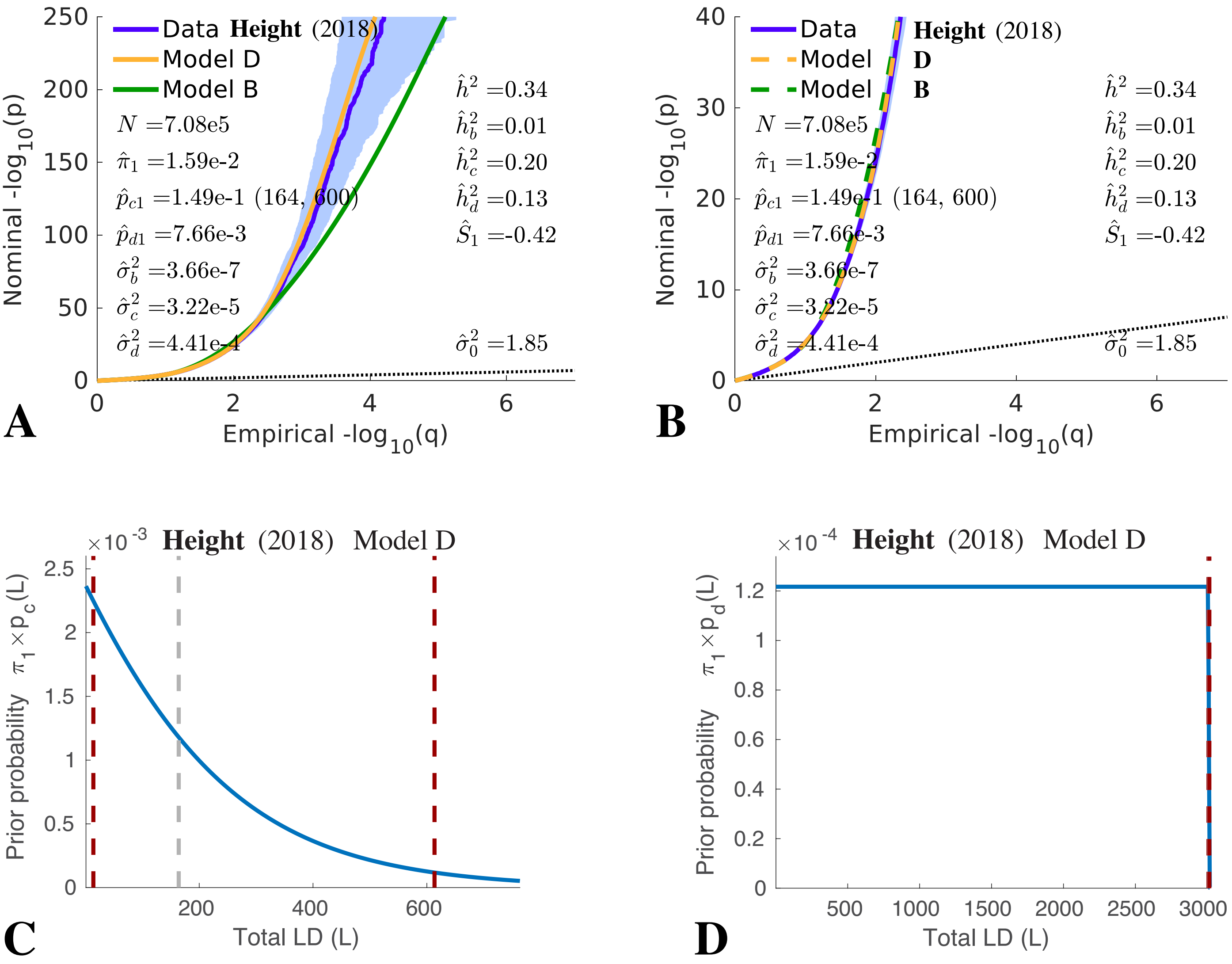


Figure S35: (A) and (B): QQ plot for height (2018 (Yengo et al., 2018)) from model D. (B) is (A) restricted to 40 on the y-axis. See Figure 3 (G). (C) Plot of $\pi_1 \times p_c(L)$ and (D) $\pi_1 \times p_d(L)$, from model D for height (2018 (Yengo et al., 2018)).

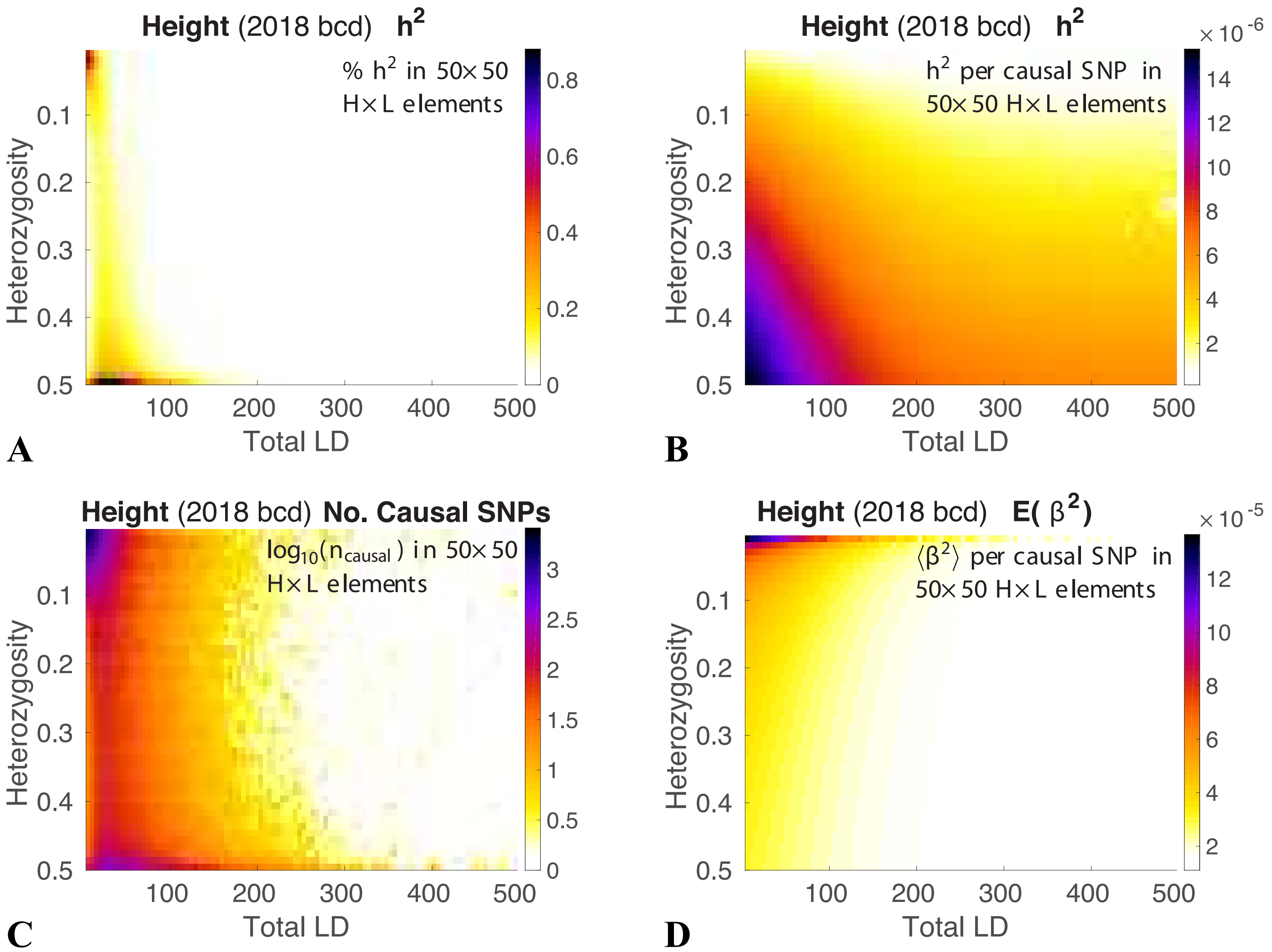


Figure S36: Model results for height (2018), model "D". The reference panel SNPs are binned with respect to both heterozygosity ($H$) and total LD ($L$) in a $50 \times 50$ grid for $0.02 \leqslant H \leqslant 0.5$ and $1 \leqslant L \leqslant 500$. Shown are model estimates of: (A) the percentage of heritability in each grid element; (B) for each element, the average heritability per causal-SNP in the element; (C) $\log_{10}$ of the number of causal SNPs in each element; and (D) the expected $\beta^2$ for the element-wise causal SNPs. Note that $H$ increases from top to bottom.

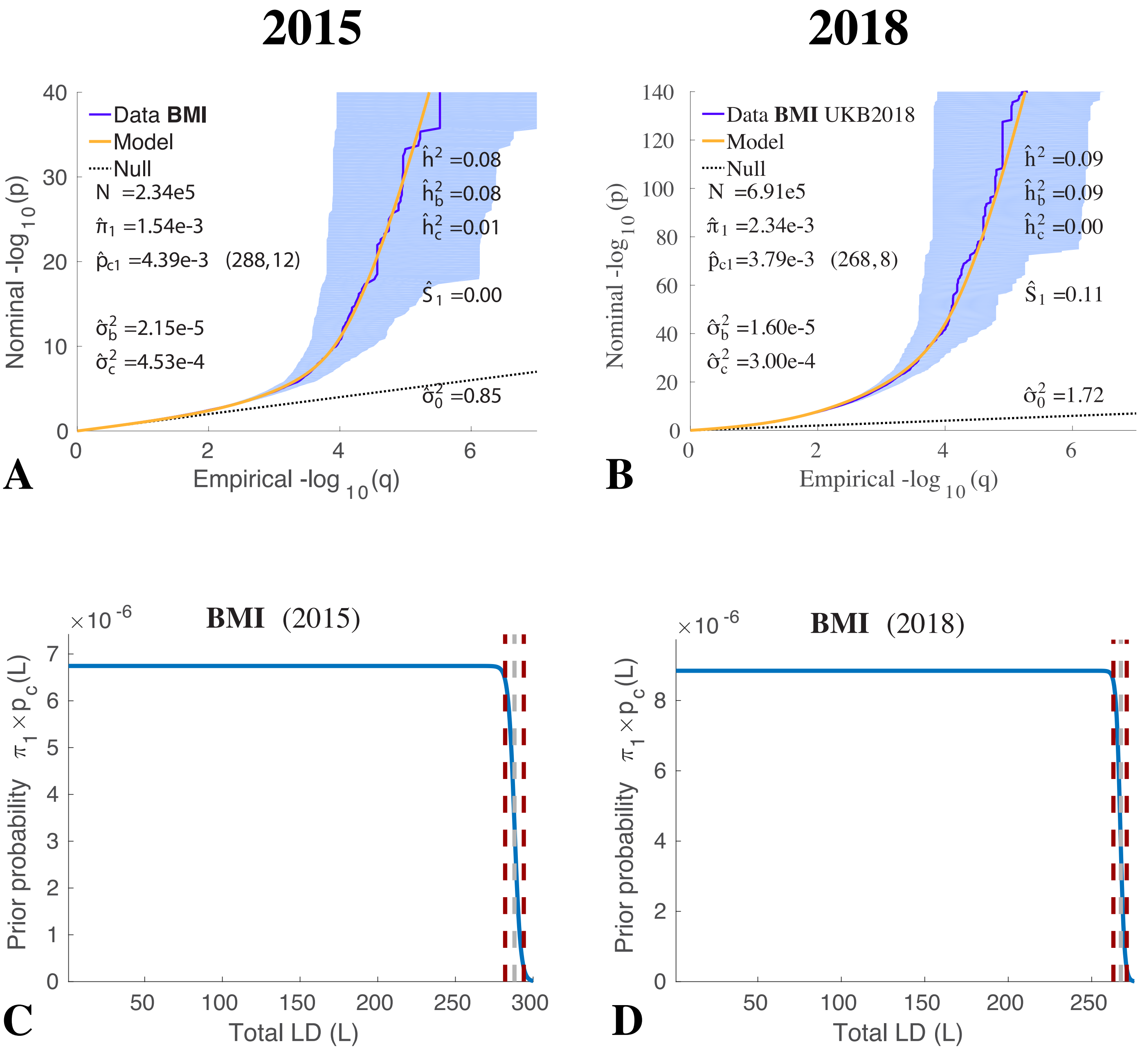


Figure S37: QQ plots for (A) BMI GIANT (2015 (Locke et al., 2015)) and (B) BMI GIANT-UKB (2018 (Yengo et al., 2018)) from model C, showing the consistency of results with very large sample sizes N. See Figure 3 (A), and Supplemental Material, Figures S23 and S24. (C) and (D) are the corresponding plot of $\pi_1 \times p_c(L)$.
